# The Poisson process is the universal law of cancer development: driver mutations accumulate randomly, silently, at constant average rate and for many decades, likely in stem cells

**DOI:** 10.1101/231027

**Authors:** Aleksey V. Belikov, Alexey D. Vyatkin, Sergey V. Leonov

**Affiliations:** Laboratory of Innovative Medicine, School of Biological and Medical Physics, Moscow Institute of Physics and Technology, Institutsky per., 9, Dolgoprudny, Moscow Region, 141701, Russia

**Keywords:** Erlang distribution, gamma distribution, probability distribution, age distribution, cancer incidence, childhood cancer

## Abstract

**Background:** It is assumed that cancers develop upon acquiring a particular number of (epi)mutations in driver genes, but the law governing the kinetics of this process is not known. We have recently shown that the age distribution of incidence for 20 most prevalent cancers of old age is best approximated by the Erlang probability distribution. The Erlang distribution describes the probability of several successive random events occurring by the given time according to the Poisson process, which allows to predict the number of critical driver events.

**Results:** Here we show that the Erlang distribution is the only classical probability distribution that can adequately model the age distribution of incidence for all studied childhood and young adulthood cancers, in addition to cancers of old age.

**Conclusions:** This validates the Poisson process as the universal law describing cancer development at any age and the Erlang distribution as a useful tool to predict the number of driver events for any cancer type. The Poisson process signifies the fundamentally random timing of driver events and their constant average rate. As waiting times for the occurrence of the required number of driver events are counted in decades, it suggests that driver mutations accumulate silently in the longest-living dividing cells in the body - the stem cells.

## Background

Since the discovery of the connection between cancer and mutations in DNA, in the middle of the XX century, there have been multiple attempts to deduce the law of driver mutation accumulation from the age distribution of cancer incidence or mortality ^1^. The proposed models, however, suffer from several serious drawbacks. For example, early models assume that cancer mortality increases with age according to the power law ^2–4^, whilst already at that time it was known that many cancers display deceleration of mortality growth at advanced age. Moreover, when large-scale incidence data have accumulated, it became clear that cancer incidence not only ceases to increase with age but, for at least some cancers, starts to decrease ^5, 6^. More recent models of cancer progression are based on multiple unverified biological assumptions, consist of complicated equations that incorporate many predetermined empirical parameters, and still have not been shown to describe the decrease in cancer incidence at an advanced age ^7–12^. It is also clear that an infinite number of such mechanistic models can be created and custom tailored to fit any set of data, leading us to question their explanatory and predictive values.

Recently, we have proposed that the age distribution of cancer incidence is, in fact, a statistical distribution of probabilities that a required number of driver events occurs precisely by the given age, i.e. a probability density function ^13^. We then tested the probability density functions of 16 well-known continuous probability distributions for fits with the CDC WONDER data on the age distribution of incidence for 20 most prevalent cancers of old age. The best fits were observed for the gamma distribution and its special case - the Erlang distribution, with the average R^2^ of 0.995 ^13^. Notably, these two distributions describe the probability of several independent random events occurring precisely by the given time, according to the Poisson process. This allowed us to estimate the number of driver events, the average time interval between them and the maximal populational susceptibility, for each cancer type. The results showed high heterogeneity in all three parameters amongst the cancer types but high reproducibility between the years of observation ^13^.

However, four other probability distributions - the extreme value (Gumbel), normal, logistic and Weibull - also showed good fits to the data, although inferior to the gamma and Erlang distributions. This leaves some uncertainty regarding the exceptionality of the gamma/Erlang distribution for the description of cancer incidence. Here we test these shortlisted distributions on the CDC WONDER data on childhood and young adulthood cancers. We show that the gamma/Erlang distribution is the only distribution that provides meaningful fits for all tested cancers. This result validates the Poisson process as the fundamental law describing the age distribution of cancer incidence for any cancer type, which also allows to predict important parameters of cancer development, including the number of driver events.

## Methods

### Data acquisition

United States Cancer Statistics Public Information Data: Incidence 1999 - 2012 were downloaded via Centers for Disease Control and Prevention Wide-ranging OnLine Data for Epidemiologic Research (CDC WONDER) online database (http://wonder.cdc.gov/cancer-v2012.HTML). The United States Cancer Statistics (USCS) are the official federal statistics on cancer incidence from registries having high-quality data for 50 states and the District of Columbia. Data are provided by The Centers for Disease Control and Prevention National Program of Cancer Registries (NPCR) and The National Cancer Institute Surveillance, Epidemiology and End Results (SEER) program. Results were grouped by 5-year Age Groups, Crude Rates were selected as an output and All Ages were selected in the Age Group box. All other parameters were left at default settings. Crude Rates are expressed as the number of cases reported each calendar year per 100 000 population. A single person with more than one primary cancer verified by a medical doctor is counted as a case report for each type of primary cancer reported. The population estimates for the denominators of incidence rates are a slight modification of the annual time series of July 1 county population estimates (by age, sex, race, and Hispanic origin) aggregated to the state or metropolitan area level and produced by the Population Estimates Program of the U.S. Bureau of the Census (Census Bureau) with support from the National Cancer Institute (NCI) through an interagency agreement. These estimates are considered to reflect the average population of a defined geographic area for a calendar year. The data were downloaded separately for each specific cancer type, upon its selection in the Childhood Cancers tab. Only cancers that show childhood/young adulthood incidence peaks and do not show middle/old age incidence peaks were analysed further. The middle age of each age group was used as the x value, e.g. 17.5 for the “15-19 years” age group.

### Data analysis

For data analysis, custom scripts were written in Python 3.7.: https://github.com/belikov-av/childhoodcancers The following packages were used: Scikit-learn 0.22.1, Numpy 1.18.1, Scipy 1.4.1, Pandas 1.0.3, Plotly 4.5.4 and Matplotlib 3.1.3.

Distributions were defined as follows:

~~~
def Erlang_pdf(k, b, t):
 return (t**(k-1) * np.exp(-t/float(b))) / (b**k * gamma(k))
def Weibull_pdf(k, b, t):
 return (k/b) * (t/b)**(k-1) * np.exp(-(t/b)**k)
def Extreme_value_pdf(mu, b, t):
 return np.exp((mu - t) / b) * (1/b) * np.exp(-np.exp((mu - t) / b))
def Logistic_pdf(mu, b, t):
 return (1/b) * np.exp((t - mu) / b) / np.square(1 + np.exp((t - mu) / b))
def Normal_pdf(mu, b, t):
 return (1/(b * np.sqrt(2 * pi))) * np.exp(-0.5 * np.square((t - mu)/b))
~~~

In order to find the optimal parameters for every distribution, a grid search method was used with a uniform grid, which consisted of 400 points along each of 2 dimensions. In fact, each distribution has two parameters plus an amplitude factor, thus a grid search is computationally feasible. As the scaled probability density function is a linear function with respect to the amplitude factor, the metric function is convex with respect to it. Thus, the optimal amplitude for a given distribution and its parameters could be found via zero-order optimization methods. In present work, the golden section method was utilized. Given a set of real data points, the area under the curve (which is actually the amplitude) was estimated as the sum of all values. The left boundary was defined as an estimated amplitude divided by 100 and the right boundary was defined as an estimated amplitude multiplied by 100. For each pair of parameters, after the optimal amplitude was found via the golden section method, the distribution values were calculated in those points, for which the real data were presented, and the goodness of fit (R^2^ score) was calculated using predicted and real values.

## Results

To test the universality and exceptionality of the gamma/Erlang distribution, the publicly available USA incidence data on childhood and young adulthood cancers were downloaded from the CDC WONDER database (see Materials and Methods). In addition to the gamma/Erlang distribution, the probability density functions of the following continuous probability distributions were selected for testing based on their good fits to adult cancers^13^: extreme value (Gumbel), logistic, normal and Weibull. We used a custom “grid search” script to explore all combinations of parameters and find true (non-local) R^2^ maxima (see Figure 1 for gamma/Erlang distribution, Supplementary figures for other distributions and Materials and methods for the detailed description). Results showed that the extreme value (Gumbel), logistic and normal distributions fit to some childhood cancers in a way that a large part of their probability density function appears in the negative range of x values (Figure 2). As the x axis corresponds to patients’ ages, this makes these three distributions uninterpretable in the biological context. Hence, they were excluded from the further analysis. The remaining gamma/Erlang and Weibull distributions are defined only for positive x values, so there was no such problem (Figure 2). The gamma/Erlang distribution provided closer fits to the childhood and young adulthood cancers than the Weibull distribution (Figure 3, Table 1).

**Fig. 1.**
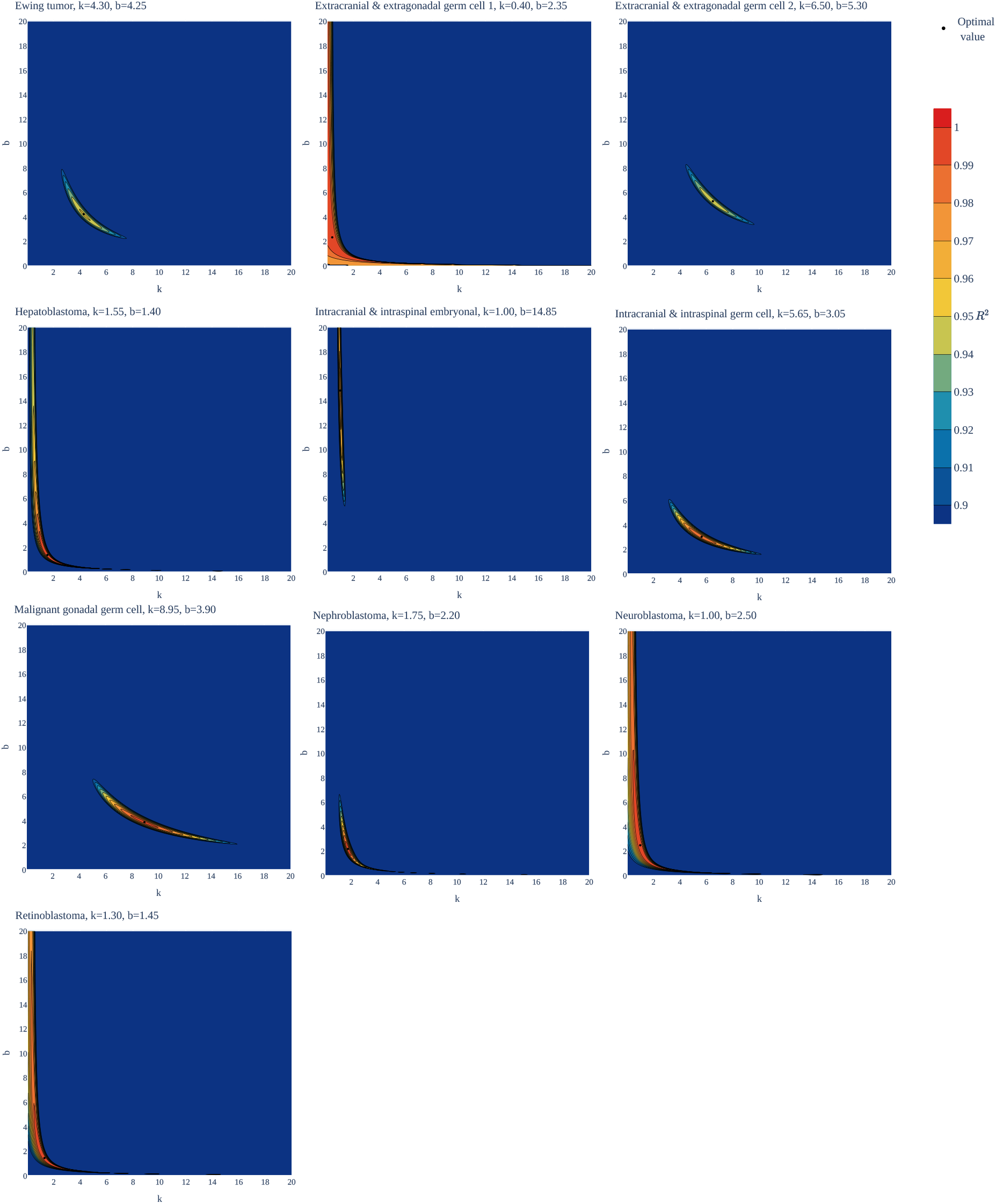
Goodness of fit of the gamma/Erlang distribution to 10 childhood/young adulthood cancer types as a function of various parameter combinations. The ***k*** and ***b*** parameters were sampled with 0.05 interval, and for each pair the amplitude (***A***, not shown) and goodness of fit (R^2^) were calculated. See Supplementary figures for the other distributions.

**Fig. 2.**
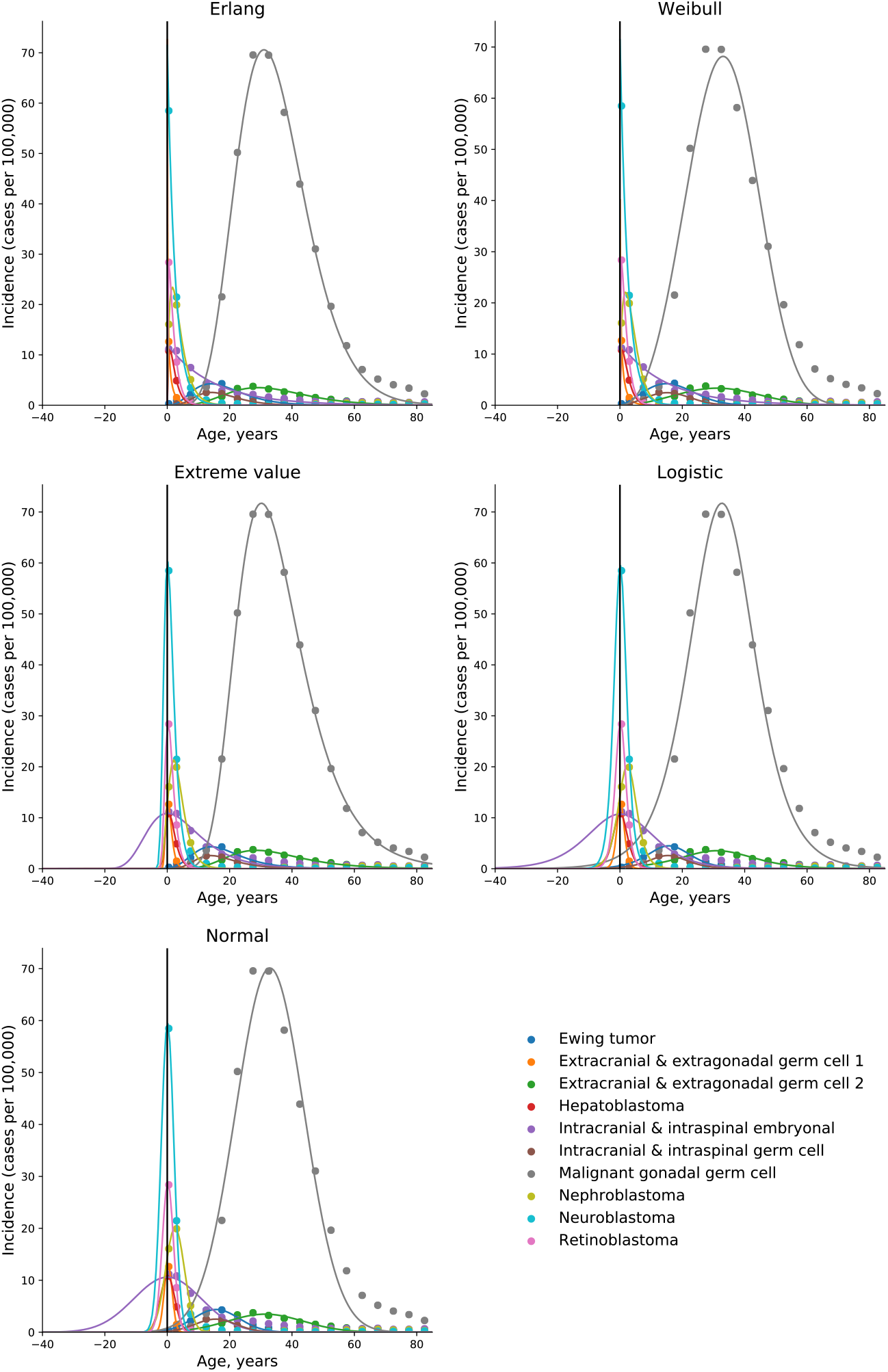
Only the gamma/Erlang and the Weibull distributions fit the actual age distributions of childhood and young adulthood cancer incidence without requiring negative age values. Dots indicate crude incidence rates for 5-year age groups, curves indicate probability density functions fitted to the incidence data for various childhood/young adulthood cancer types. The middle age of each age group is plotted. See Supplementary figures for the optimal parameter values.

**Fig. 3.**
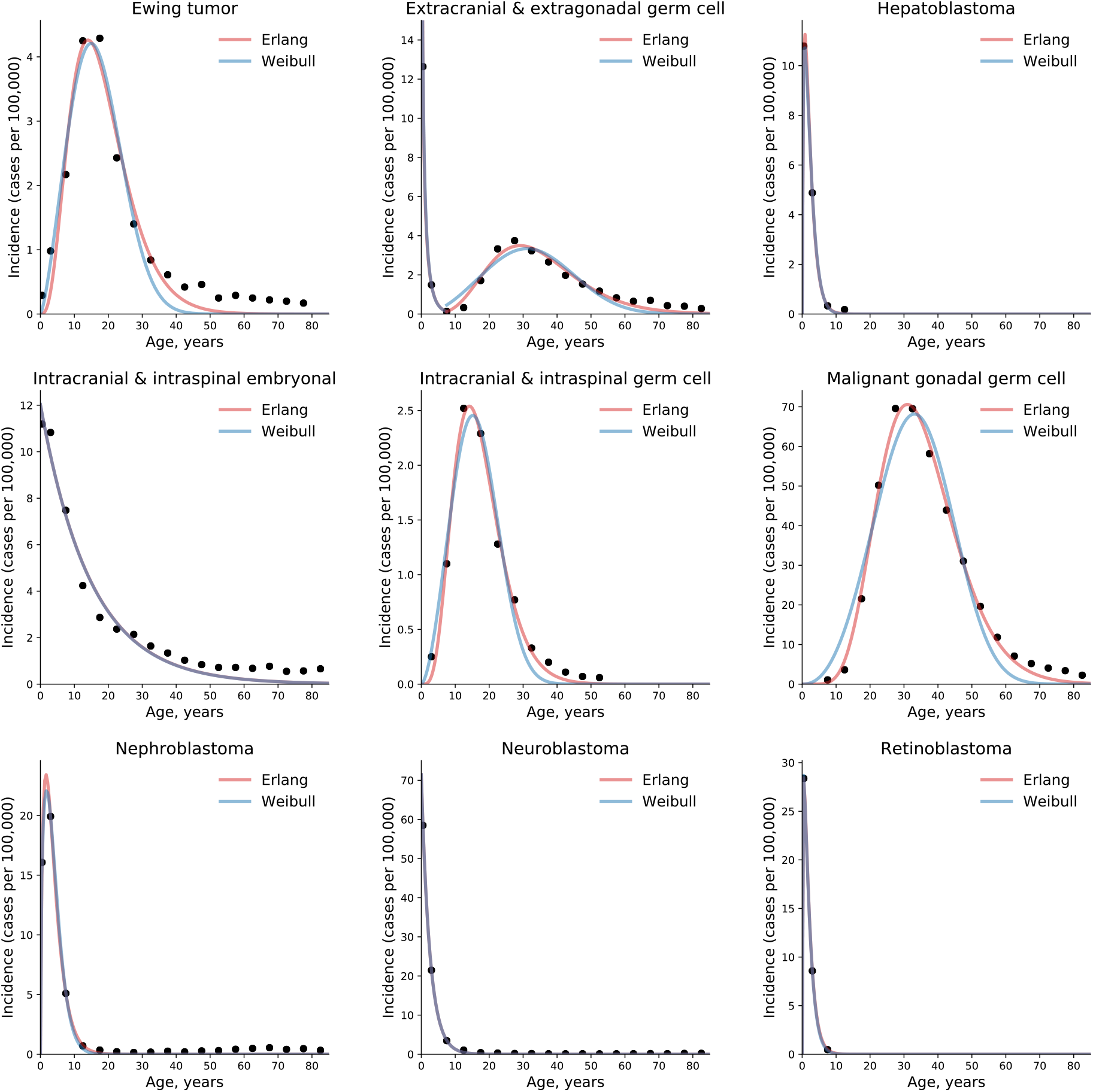
The gamma/Erlang distribution approximates the age distribution of incidence for childhood and young adulthood cancers better than the Weibull distribution. Dots indicate crude incidence rates for 5-year age groups, curves indicate the probability density function of the gamma/Erlang (red) or Weibull (blue) distribution fitted to the incidence data (see Table 1 for R^2^ comparison and Table 2 for estimated parameters). The middle age of each age group is plotted. Extracranial and extragonadal germ cell tumours of childhood and young adulthood are shown on the same plot.

**Table 1.** Comparison of the goodness of fit (R^2^) of the gamma/Erlang and Weibull distributions to the actual age distributions of childhood/young adulthood cancer incidence.

| **Cancer type** | **gamma/**<br>**Erlang** | **Weibull** |
| --- | --- | --- |
| Ewing tumour and related sarcomas of bone | **0.9516** | 0.9476 |
| Extracranial and extragonadal germ cell tumours of childhood | **1.000** | **1.000** |
| Extracranial and extragonadal germ cell tumours of young adulthood | **0.9495** | 0.8689 |
| Hepatoblastoma | **0.9996** | **0.9996** |
| Intracranial and intraspinal embryonal tumours | **0.9738** | **0.9738** |
| Intracranial and intraspinal germ cell tumours | **0.9890** | 0.9674 |
| Malignant gonadal germ cell tumours | **0.9941** | 0.9530 |
| Nephroblastoma and other nonepithelial renal tumours | **0.9970** | 0.9969 |
| Neuroblastoma and ganglioneuroblastoma | **0.9998** | **0.9998** |
| Retinoblastoma | **1.000** | **1.000** |
| **Average** | **0.9854** | 0.9707 |

The best fit for each cancer type is highlighted in bold. See Figure 3 for graphical representation.

Importantly, the gamma distribution and the Erlang distribution derived from it are the only classical continuous probability distributions that describe the cumulative waiting time for *k* successive random events, with the Erlang distribution differing only in counting events as integer numbers. Because these properties suit excellently to describe the waiting time for real discrete random events such as driver mutations, the gamma/Erlang distribution provides the opportunity to get unique insights into the carcinogenesis process. We have previously proposed that the shape parameter *k* of the gamma/Erlang distribution indicates the average number of driver events that need to occur in order for a cancer to develop to a stage that can be detected during clinical screening; the scale parameter *b* indicates the average time interval (in years) between such events; and the amplitude parameter *A* divided by 1000 estimates the maximal susceptibility (in percent) of a given population to a given type of cancer ^13^.

To obtain these parameter values, the gamma/Erlang distribution was fitted individually to incidence of each of 10 childhood/young adulthood cancer types (Figure 3, Table 2). The non-integer values of the shape parameter *k* can be easily explained if we suppose that the studied population consists of the mixture of patients with slightly different numbers of driver events. The goodness of fit varied from 0.9495, for extracranial and extragonadal germ cell tumours of young adulthood, to 1.000, for extracranial and extragonadal germ cell tumours of childhood and retinoblastoma, with the average of 0.9854. The predicted number of driver events varied from 0.4, for extracranial and extragonadal germ cell tumours of childhood, to 8.95, for malignant gonadal germ cell tumours. The predicted average time between the events varied from 1.4 years, for hepatoblastoma, to 14.85 years, for intracranial and intraspinal embryonal tumours. The predicted maximal populational susceptibility varied from 0.0322%, for extracranial and extragonadal germ cell tumours of childhood, to 1.966%, for malignant gonadal germ cell tumours. Overall, the data confirm high heterogeneity in carcinogenesis patterns revealed in the previous study ^13^.

**Table 2.** Estimated carcinogenesis parameters for 10 childhood/young adulthood cancer types.

| **Cancer type** | ***k*** | ***b*** | ***A/1000*** |
| --- | --- | --- | --- |
|  | **Number of driver events** | **Average time between events, years** | **Maximal populational susceptibility, %** |
| Ewing tumour and related sarcomas of bone | 4.3 | 4.25 | 0.0846 |
| Extracranial and extragonadal germ cell tumours of childhood | 0.4 | 2.35 | 0.0322 |
| Extracranial and extragonadal germ cell tumours of young adulthood | 6.5 | 5.3 | 0.111 |
| Hepatoblastoma | 1.55 | 1.4 | 0.0338 |
| Intracranial and intraspinal embryonal tumour | 1.0 | 14.85 | 0.179 |
| Intracranial and intraspinal germ cell tumours | 5.65 | 3.05 | 0.0426 |
| Malignant gonadal germ cell tumours | 8.95 | 3.9 | 1.966 |
| Nephroblastoma and other nonepithelial renal tumours | 1.75 | 2.2 | 0.124 |
| Neuroblastoma and ganglioneuroblastoma | 1.0 | 2.5 | 0.179 |
| Retinoblastoma | 1.3 | 1.45 | 0.072 |

The parameters are determined for the gamma/Erlang distribution fitted to actual cancer incidence data (see Figure 3).

## Discussion

We have previously shown that five probability distributions - the extreme value (Gumbel), gamma/Erlang, normal, logistic and Weibull - approximate the age distribution of incidence for 20 most prevalent cancers of old age ^13^. The shape of those incidence distributions resembles the bell shape of the normal distribution, with some asymmetry, or at least the left part of it. However, many cancers of childhood have a radically different shape of the incidence distribution, the shape of the exponential distribution (Figure 3). Of the five shortlisted distributions, only the gamma/Erlang and Weibull distributions can assume that shape, i.e. reduce to the exponential distribution when the parameter *k* equals one or less. Other distributions can fit the data only by extending their probability density function into the negative range of age values (Figure 2), which makes their proper biological interpretation highly unlikely. Of the remaining two distributions, the gamma/Erlang provides superior fit compared to Weibull. In fact, for cancers of old age, the average R^2^ for the Weibull distribution is 0.9938, whereas for the gamma/Erlang distribution is 0.9954 ^13^. For cancers of childhood and young adulthood, the average R^2^for the Weibull distribution is 0.9707, whereas for the gamma/Erlang distribution is 0.9854 (Table 1). Thus, it appears that the gamma/Erlang distribution is the only classical probability distribution that fits universally well to cancers of childhood, young adulthood and old age.

We have proposed that the parameter *k* of the Erlang distribution indicates the average number of driver events that need to occur in order for a cancer to develop to a stage that can be detected during clinical screening ^13^. As mentioned above, the Erlang distribution reduces to the exponential distribution when *k* equals one, because the exponential distribution describes the waiting time for a single random event. It would thus mean that cancers with the exponential shape of the age distribution of incidence require only a single driver event with random time of occurrence, most likely a somatic driver mutation ^14^ or epimutation ^15^. This explains their maximal prevalence in the early childhood.

In his seminal paper ^16^, Alfred Knudson has proposed that hereditary retinoblastoma, a childhood cancer with the exponential age distribution of incidence, is caused by a single somatic mutation in addition to one heritable mutation. He also proposed that in the nonhereditary form of the disease, both mutations should occur in somatic cells. As hereditary form is estimated to represent about 45% of all cases ^16, 17^, the number of driver mutations predicted from combined incidence data should be around 1.55. Interestingly, whilst the gamma/Erlang distribution fits the incidence data excellently, with R^2^=1.0, it predicts 1.3 driver events (Table 2). This yields the estimate of the hereditary form prevalence at 70%. This higher value may point to the general underestimation of the hereditary component in unilateral retinoblastoma, perhaps due to limitations of routine genetic screening and the influence of genetic mosaicism ^18^. In contrast to retinoblastoma, the hereditary form of neuroblastoma is estimated to comprise only 1-2% of all cases ^19^, hence the exponential age distribution of incidence would mean that only one somatic mutation is required. Indeed, the gamma/Erlang distribution predicts one driver event (Table 2).

The prediction of a single driver event in cancers with the exponential age distribution of incidence does not mean that only a single driver gene can be discovered in such cancer types. In fact, many driver genes are redundant or even mutually exclusive, e.g. when the corresponding proteins are components of the same signalling pathway ^20^. Thus, each tumour in such cancer types is expected to have a mutation in one driver gene out of a set of several possible ones, in which all genes most likely encode members of the same pathway or are responsible for the same cellular function. For example, in each neuroblastoma tumour sample, a mutation was present in only one out of five putative driver genes - *ALK, ATRX, PTPN11, MYCN* or *NRAS*^21^.

Another aspect to consider is that while one mutation is usually sufficient to activate an oncogene, two mutations are typically required to inactivate both alleles of a tumour suppressor gene. Therefore, cancers with the exponential age distribution of incidence are predicted to have either a single somatic mutation in an oncogene, or a single somatic mutation in a tumour suppressor gene plus an inherited mutation in the same gene. The former is the case for neuroblastoma, where an amplification or an activating point mutation in *ALK* is often present ^22–24^. The latter is the case for retinoblastoma, where an inactivating mutation in one allele of *RB1* is usually inherited, whereas an inactivating mutation in the other *RB1* allele occurs in a somatic cell ^25^.

Finally, the number of driver events predicted by the Erlang distribution refers exclusively to rate-limiting events responsible for cancer progression. For example, it was shown that inactivation of both alleles of *RB1* leads first to retinoma, a benign tumour with genomic instability that easily transforms to retinoblastoma upon acquiring additional mutations ^26^. In this case, two mutations in *RB1* are rate-limiting, whereas mutations in other genes are not, because genomic instability allows them to occur very quickly. In neuroblastoma, frequent *MYCN* amplification and chromosome 17q gain are found only in advanced stages of the disease ^27, 28^, so they are unlikely to be the initiating rate-limiting events.

## Conclusions

Overall, application of the gamma/Erlang distribution to childhood and young adulthood cancers showed its exceptionality amongst other probability distributions. The fact that it can successfully describe the radically different age distributions of incidence for cancers of any age and any type allows to call the underlying Poisson process the universal law of cancer development. The Poisson process signifies the fundamentally random timing of driver events and their constant average rate ^13^. The gamma/Erlang distribution allows to calculate, by multiplying the number of driver events by the average time interval between them, that an average person needs from 73 to 324 years to accumulate the required number of driver alterations, depending on the cancer type ^13^. This finding is consistent with the silent accumulation of driver mutations in stem cells before the terminal clonal expansion ^29–31^, because this is the only type of dividing cells surviving for so long in the body, and mutations require cellular division to be fixed in the DNA. For childhood and young adulthood cancers, these estimates range from one to 35 years (see Table 2), but the mechanism is likely the same. Finally, as the gamma/Erlang distribution allows to predict the number and rate of driver events in any cancer subtype for which the data on the age distribution of incidence are available, it may help to optimize the algorithms for distinguishing between driver and passenger mutations ^32^, leading to the development of more effective targeted therapies.

## Declarations

### Ethics approval and consent to participate

The manuscript does not report any unpublished studies involving human participants, human data or human tissue.

### Consent for publication

The manuscript does not contain any individual person’s data in any form.

### Availability of data and materials

The datasets and code generated and/or analysed during the current study are available at https://github.com/belikov-av/childhoodcancers.

### Competing interests

The authors declare that they have no competing interests.

### Funding

This study received no specific funding.

### Authors’ contributions

AVB: conceptualization, data curation, formal analysis, investigation, methodology, project administration, supervision, validation, visualization, writing - original draft, writing - review and editing.

ADV: formal analysis, investigation, methodology, software, visualization, writing - review and editing.

SVL: project administration, resources, supervision, validation, writing - review and editing.

All authors read and approved the final manuscript.

## Acknowledgements

AVB acknowledges MIPT 5-100 program support for early career researchers.

## Notes

### Competing Interest Statement

The authors have declared no competing interest.

### Summary of Updates

New computational method for the optimal distribution parameter finding was employed

