## Supplementary figures and images for "The Poisson process is the universal law of cancer development: driver mutations accumulate randomly, silently, at constant average rate and for many decades, likely in stem cells"

### Ewing tumor.pdf

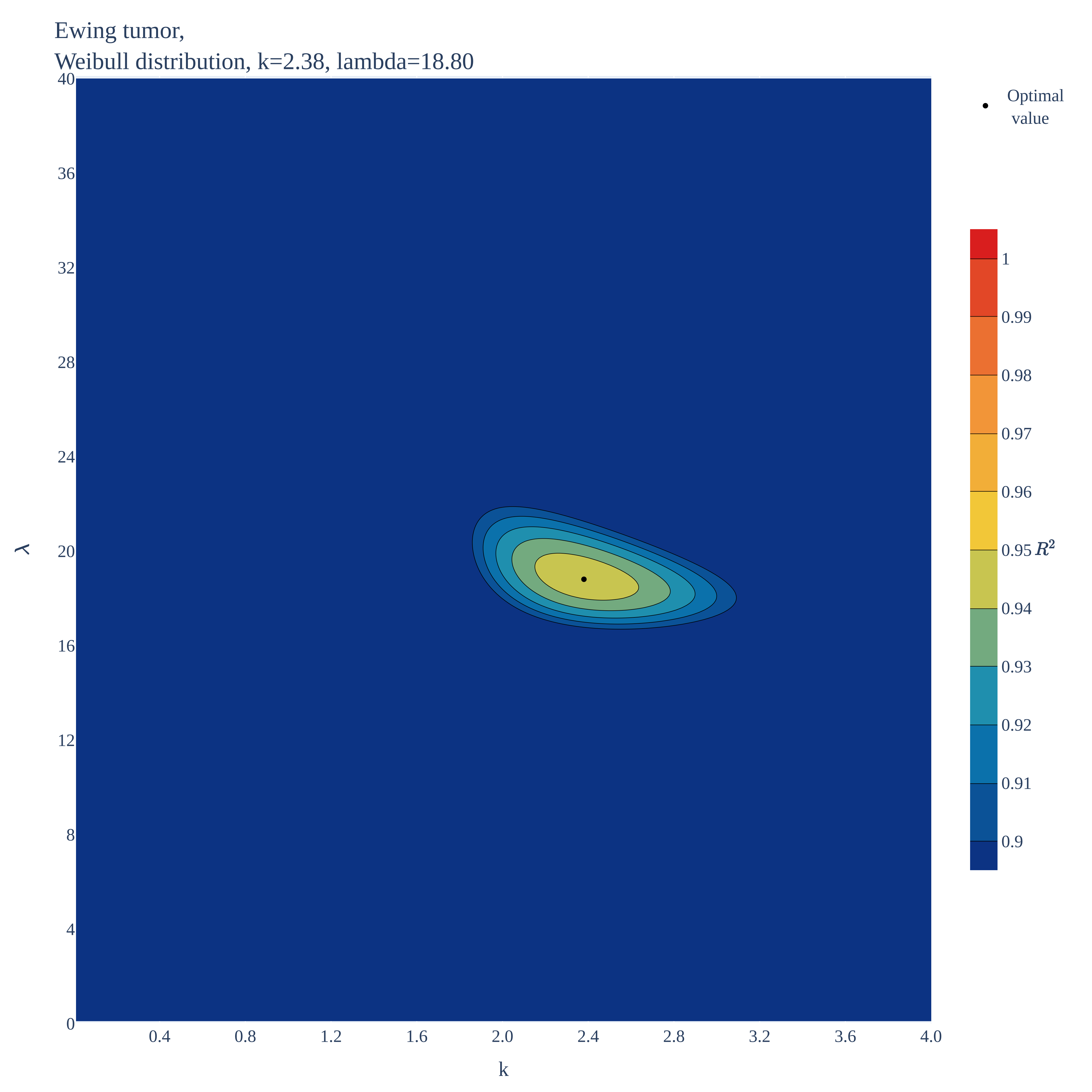

### Ewing tumor.pdf

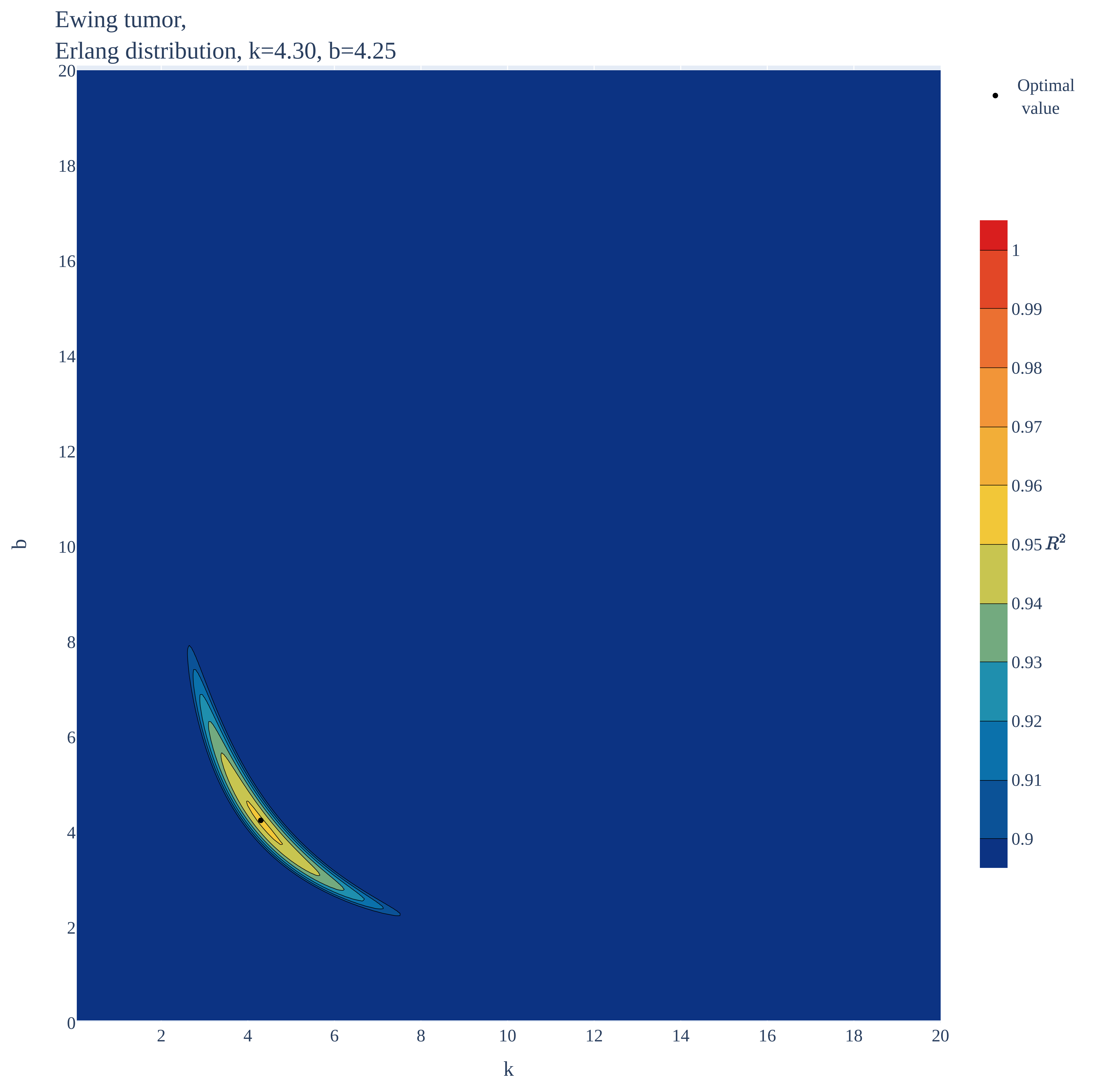

### Ewing tumor.pdf

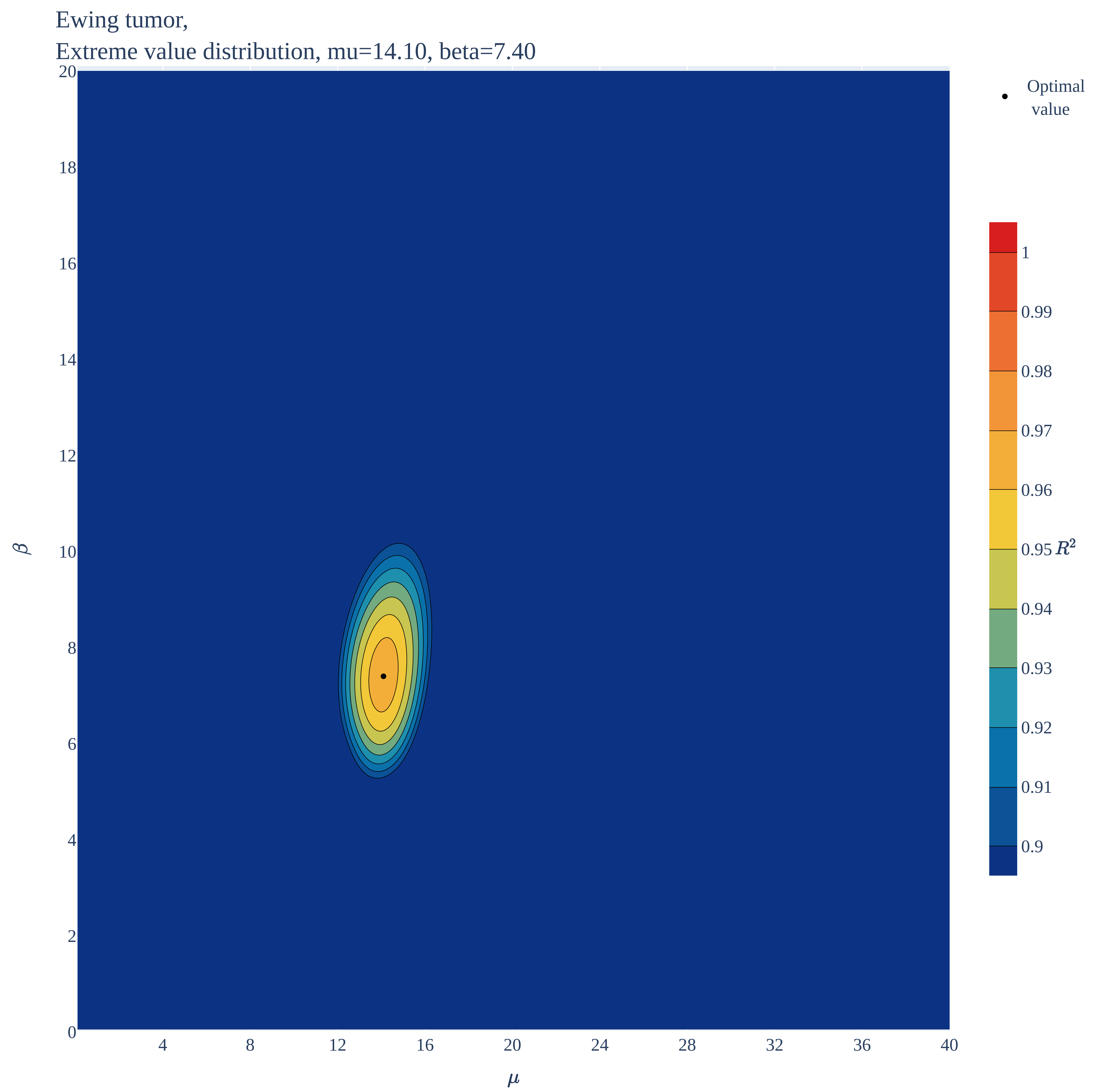

### Ewing tumor.pdf

Ewing tumor,
Normal distribution, mu=15.70, sigma=7.35

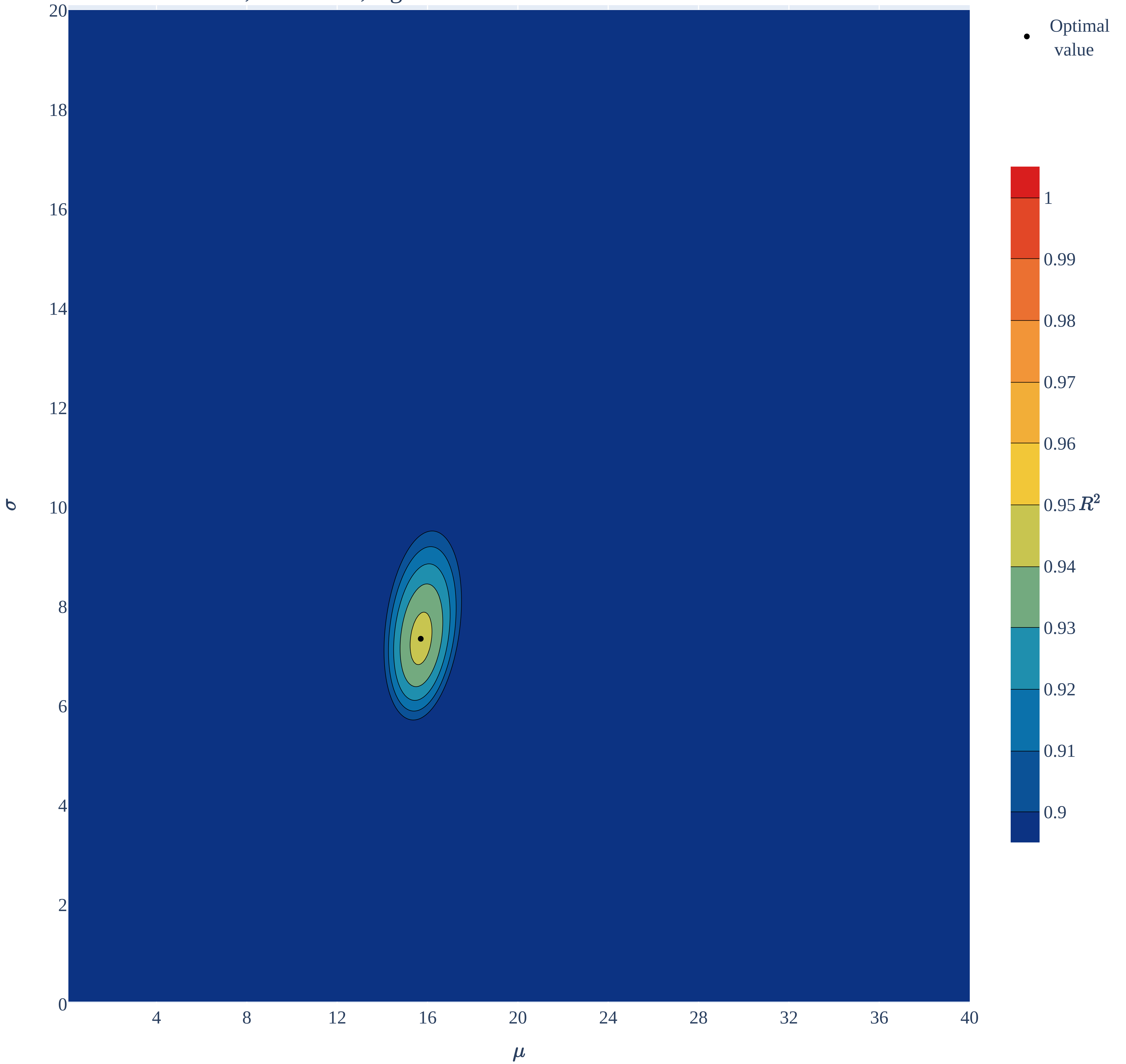

### Ewing tumor.pdf

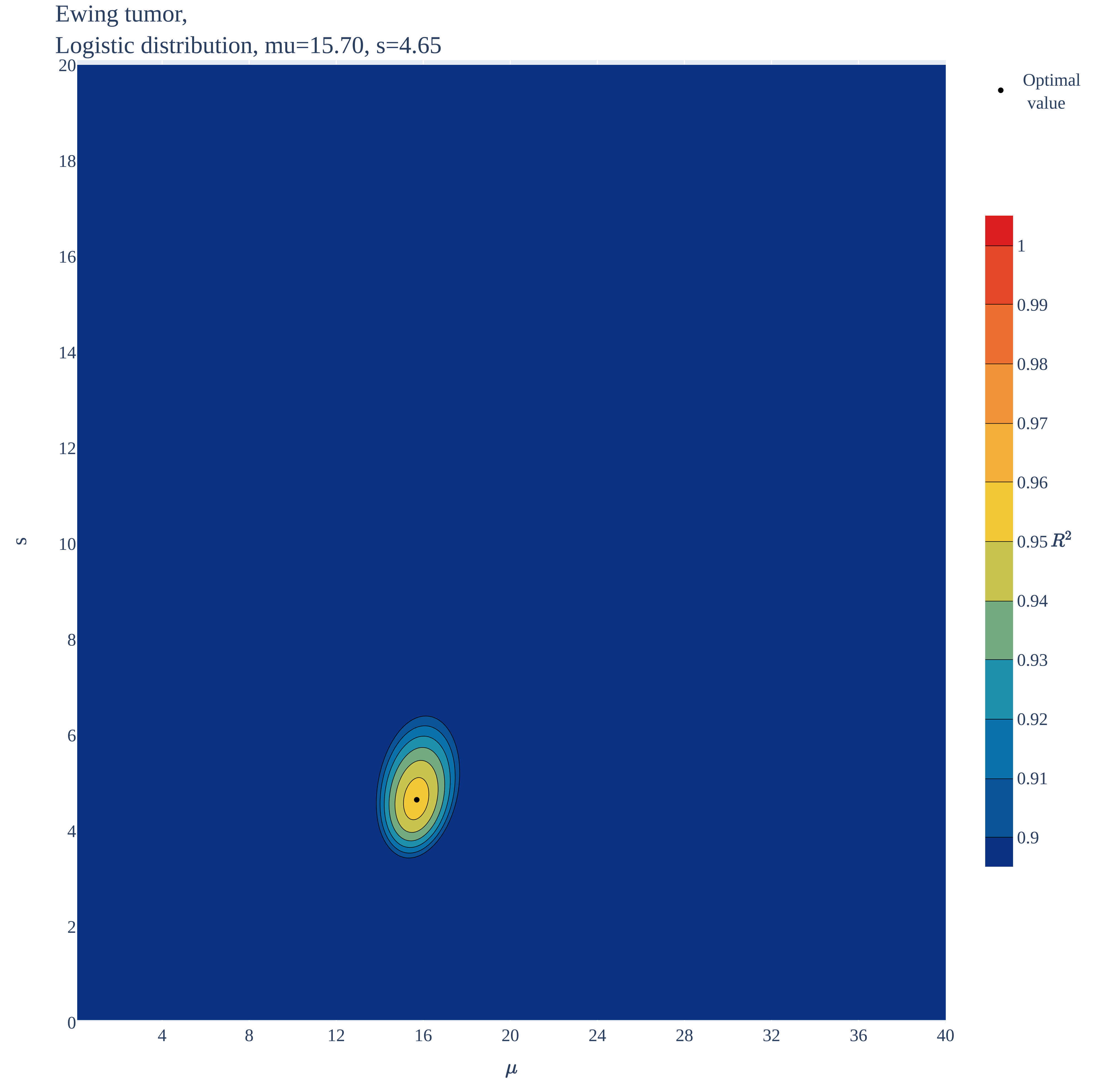

### Extracranial & extragonadal germ cell 1.pdf

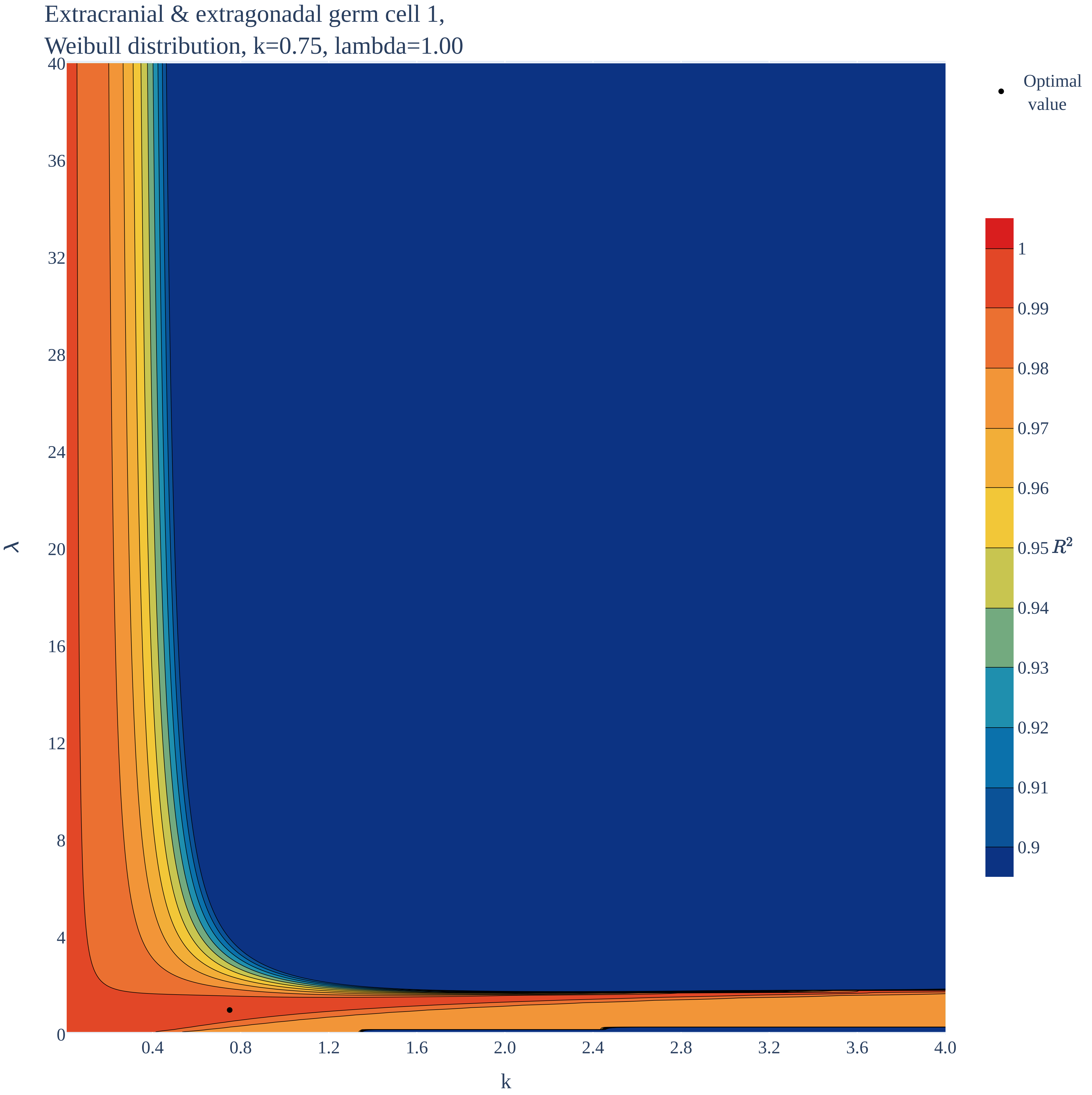

### Extracranial & extragonadal germ cell 1.pdf

Extracranial & extragonadal germ cell 1,
Erlang distribution, k=0.40, b=2.35

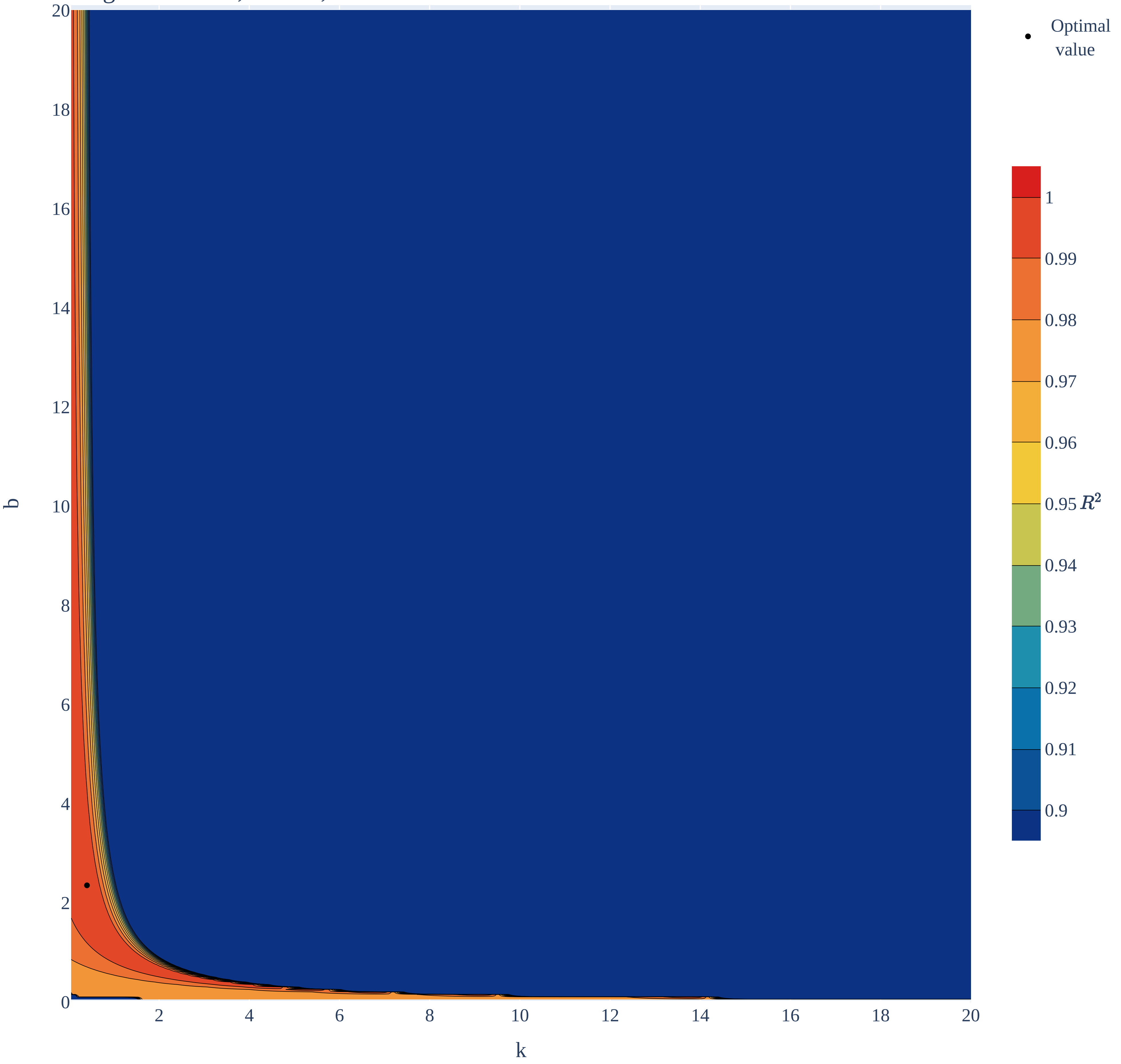

### Extracranial & extragonadal germ cell 1.pdf

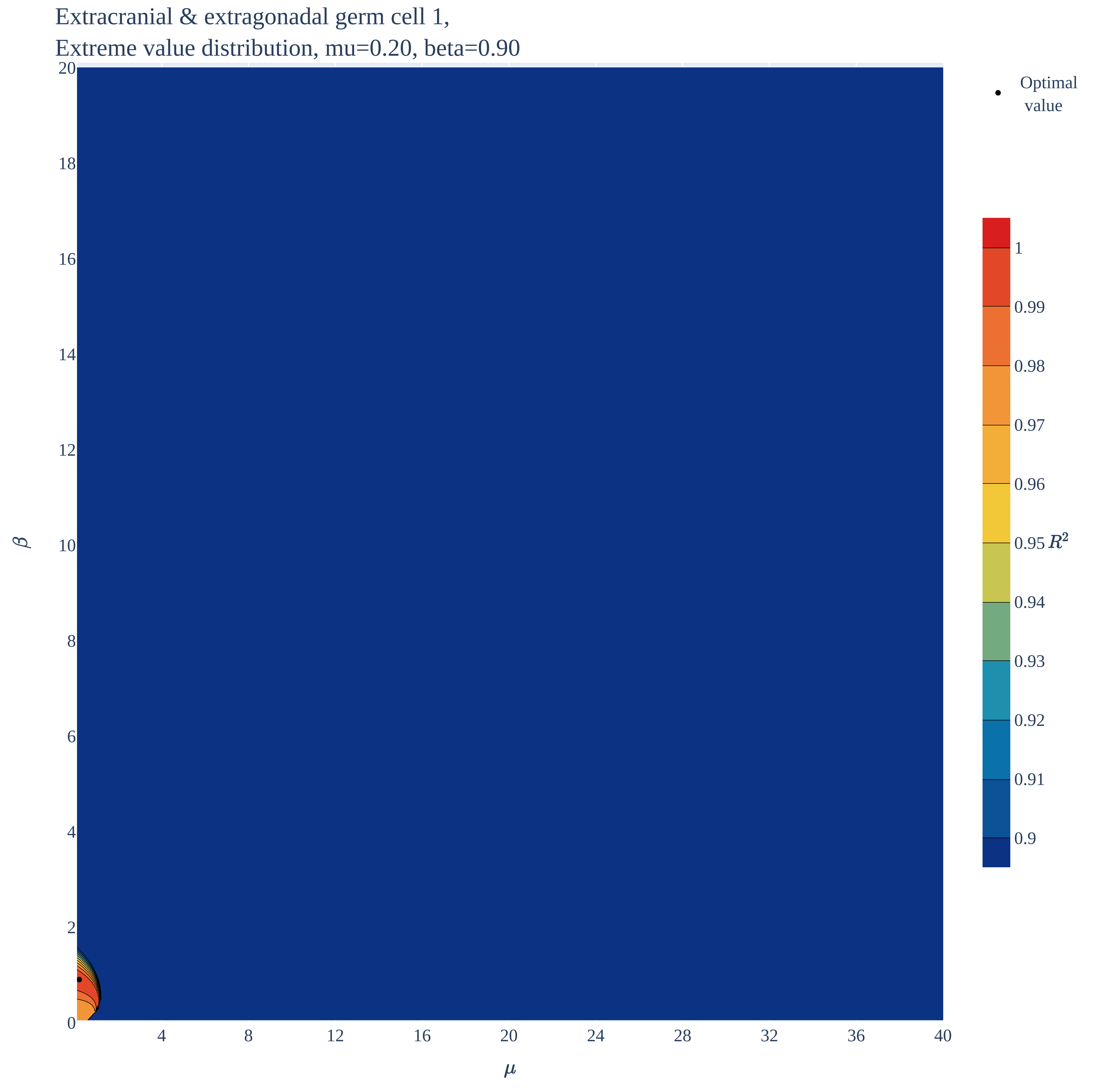

### Extracranial & extragonadal germ cell 1.pdf

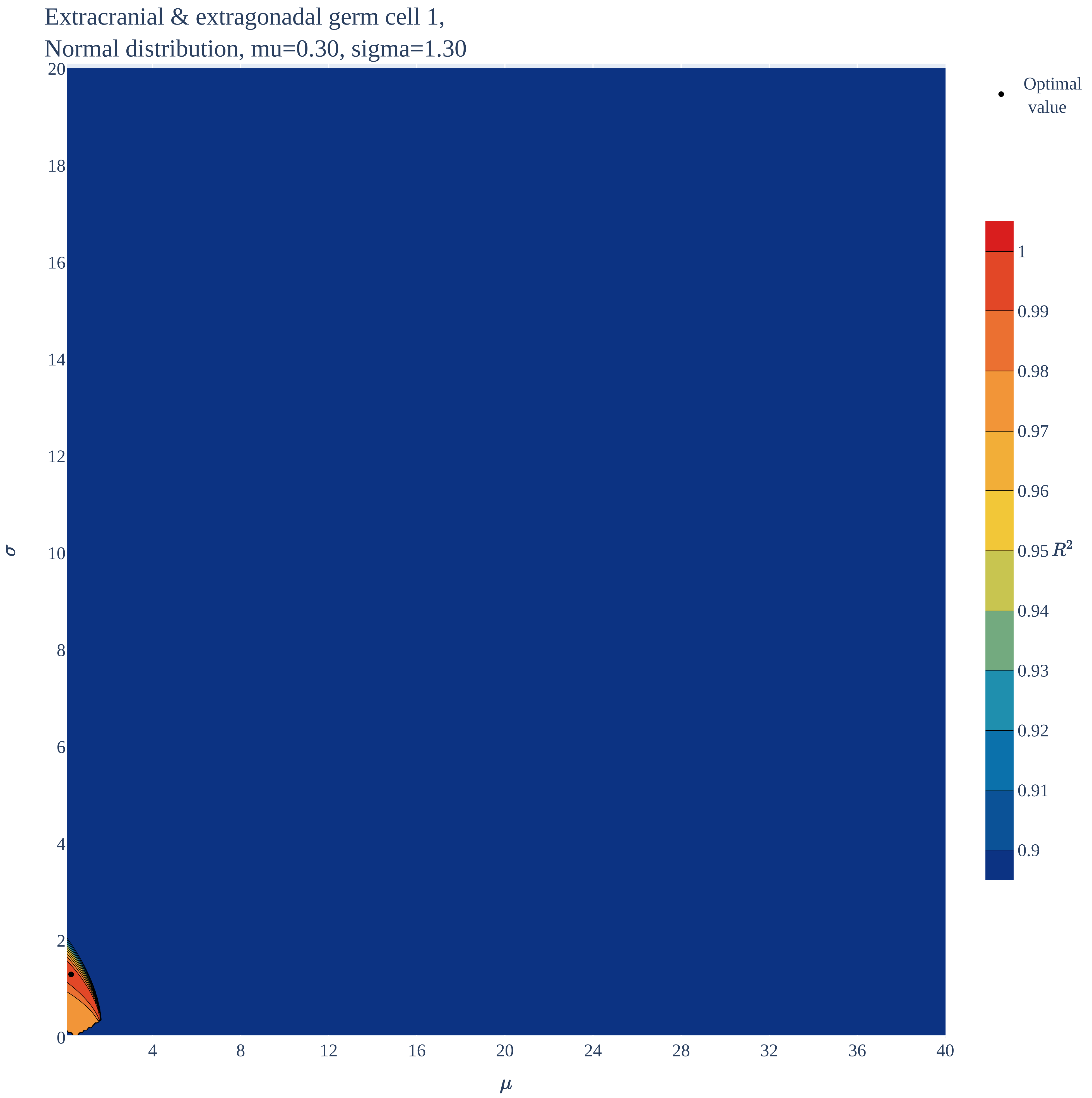

### Extracranial & extragonadal germ cell 1.pdf

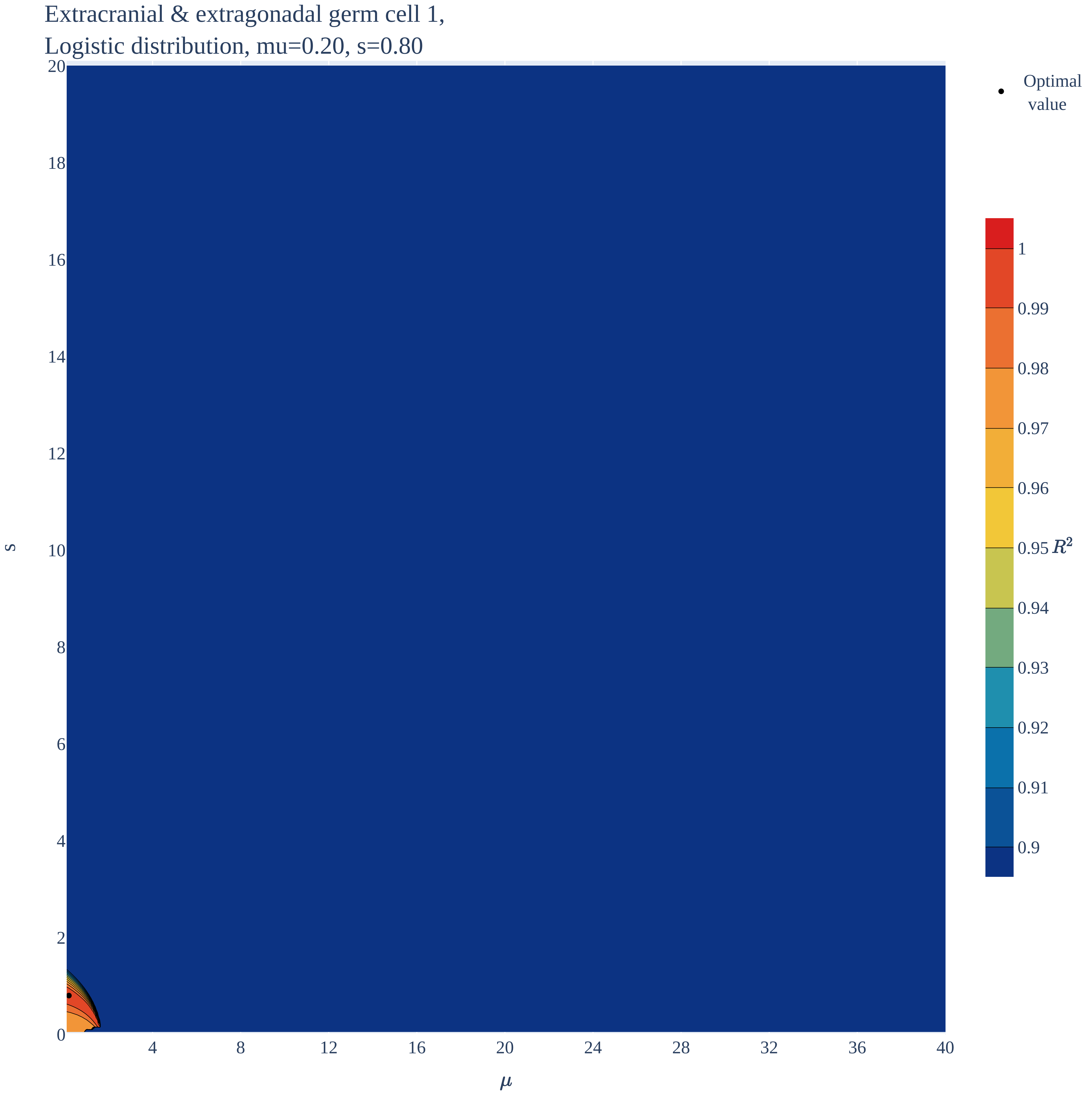

### Extracranial & extragonadal germ cell 2.pdf

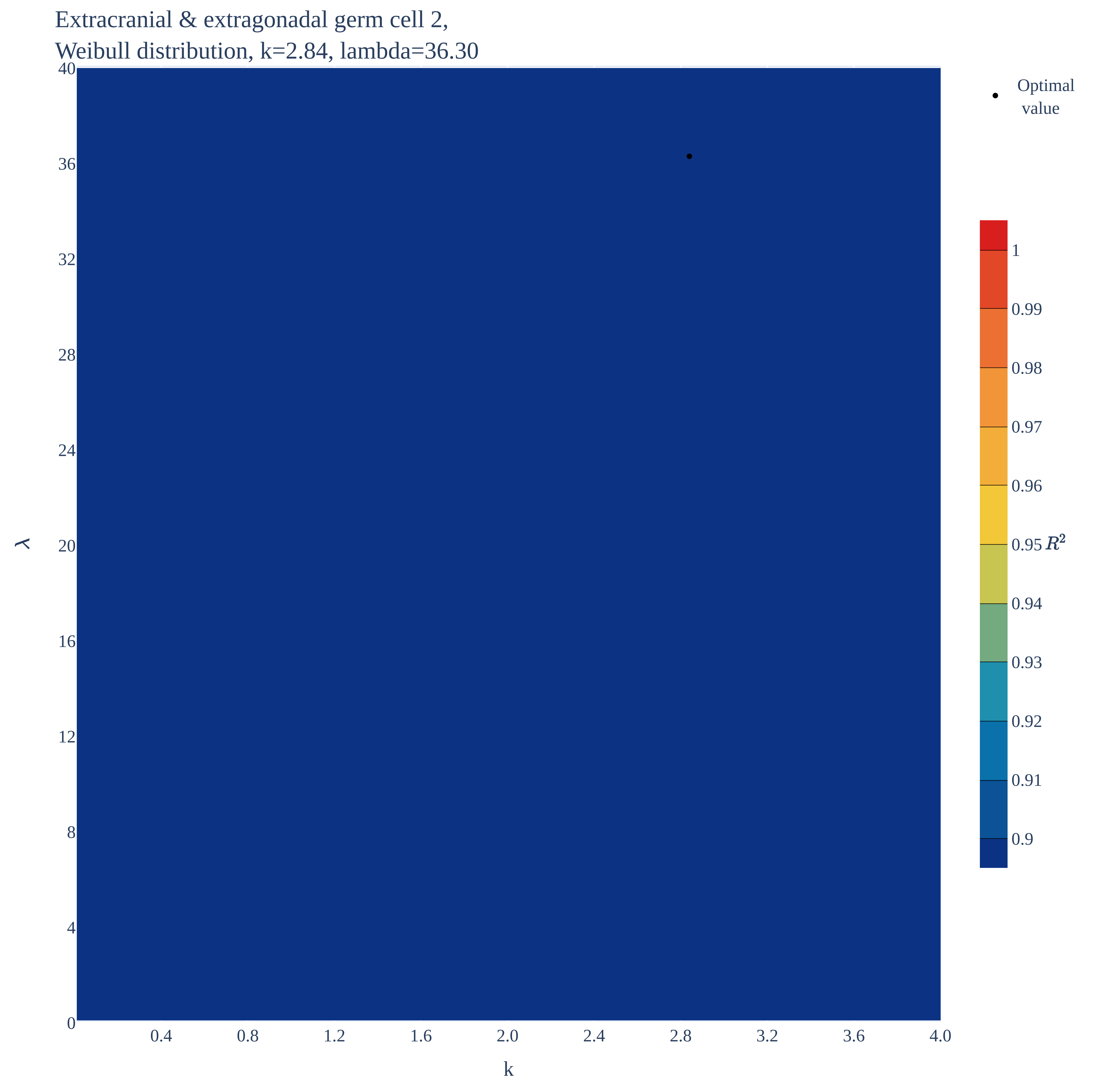

### Extracranial & extragonadal germ cell 2.pdf

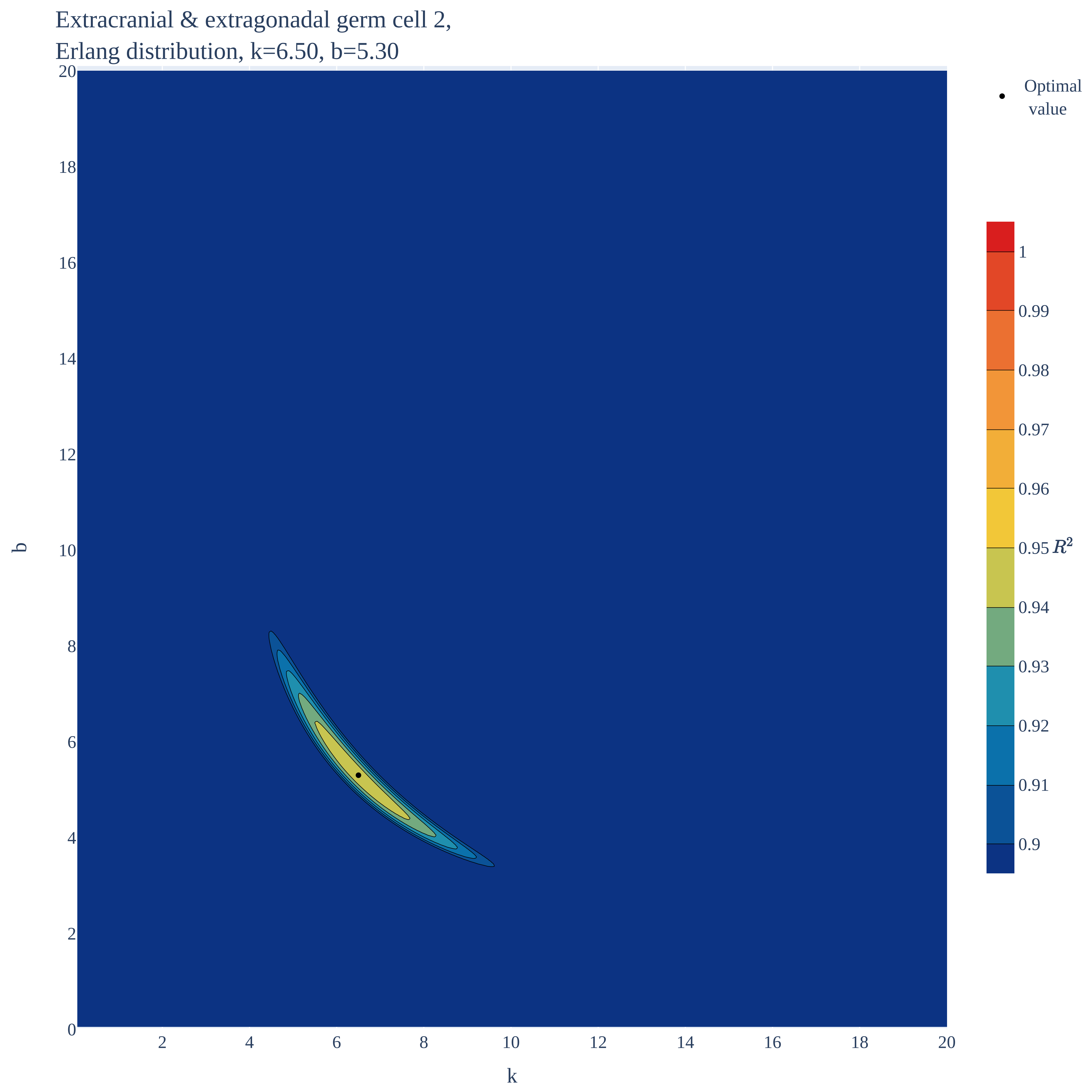

### Extracranial & extragonadal germ cell 2.pdf

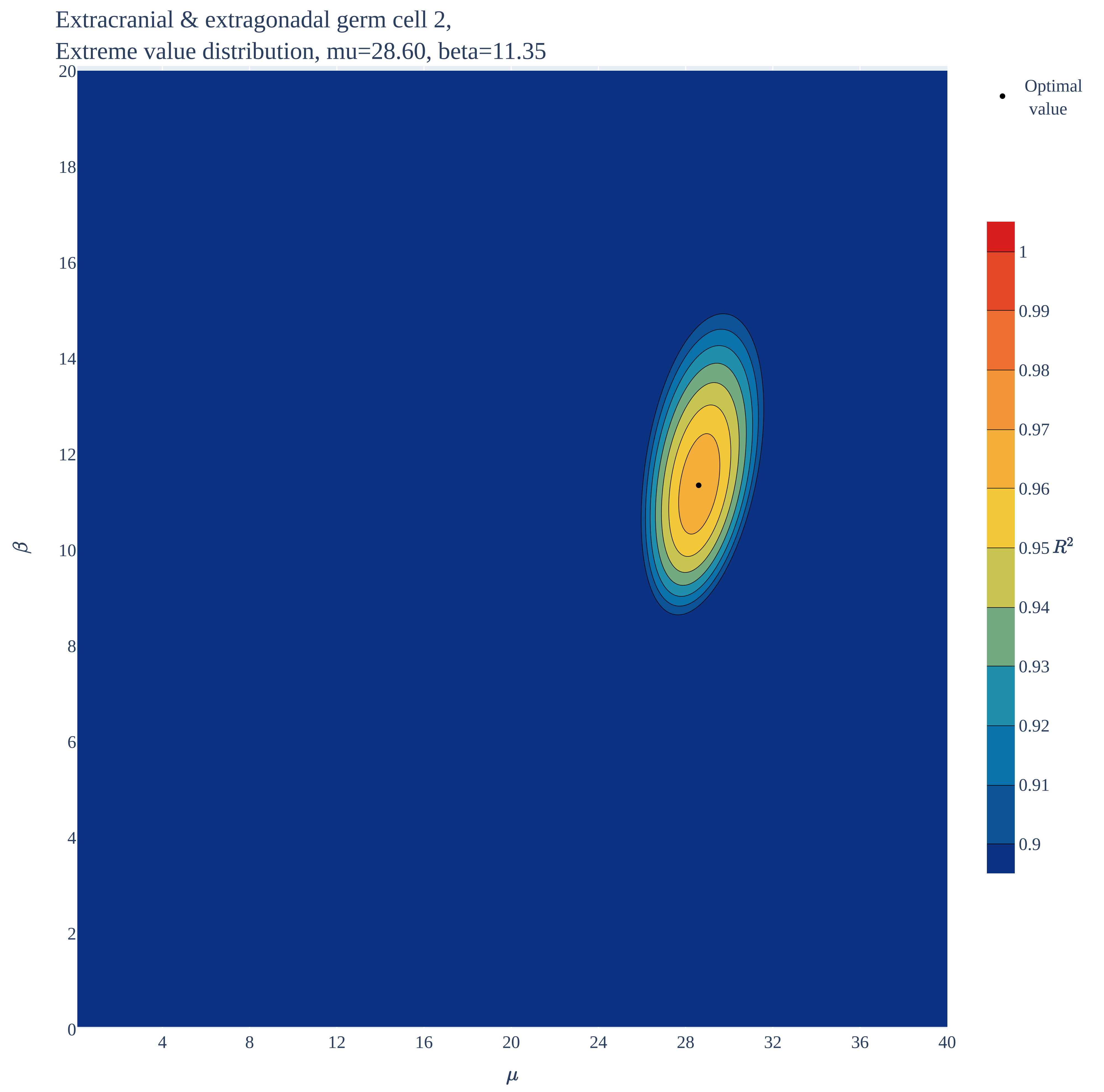

### Extracranial & extragonadal germ cell 2.pdf

Extracranial & extragonadal germ cell 2,
Normal distribution, mu=31.40, sigma=12.20

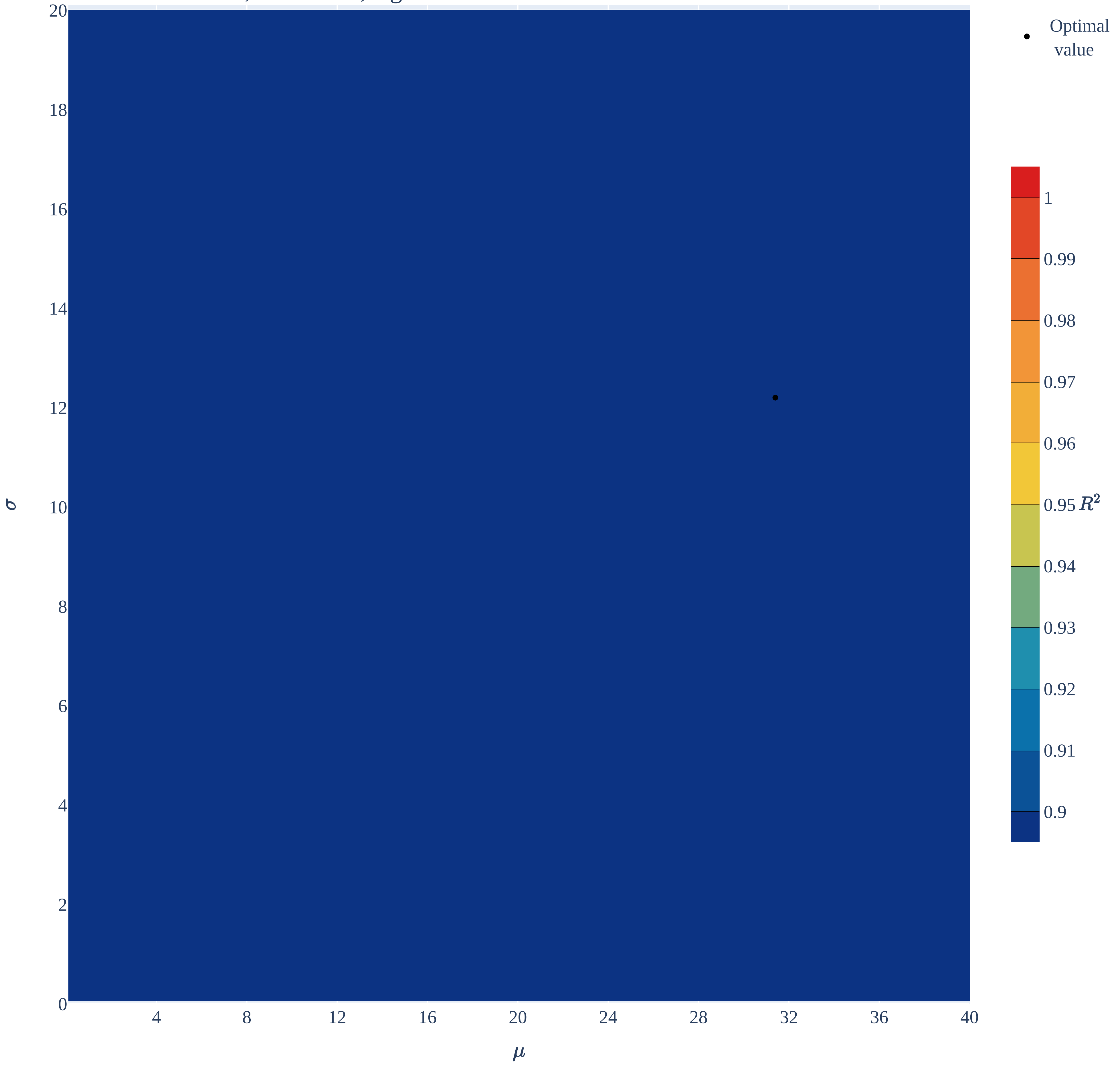

### Extracranial & extragonadal germ cell 2.pdf

Extracranial & extragonadal germ cell 2,
Logistic distribution, mu=31.10, s=7.90

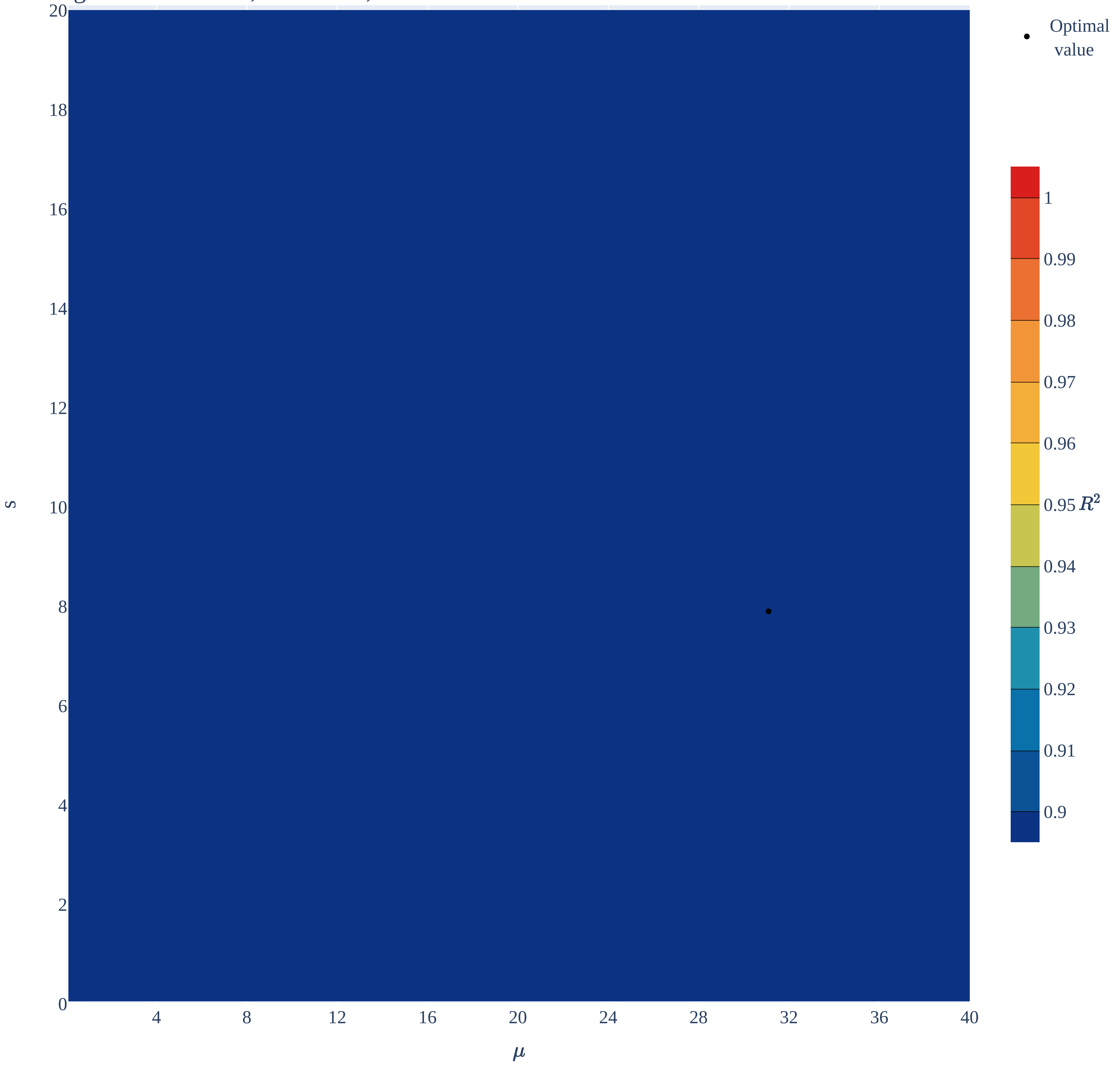

### Hepatoblastoma.pdf

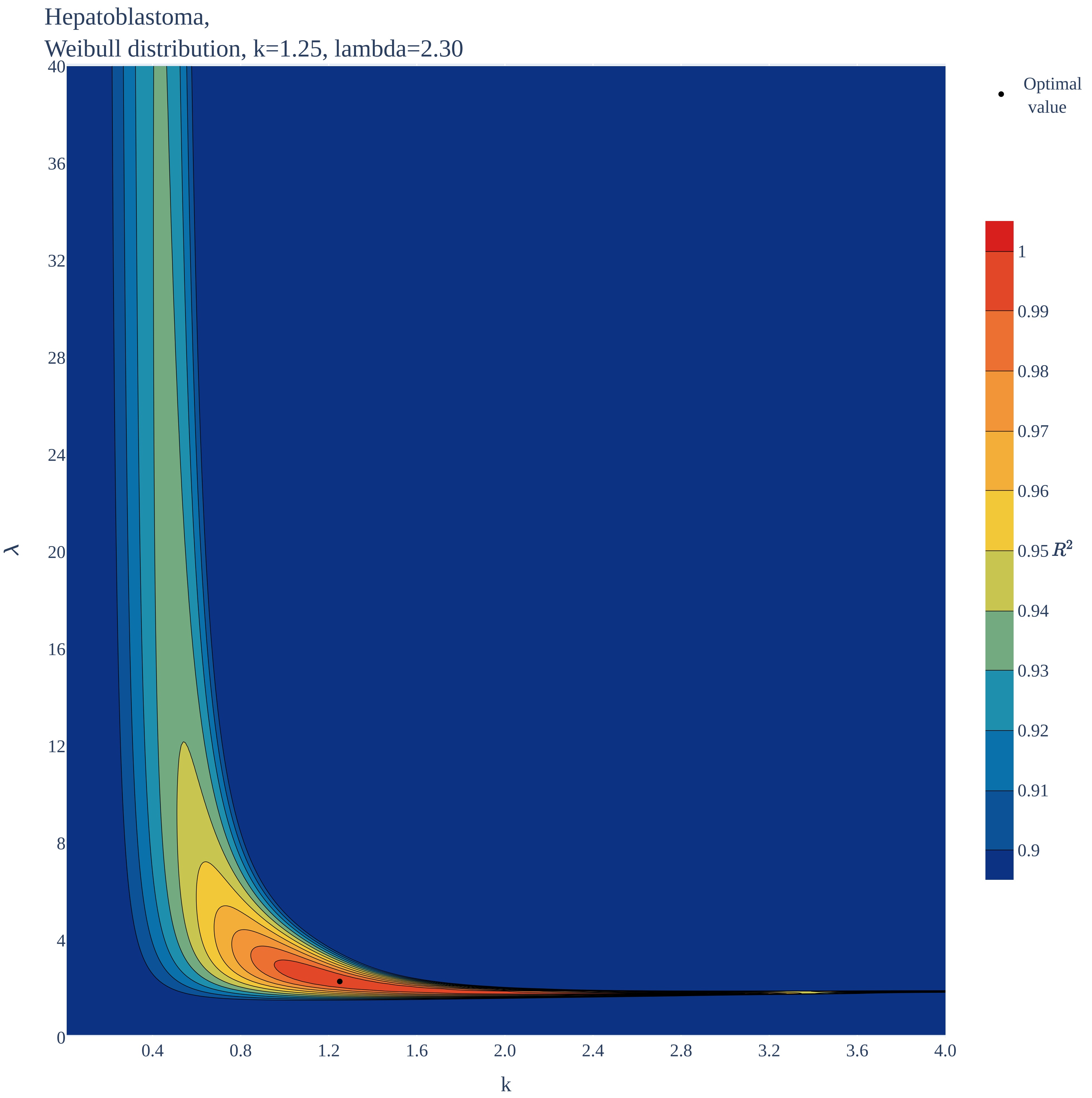

### Hepatoblastoma.pdf

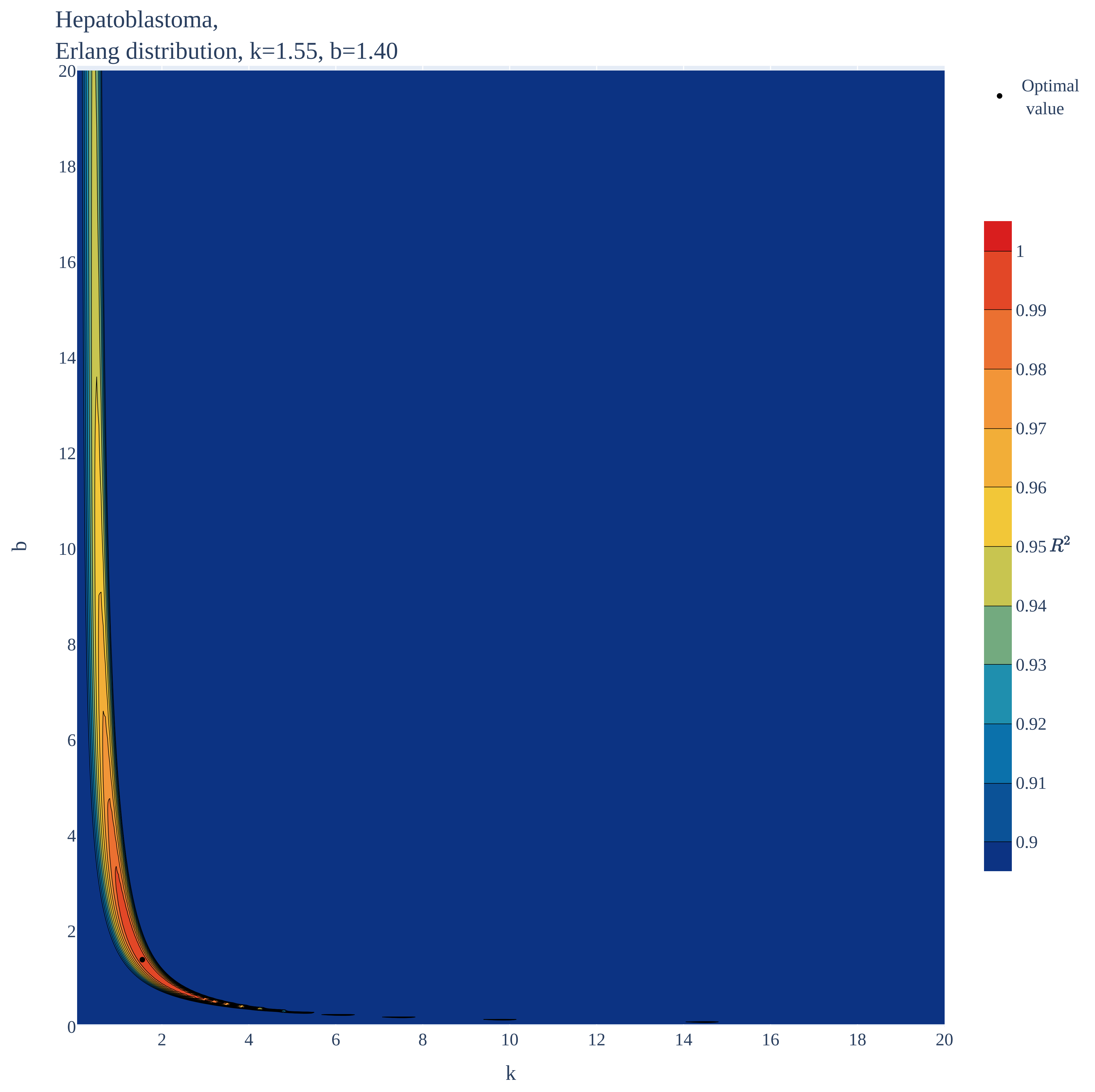

### Hepatoblastoma.pdf

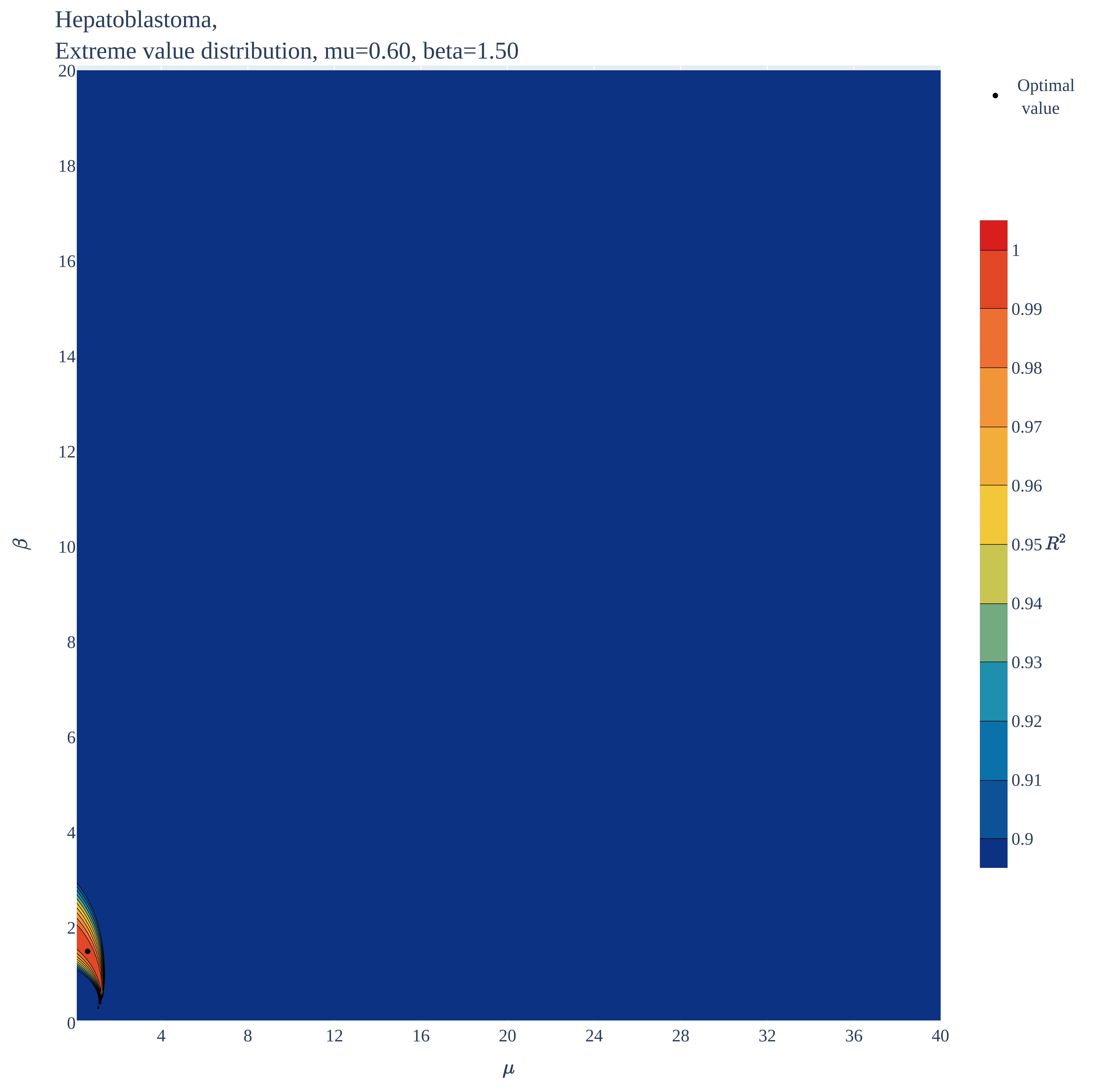

### Hepatoblastoma.pdf

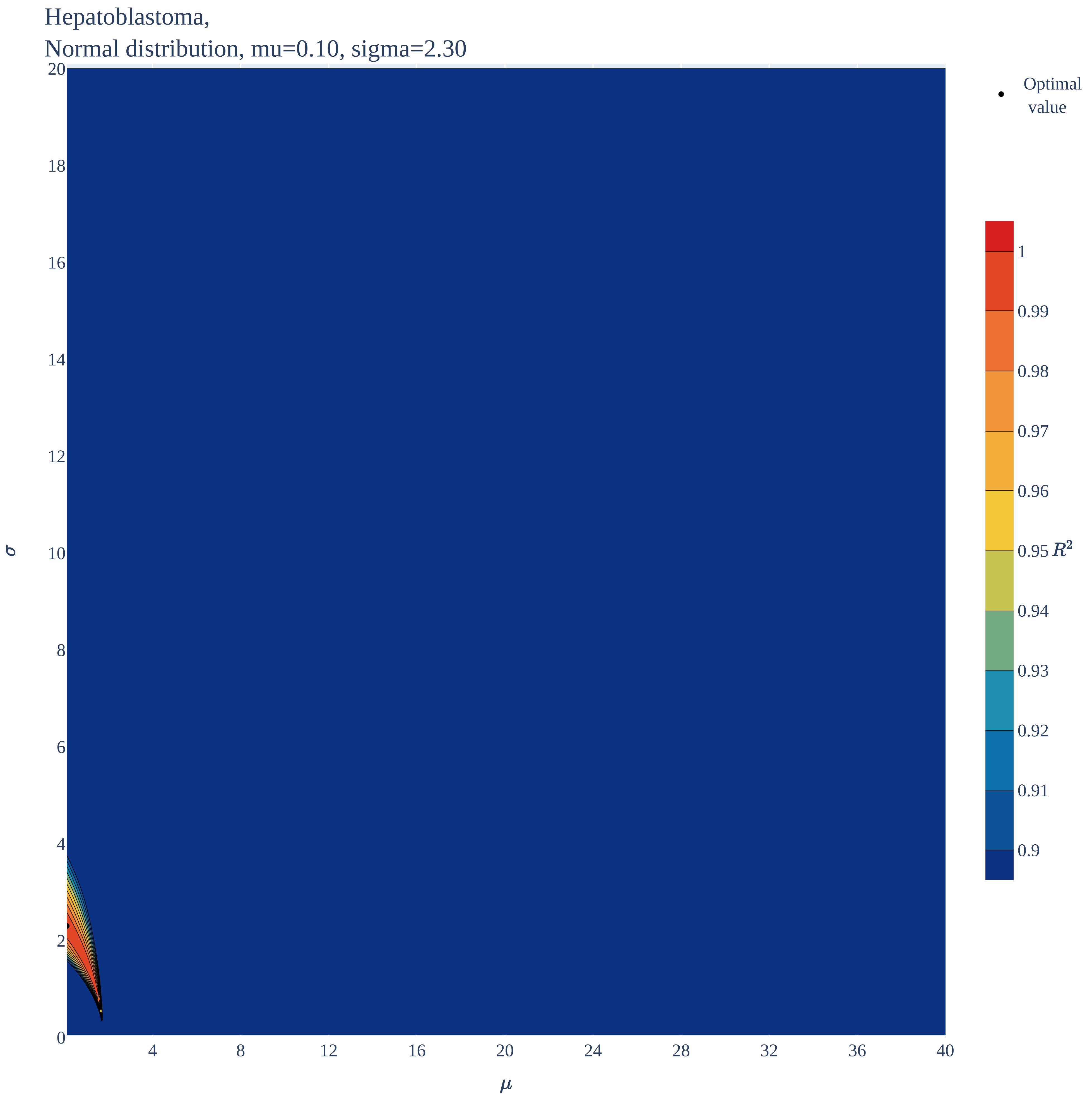

### Hepatoblastoma.pdf

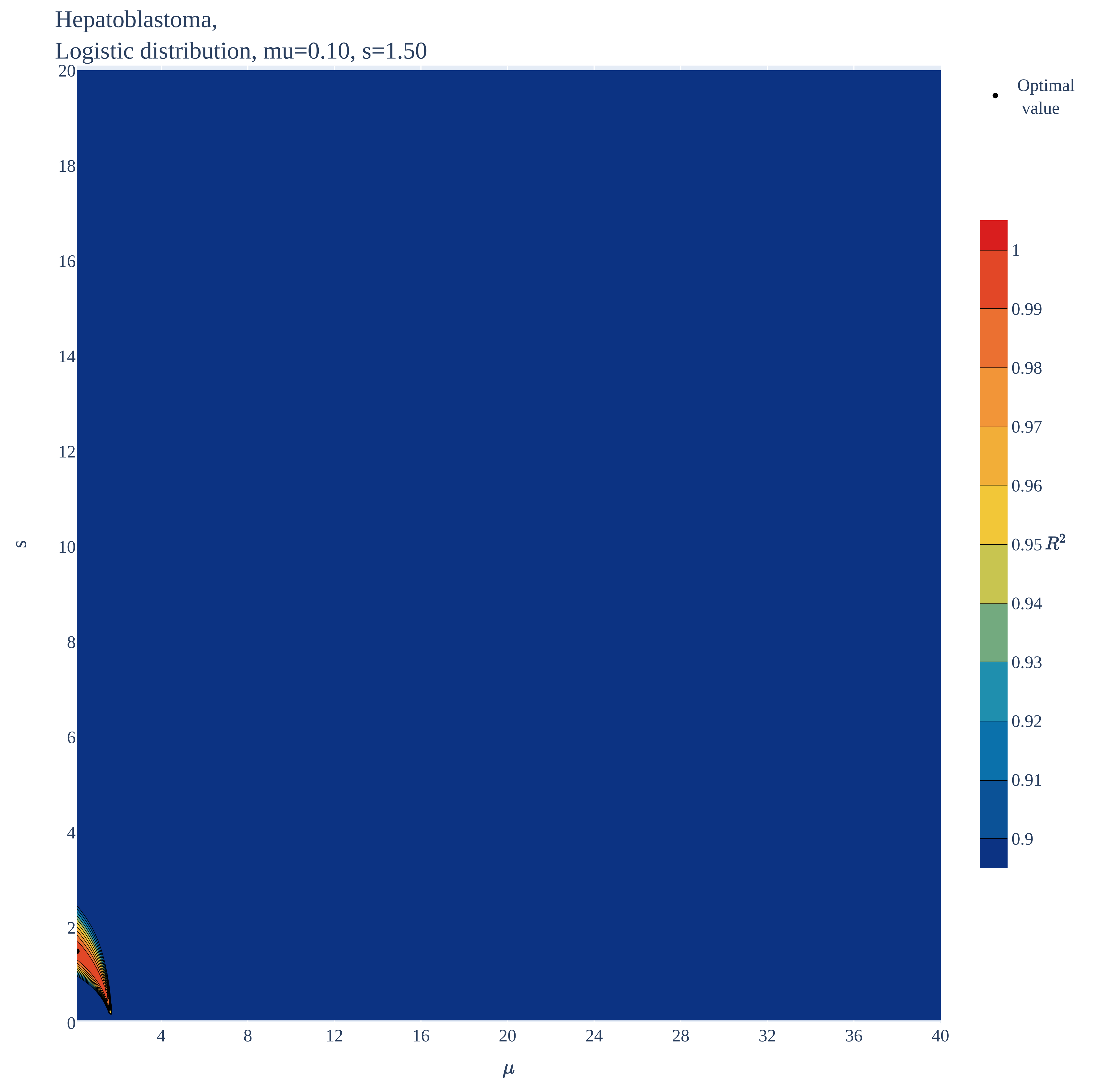

### Intracranial & intraspinal embryonal.pdf

Intracranial & intraspinal embryonal,
Weibull distribution, k=1.00, lambda=14.80

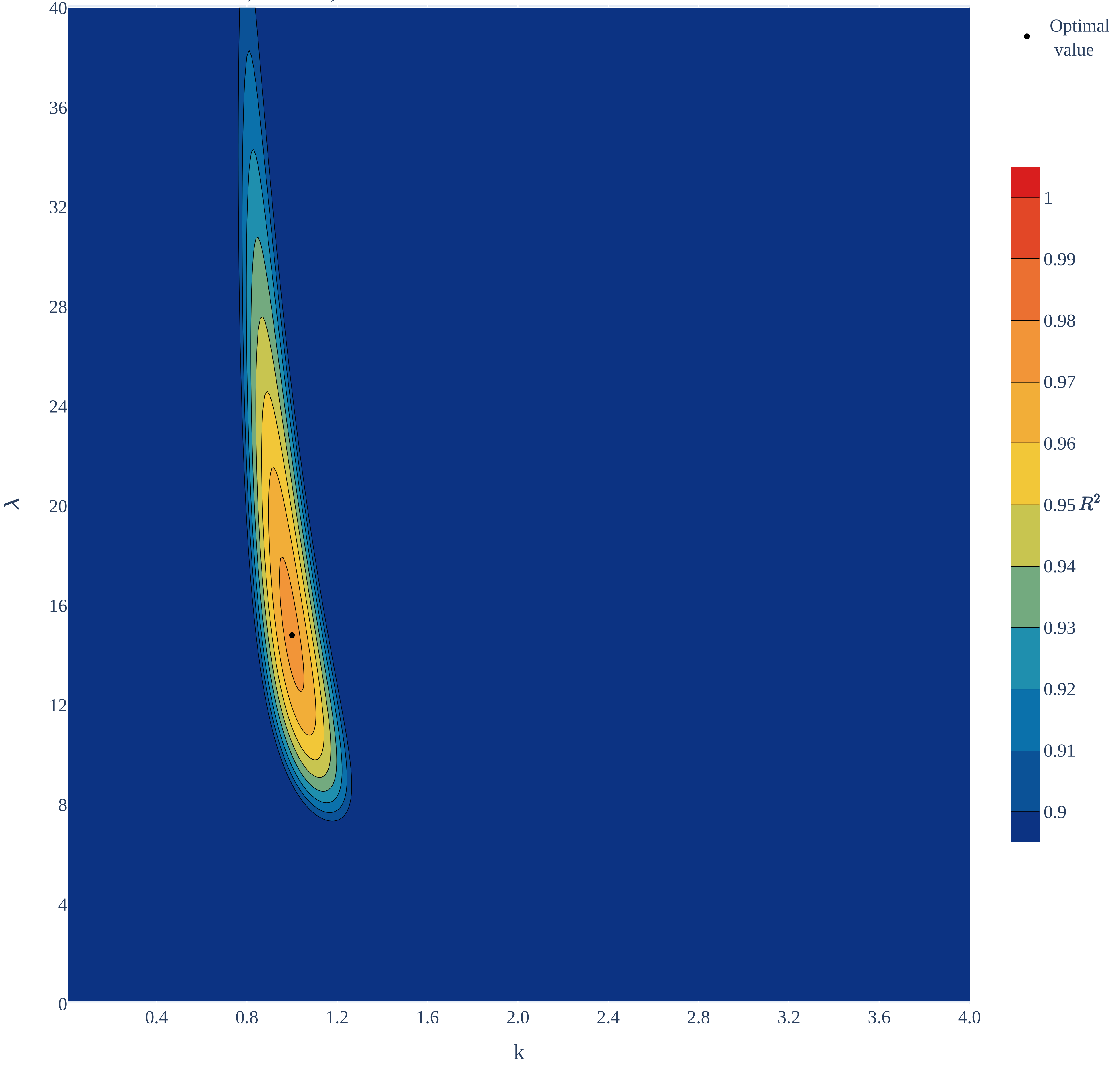

### Intracranial & intraspinal embryonal.pdf

Intracranial & intraspinal embryonal,
Erlang distribution, k=1.00, b=14.85

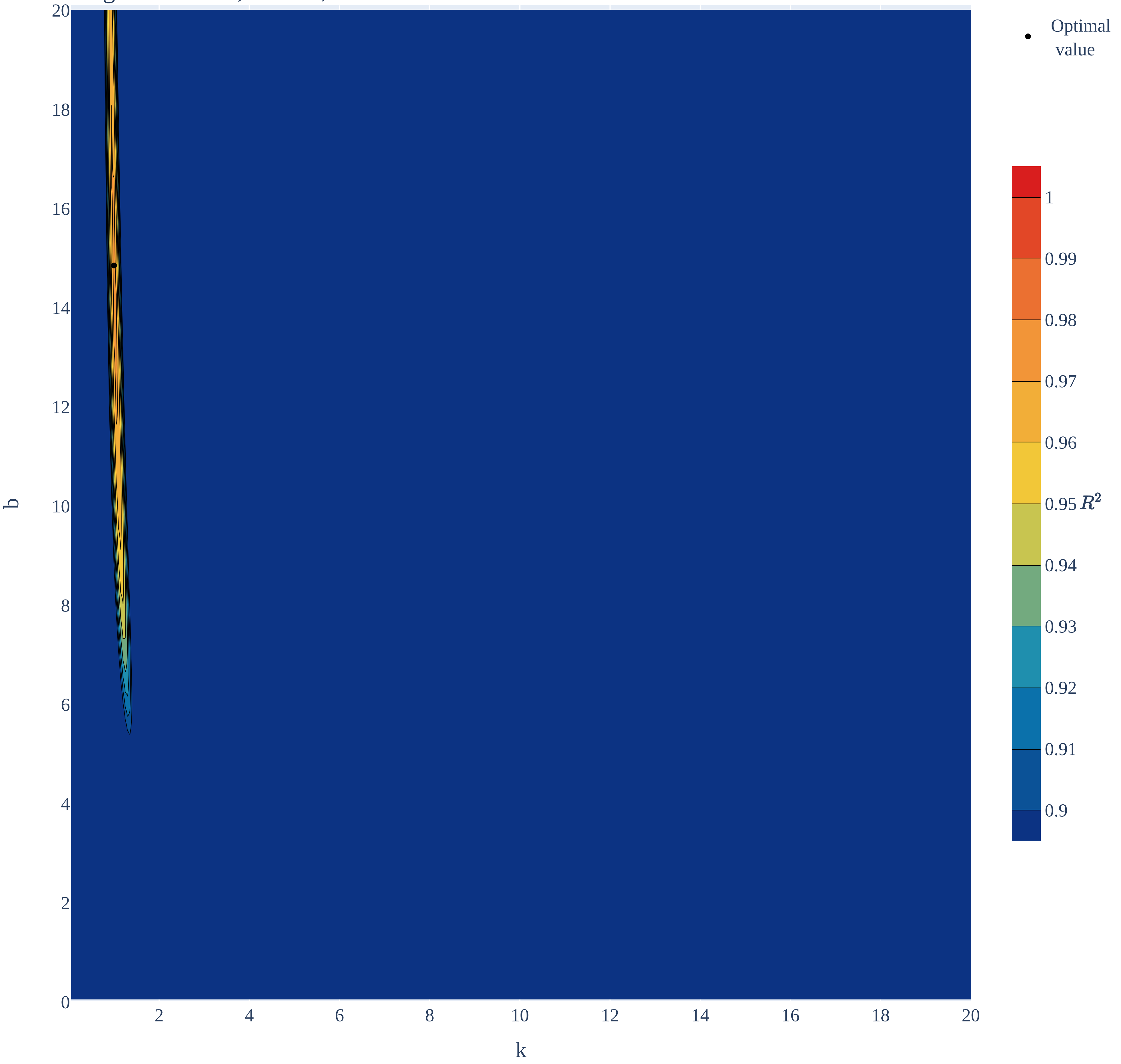

### Intracranial & intraspinal embryonal.pdf

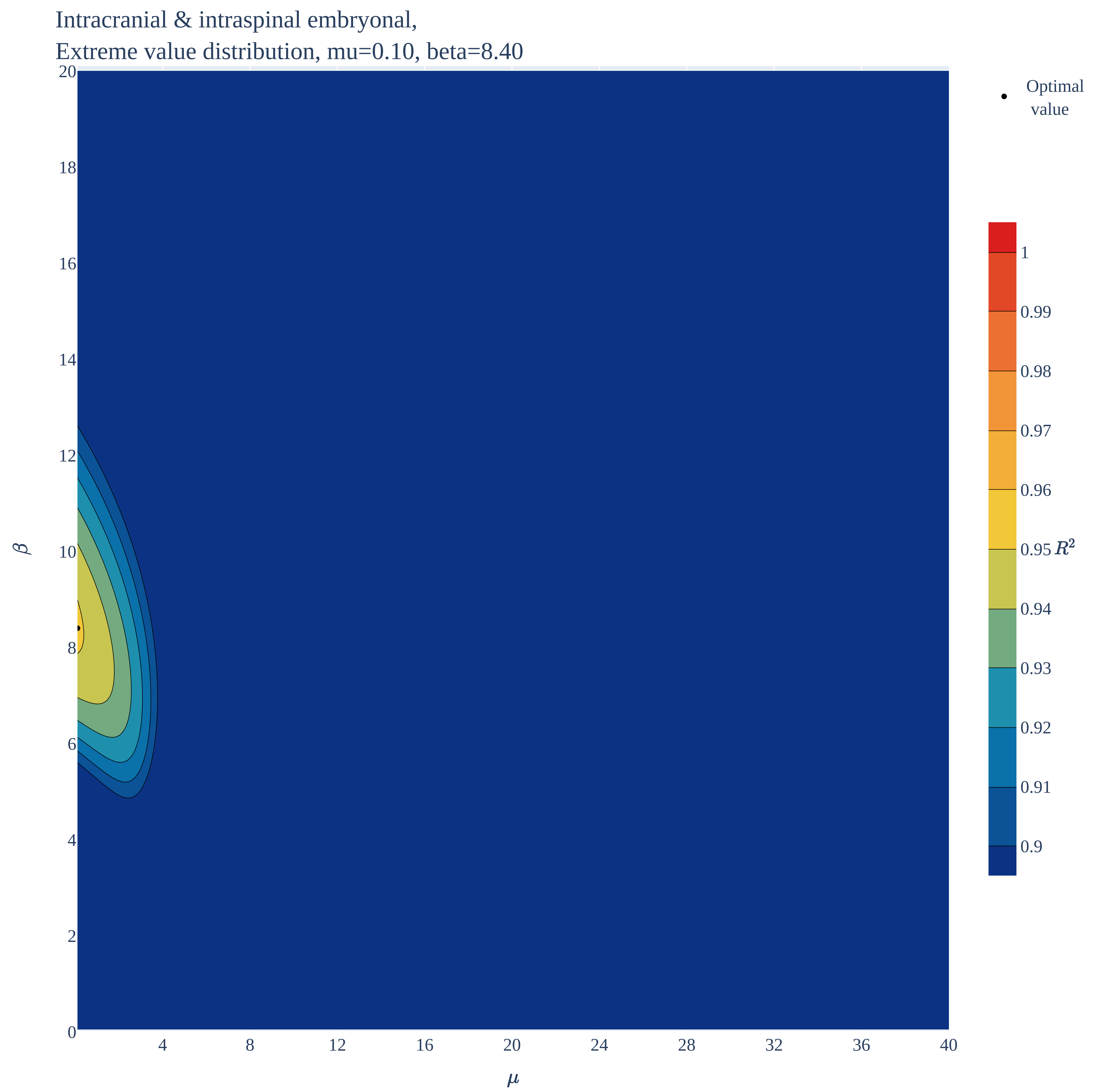

### Intracranial & intraspinal embryonal.pdf

Intracranial & intraspinal embryonal,
Normal distribution, mu=0.10, sigma=10.85

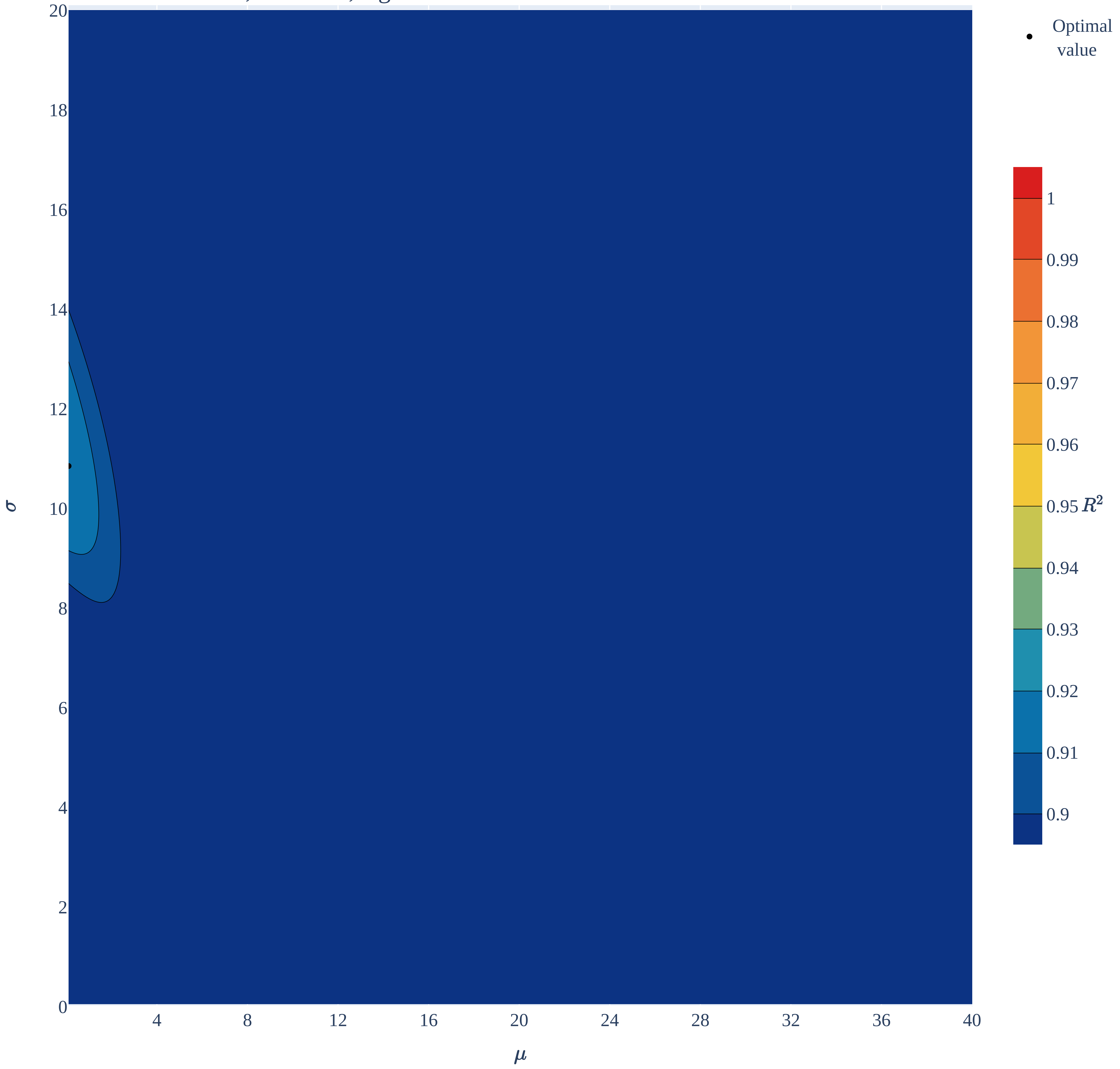

### Intracranial & intraspinal embryonal.pdf

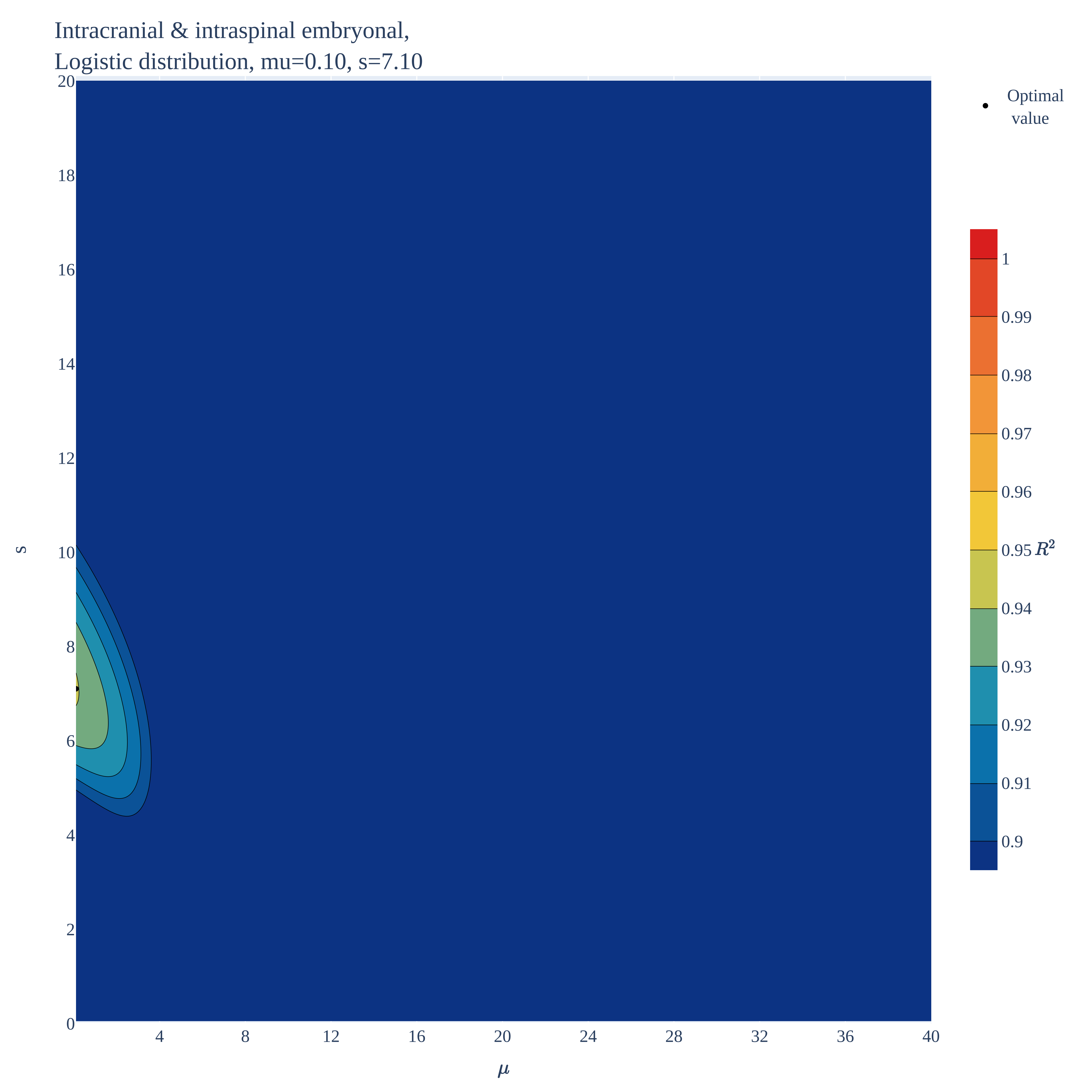

### Intracranial & intraspinal germ cell.pdf

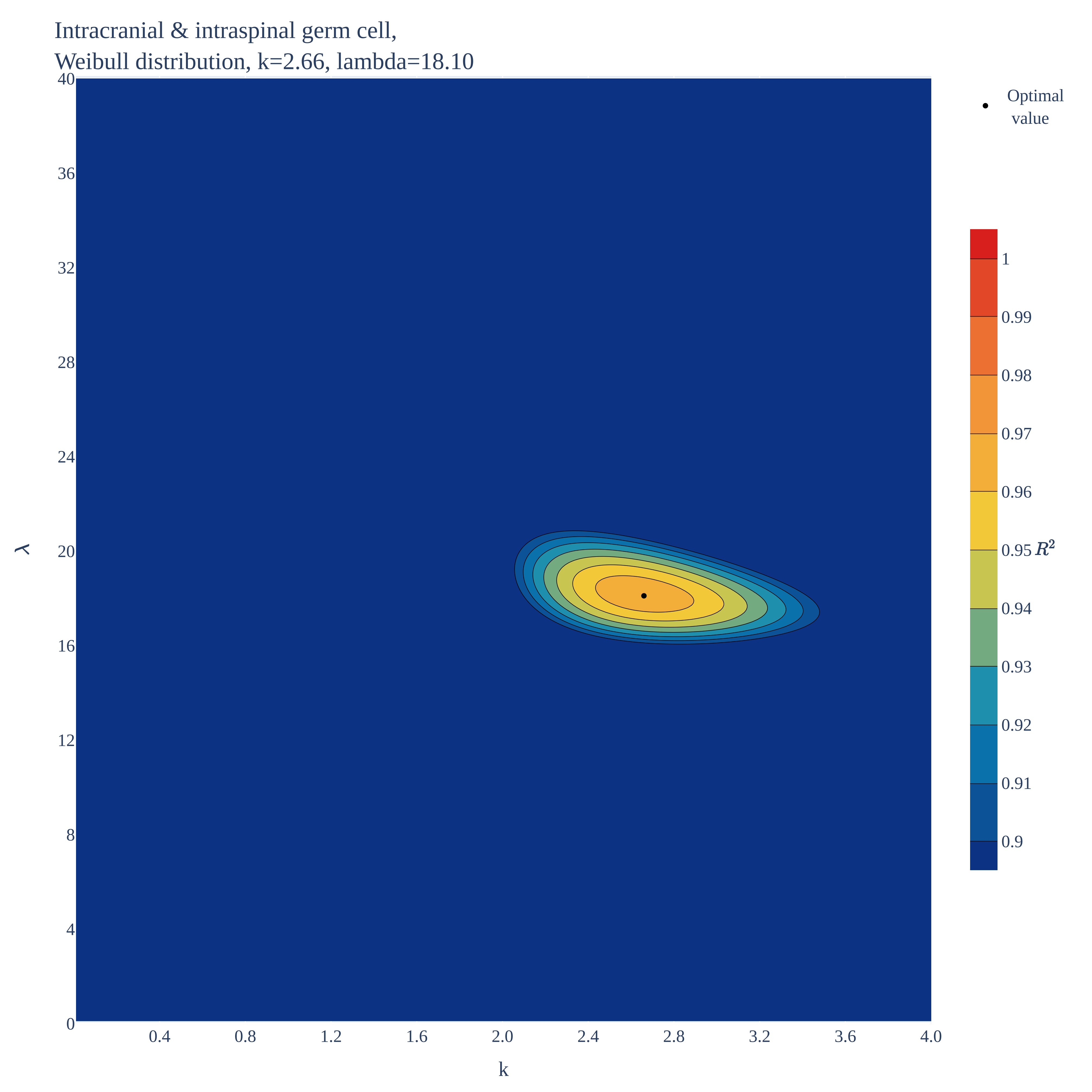

### Intracranial & intraspinal germ cell.pdf

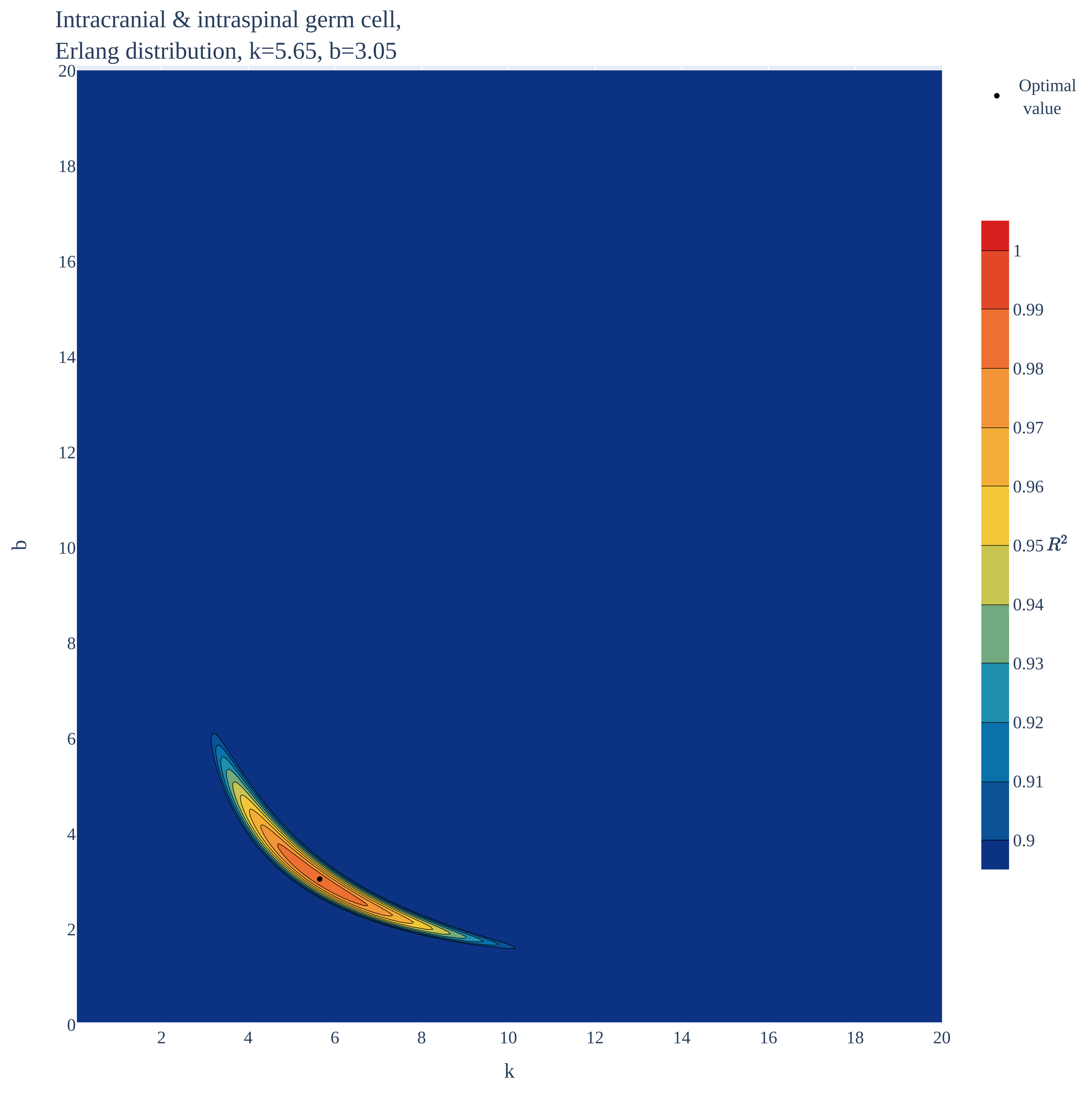

### Intracranial & intraspinal germ cell.pdf

Intracranial & intraspinal germ cell,
Extreme value distribution, mu=14.00, beta=6.20

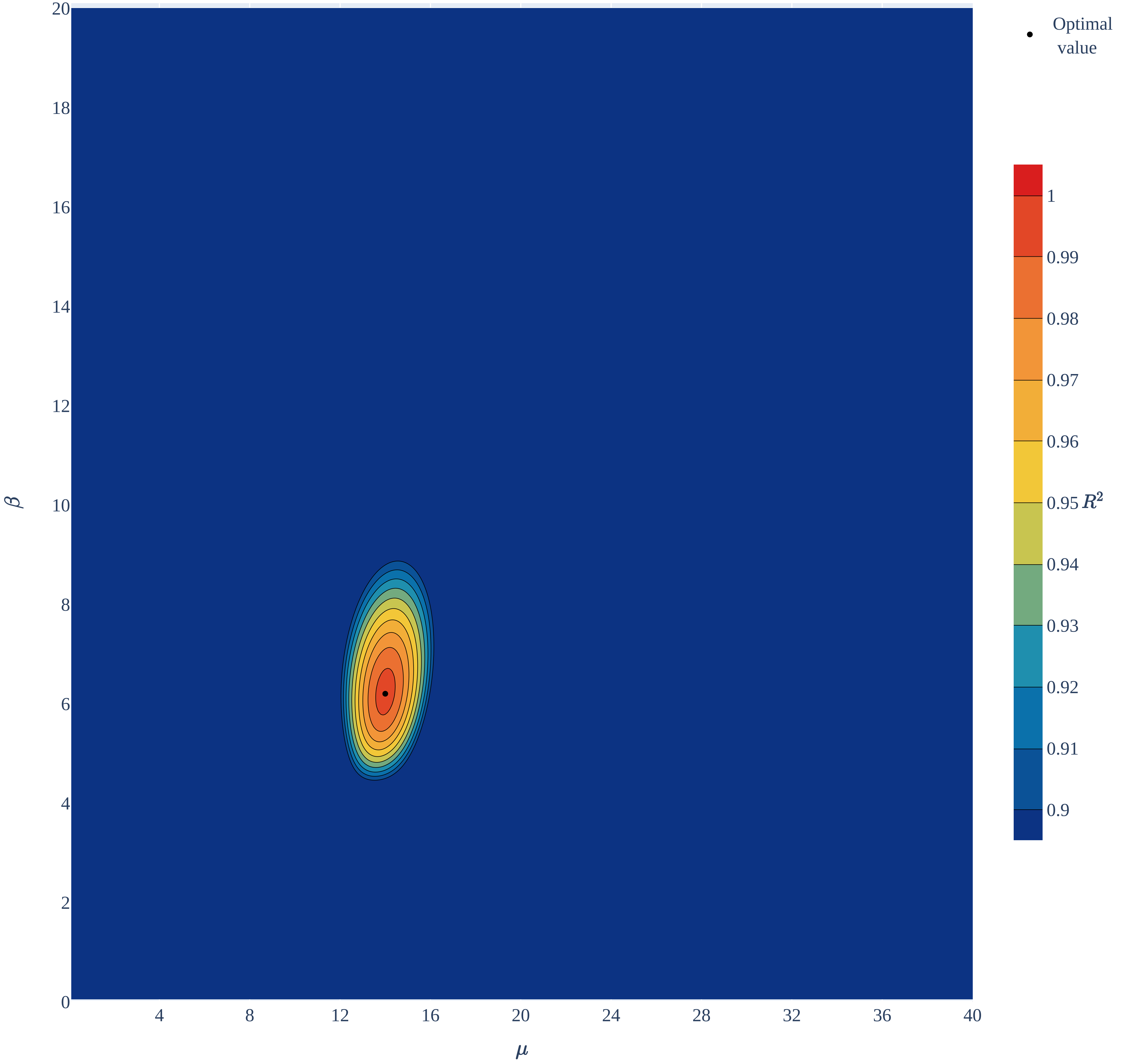

### Intracranial & intraspinal germ cell.pdf

Intracranial & intraspinal germ cell,
Normal distribution, mu=15.50, sigma=6.60

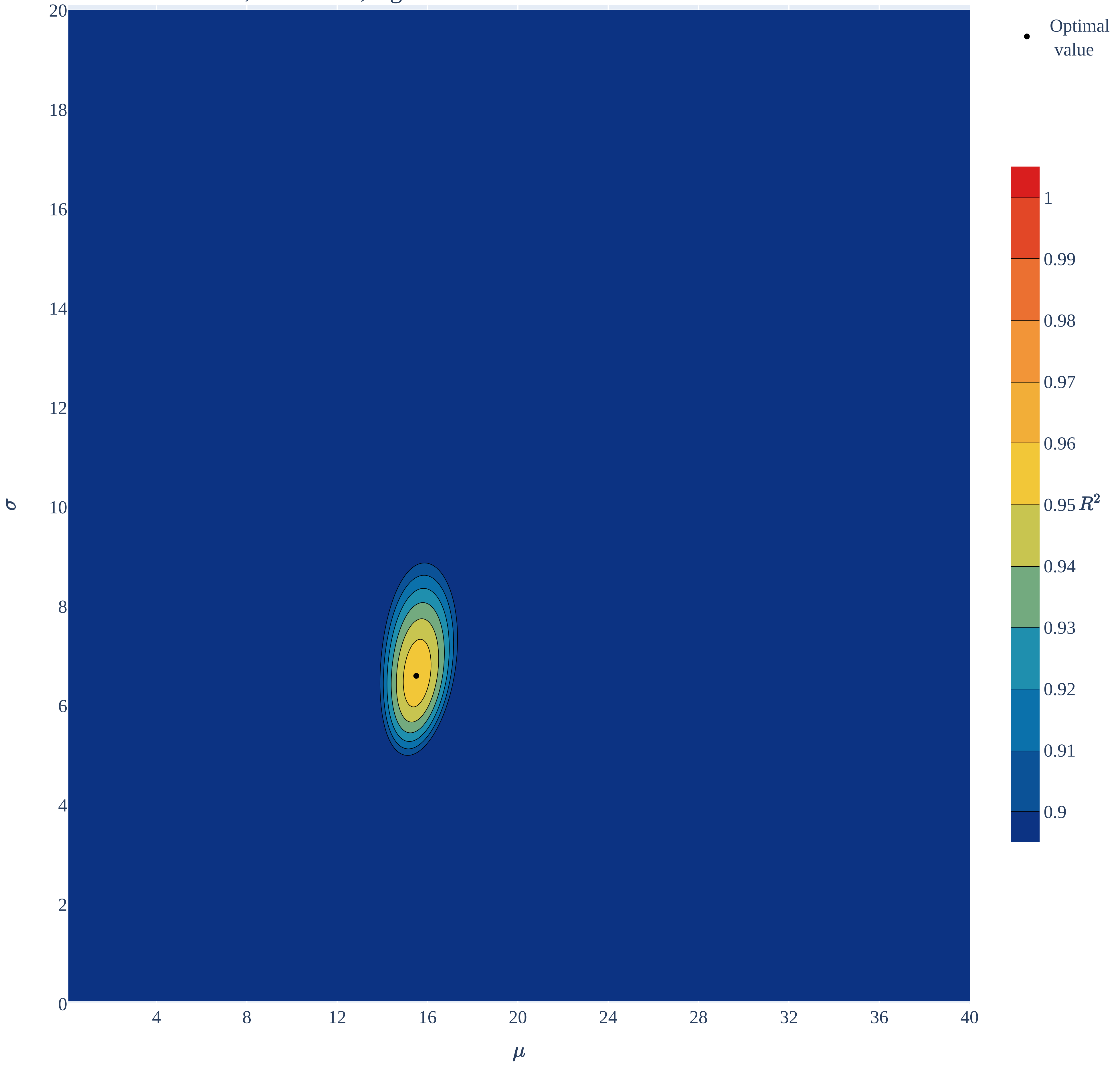

### Intracranial & intraspinal germ cell.pdf

Intracranial & intraspinal germ cell,
Logistic distribution, mu=15.40, s=4.20

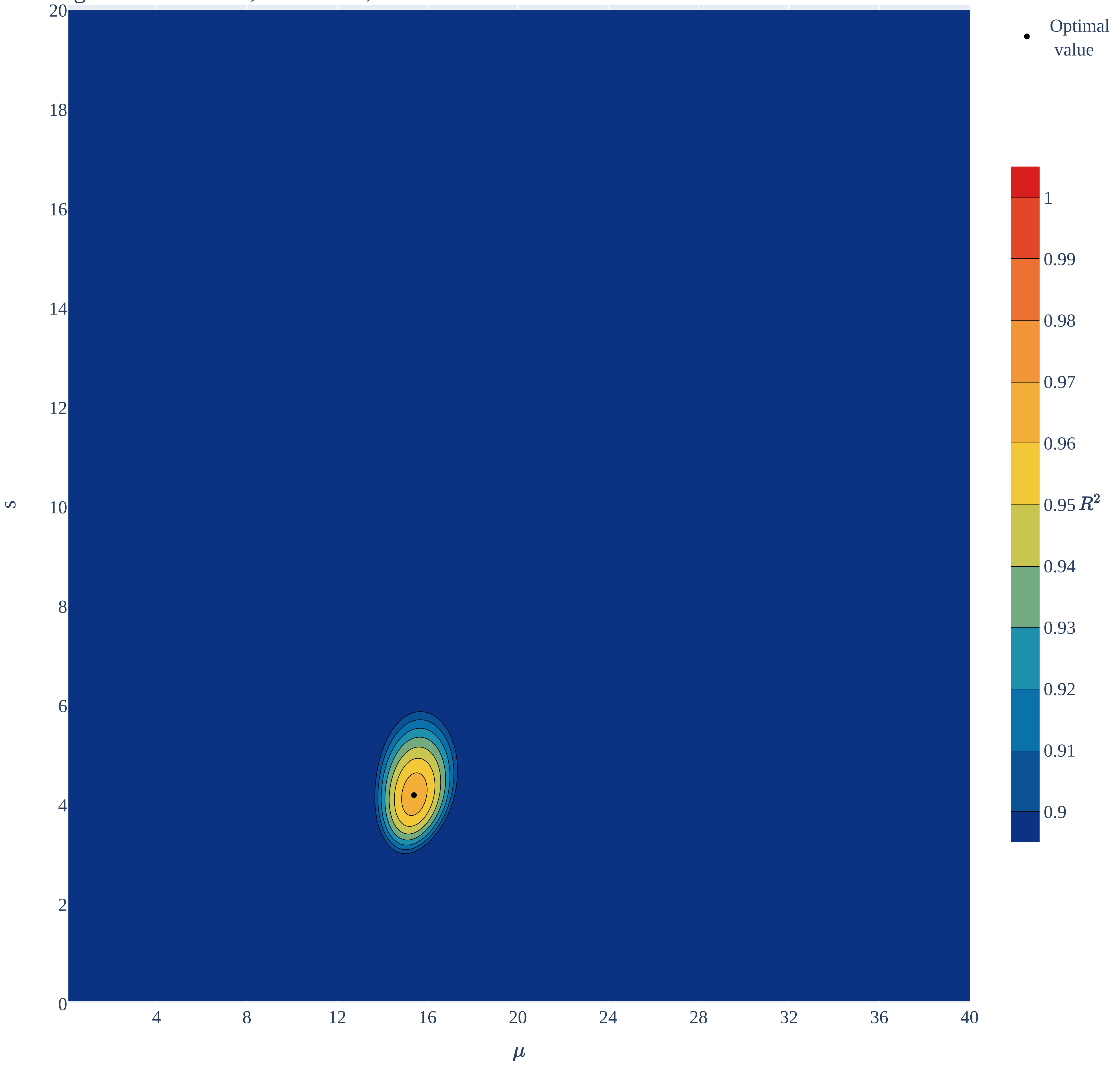

### Malignant gonadal germ cell.pdf

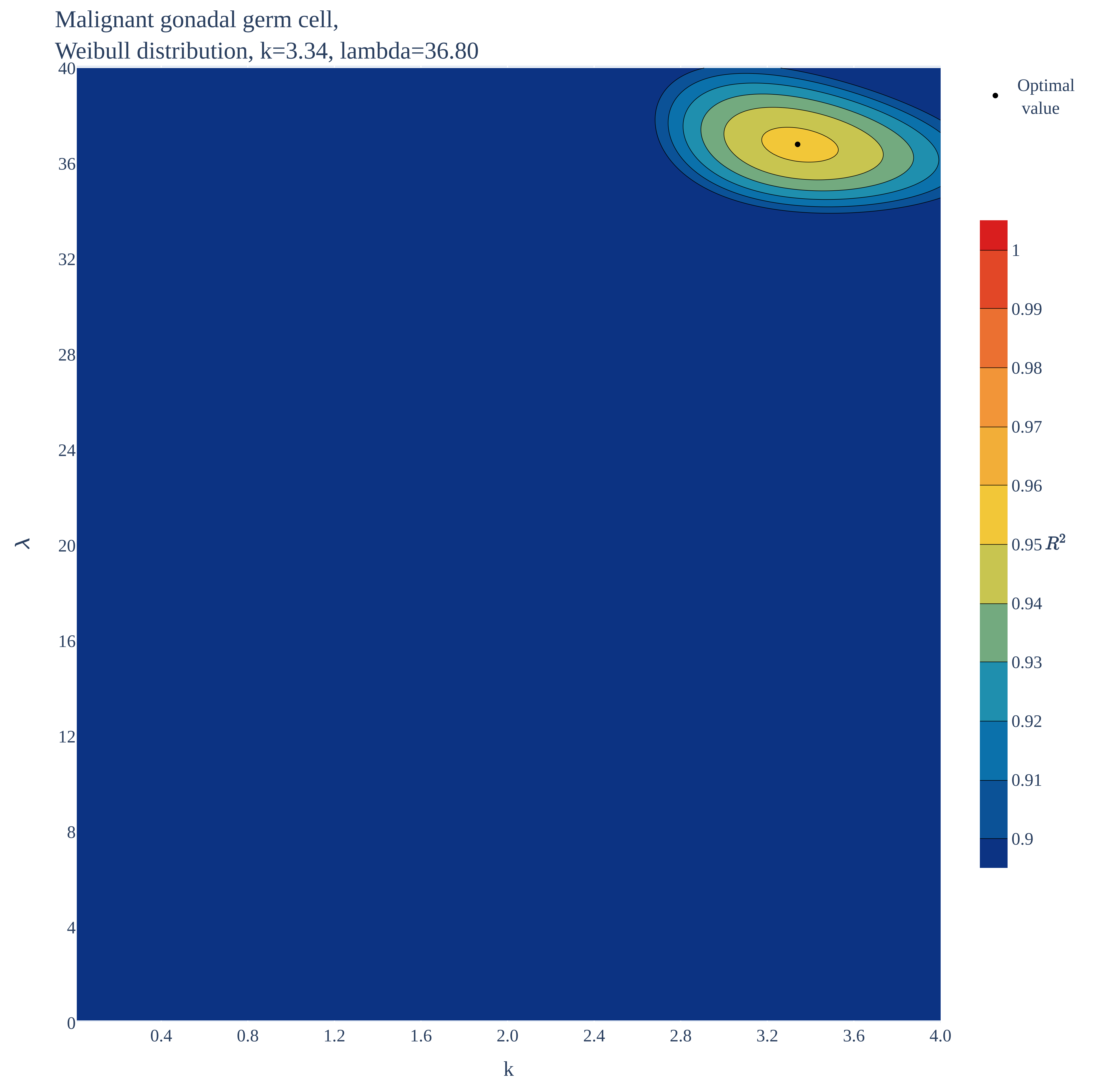

### Malignant gonadal germ cell.pdf

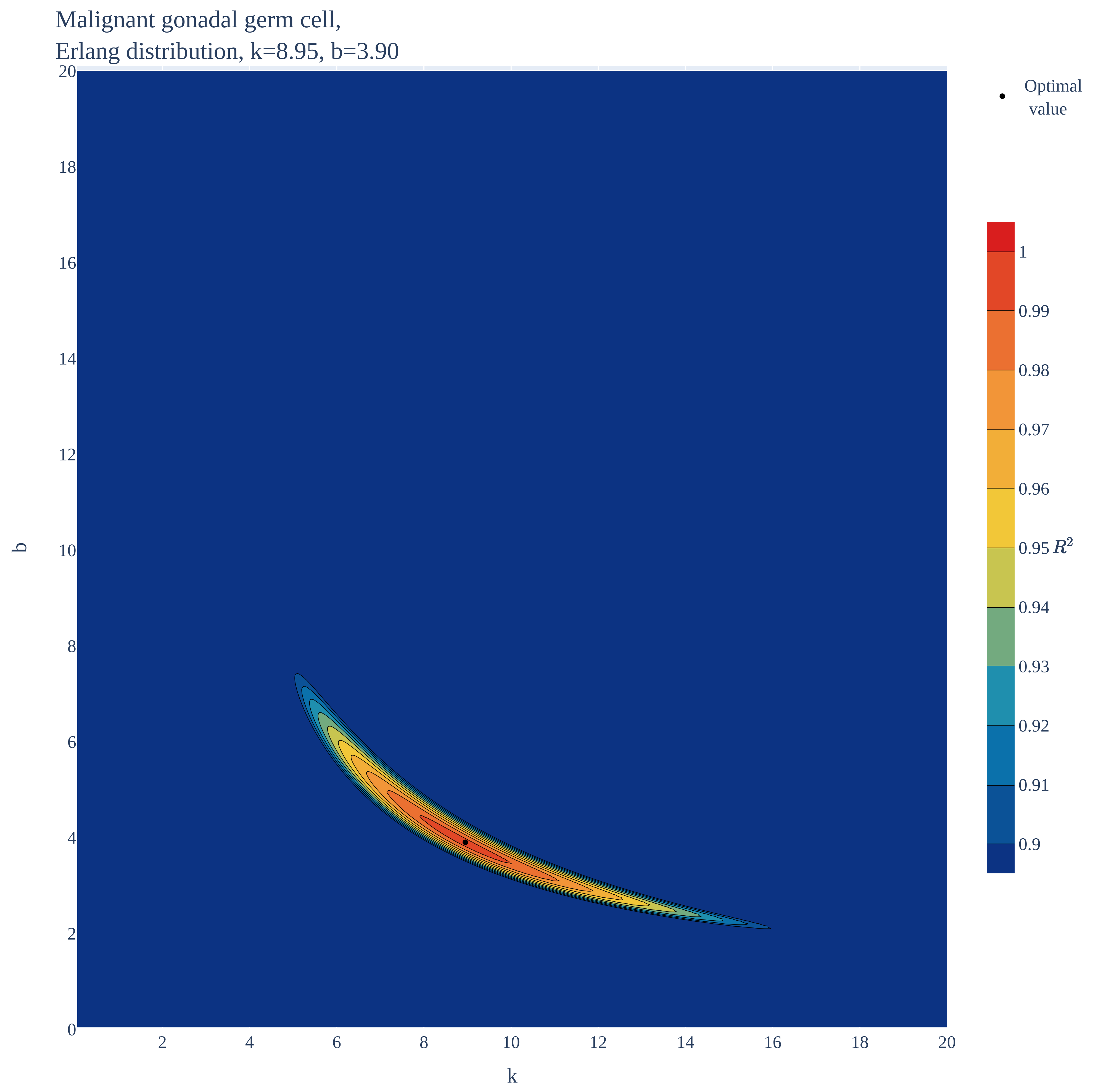

### Malignant gonadal germ cell.pdf

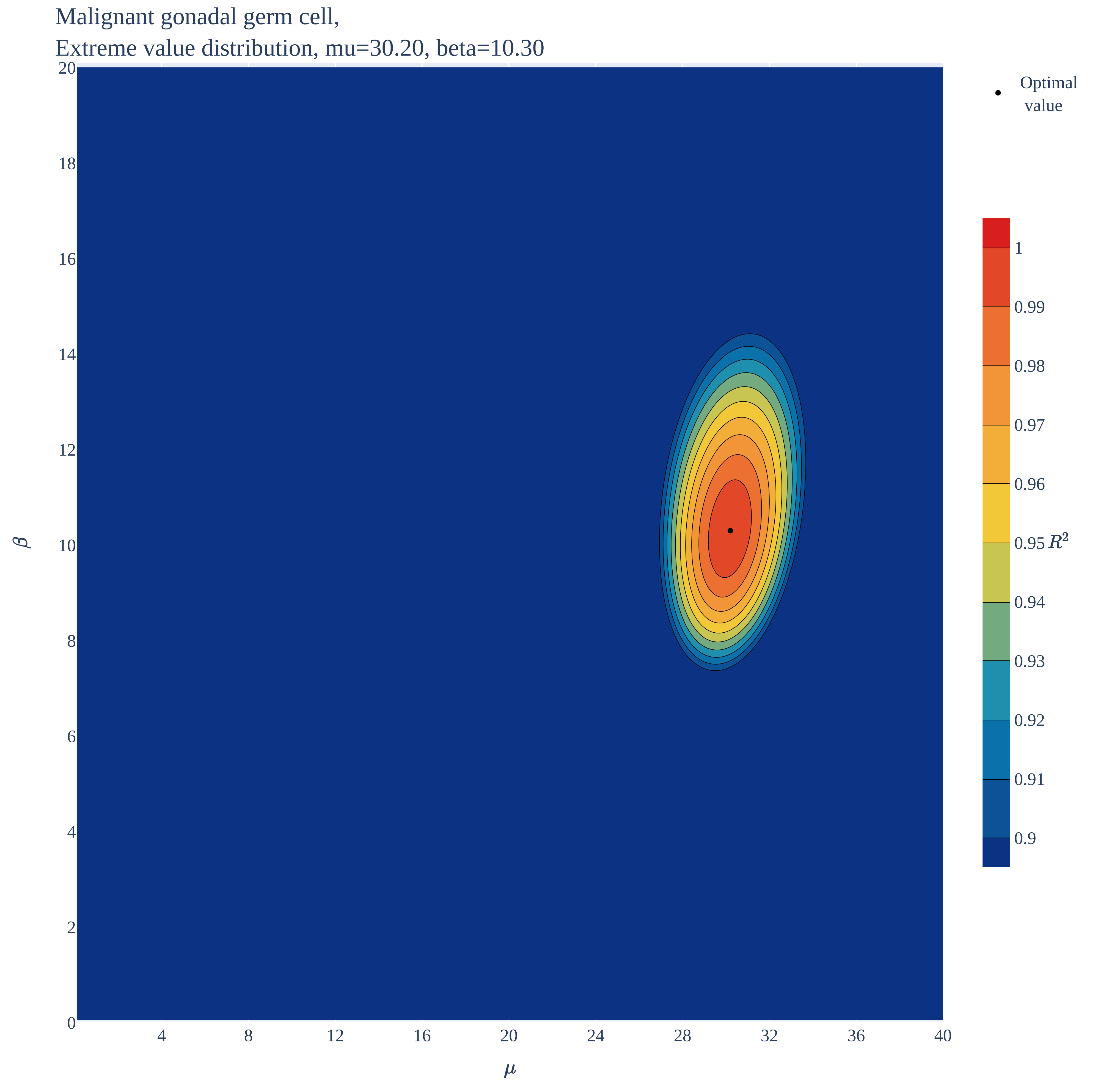

### Malignant gonadal germ cell.pdf

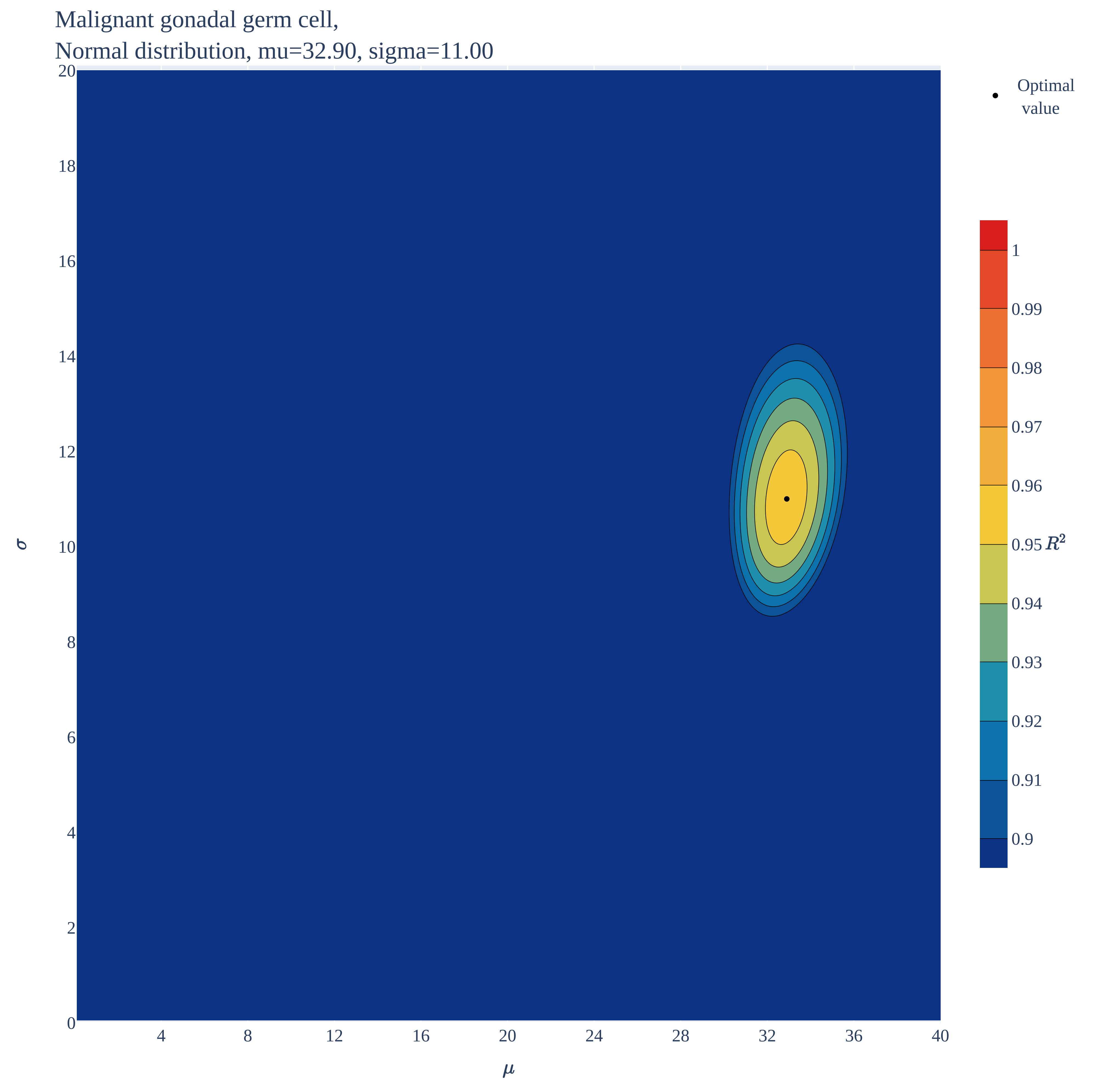

### Malignant gonadal germ cell.pdf

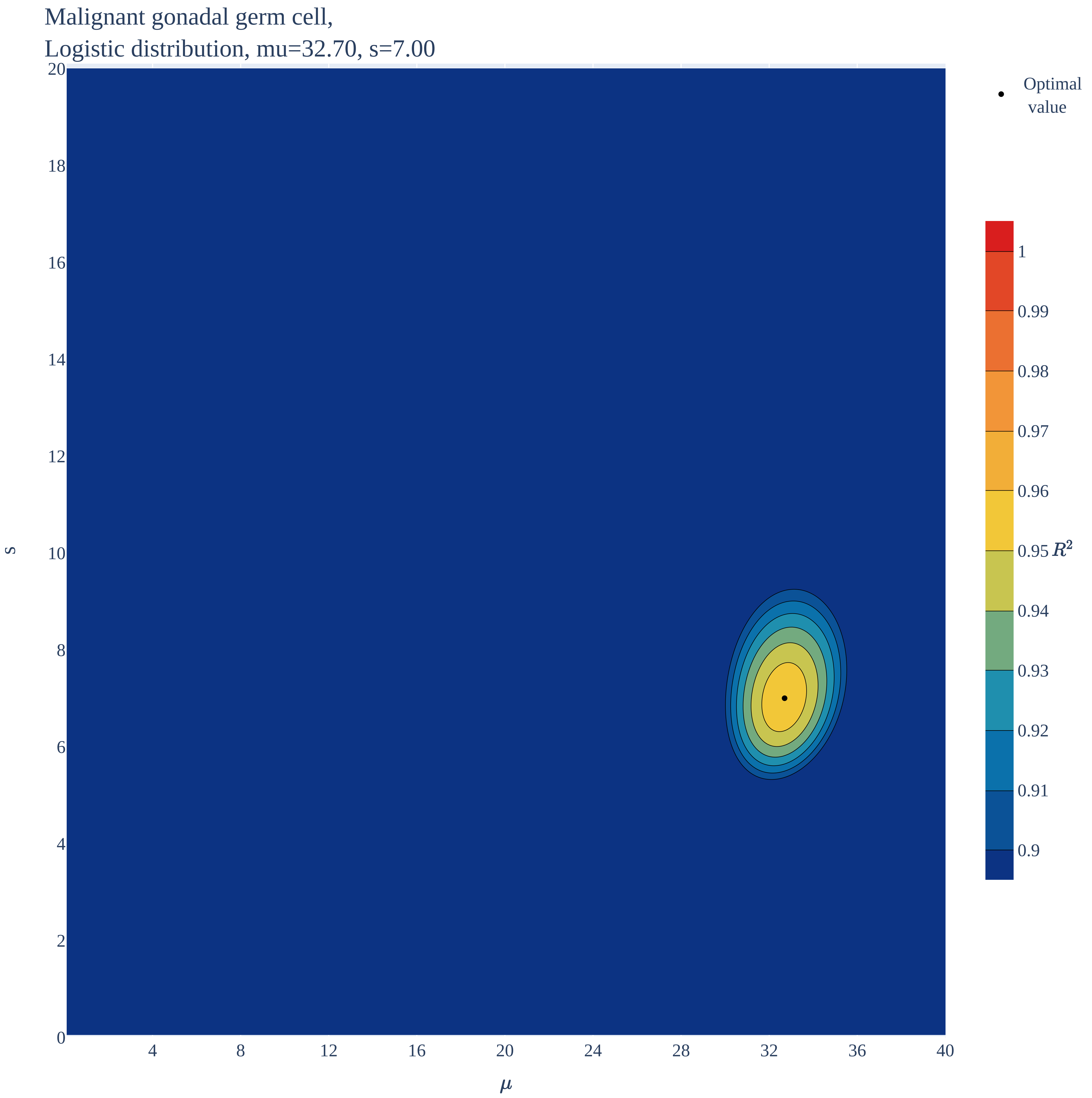

### Nephroblastoma.pdf

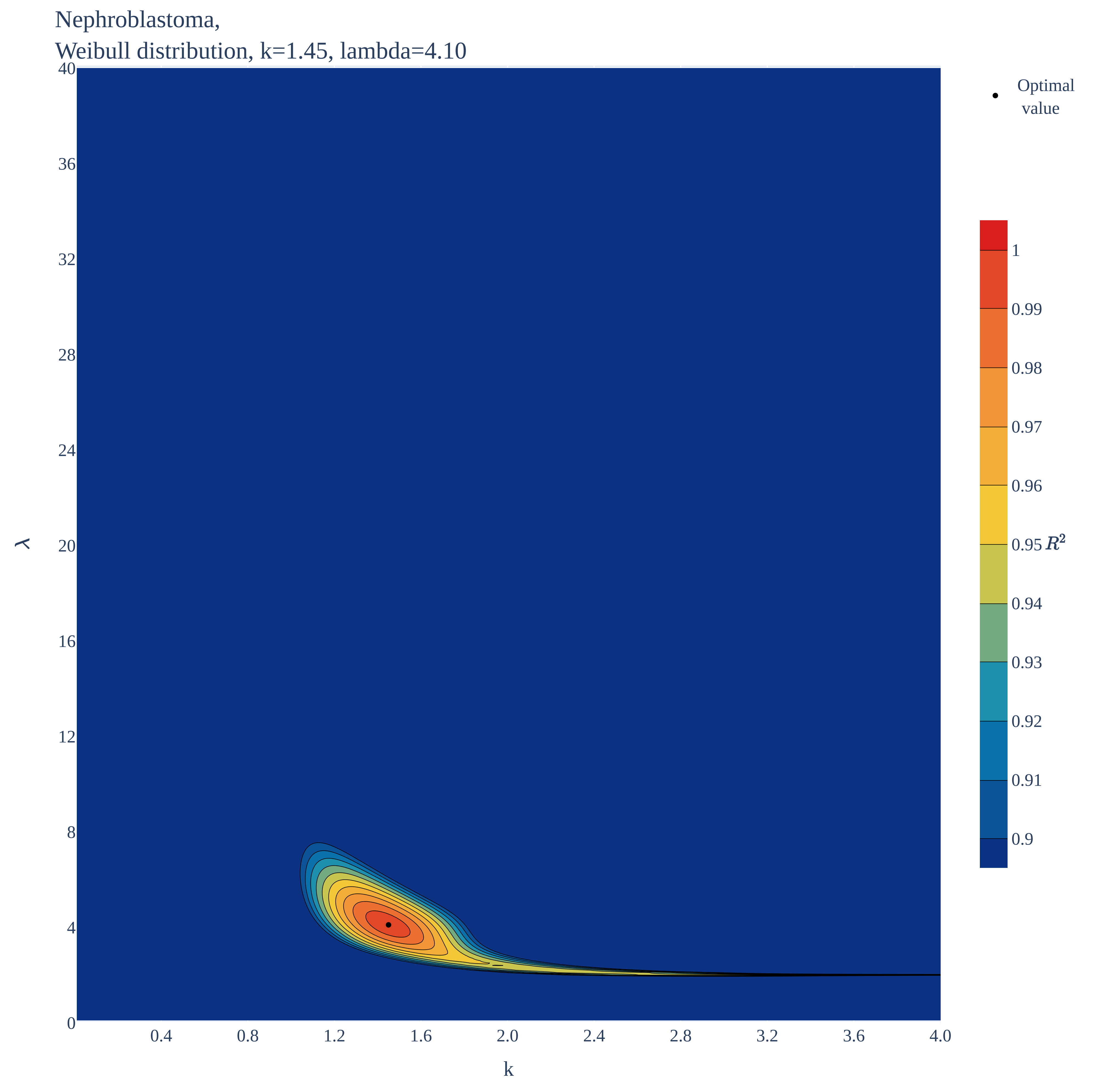

### Nephroblastoma.pdf

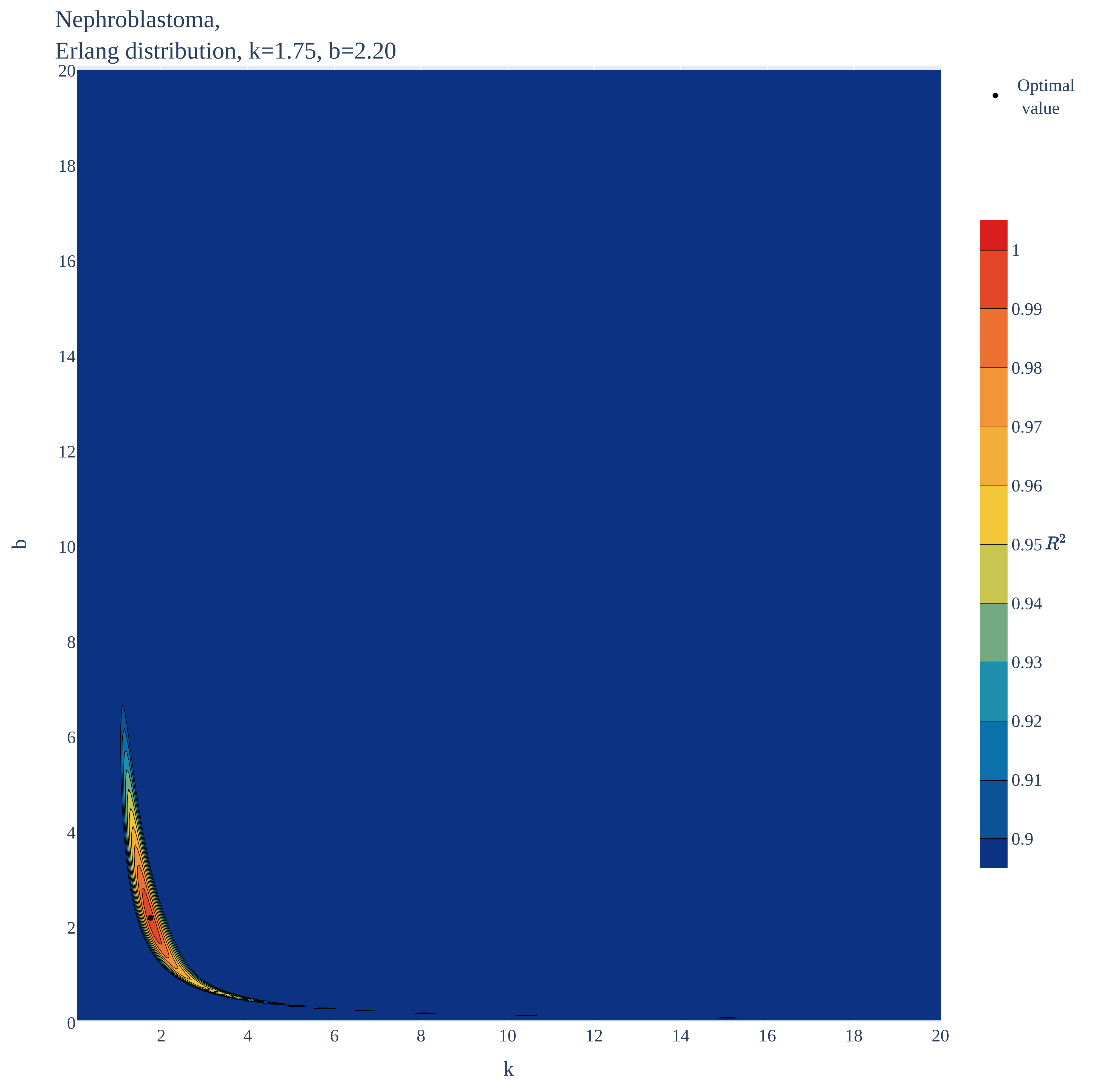

### Nephroblastoma.pdf

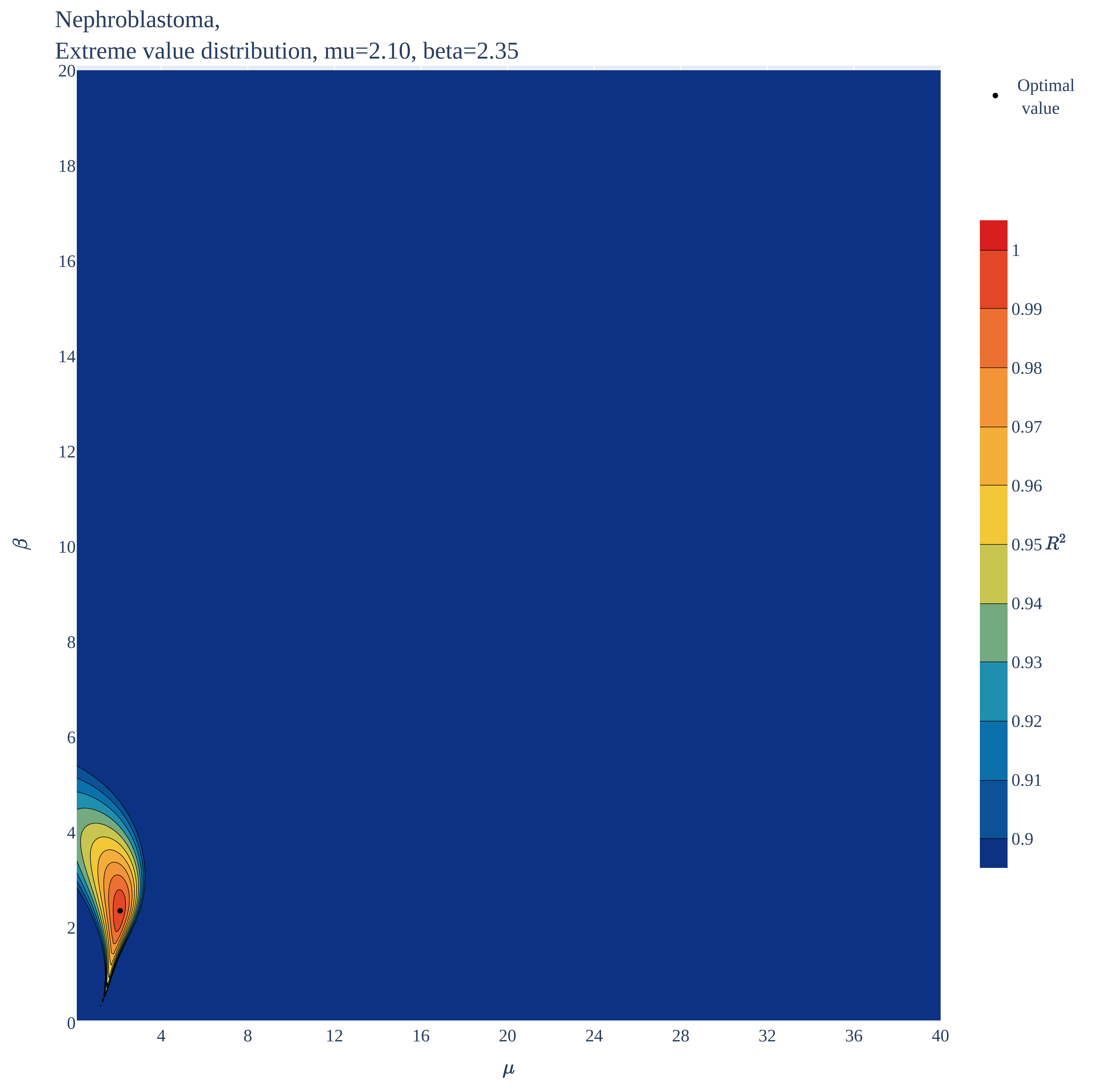

### Nephroblastoma.pdf

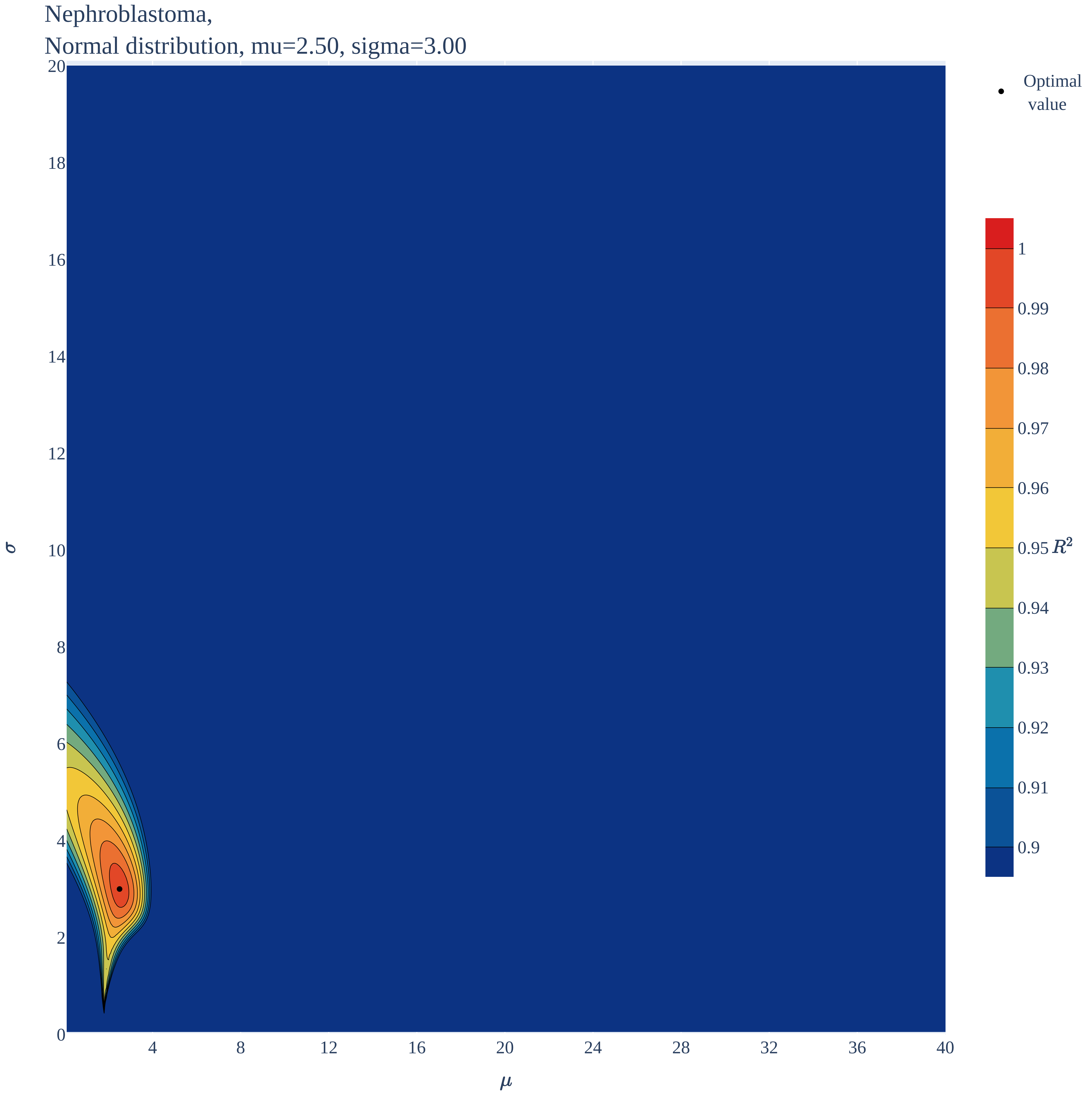

### Nephroblastoma.pdf

Nephroblastoma,
Logistic distribution, mu=2.40, s=1.95

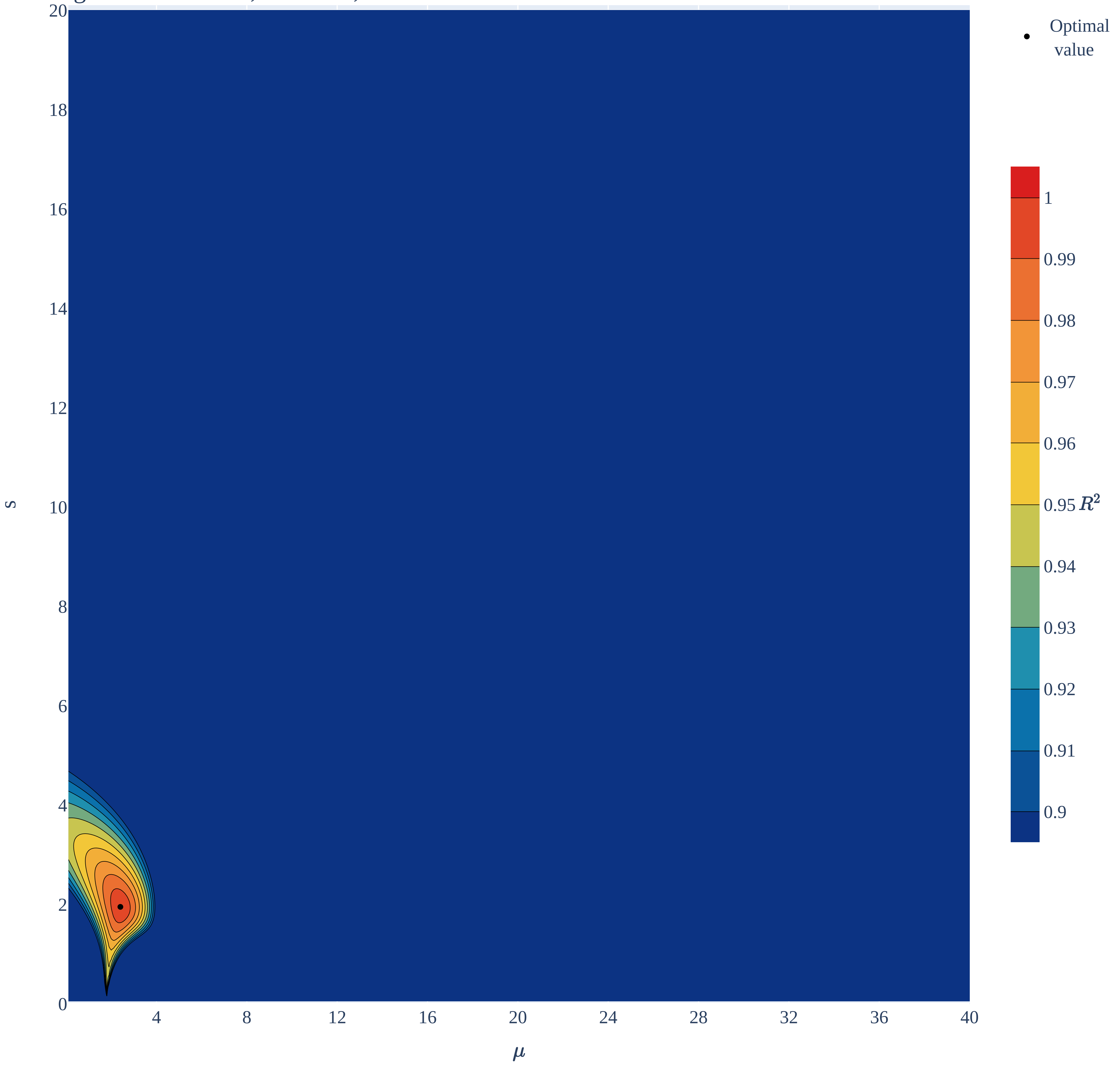

### Neuroblastoma.pdf

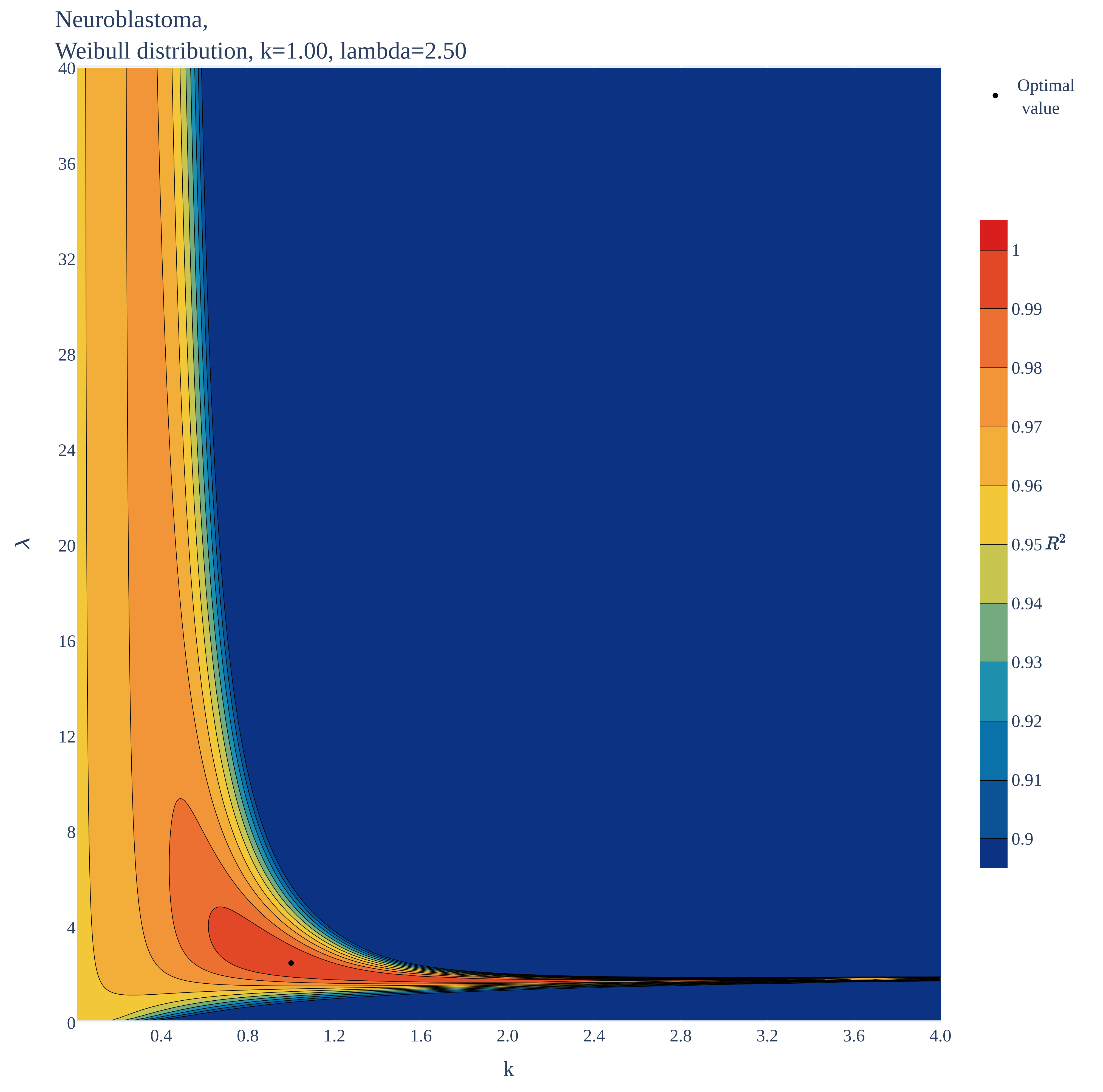

### Neuroblastoma.pdf

Neuroblastoma,
Erlang distribution, k=1.00, b=2.50

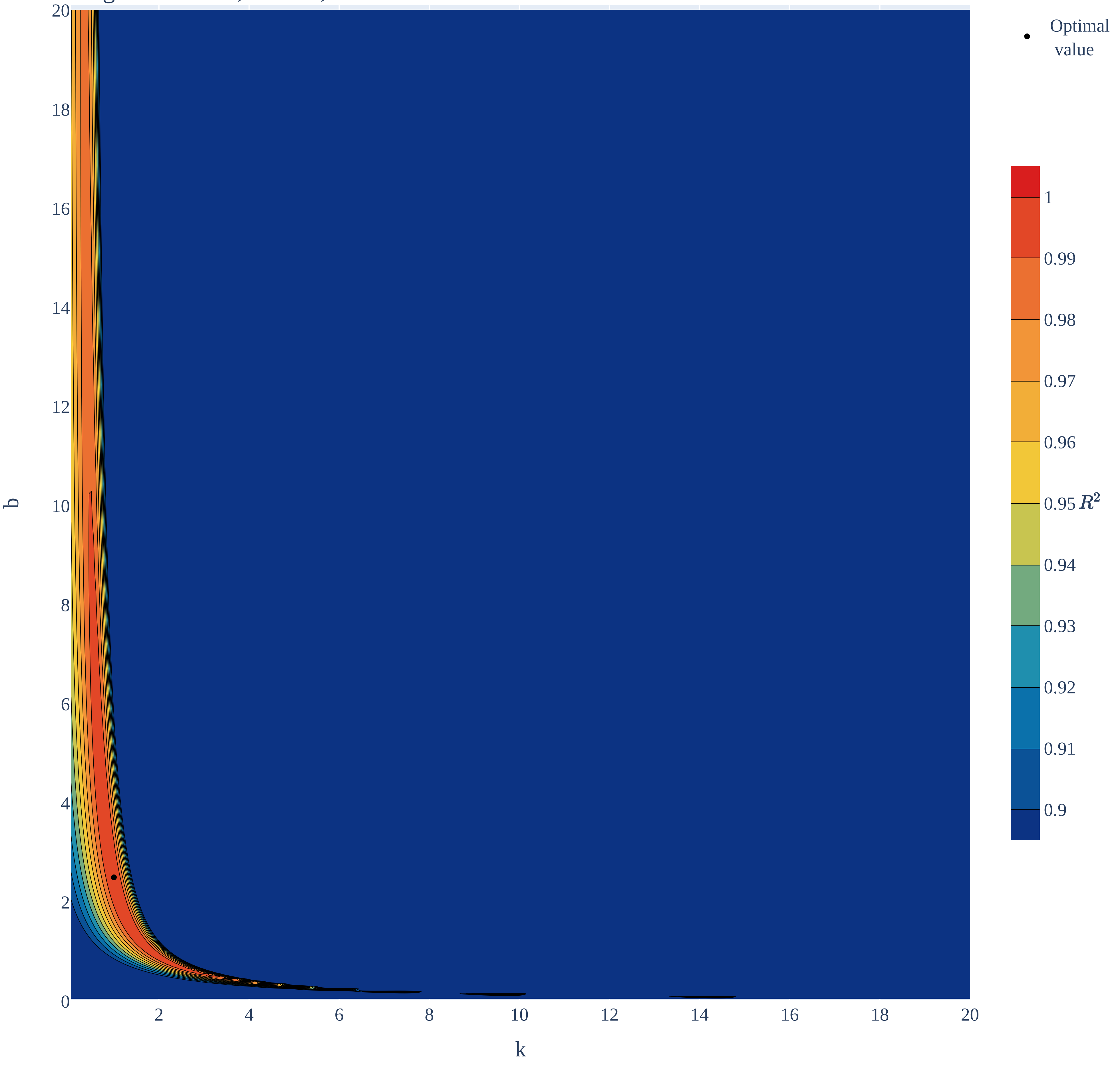

### Neuroblastoma.pdf

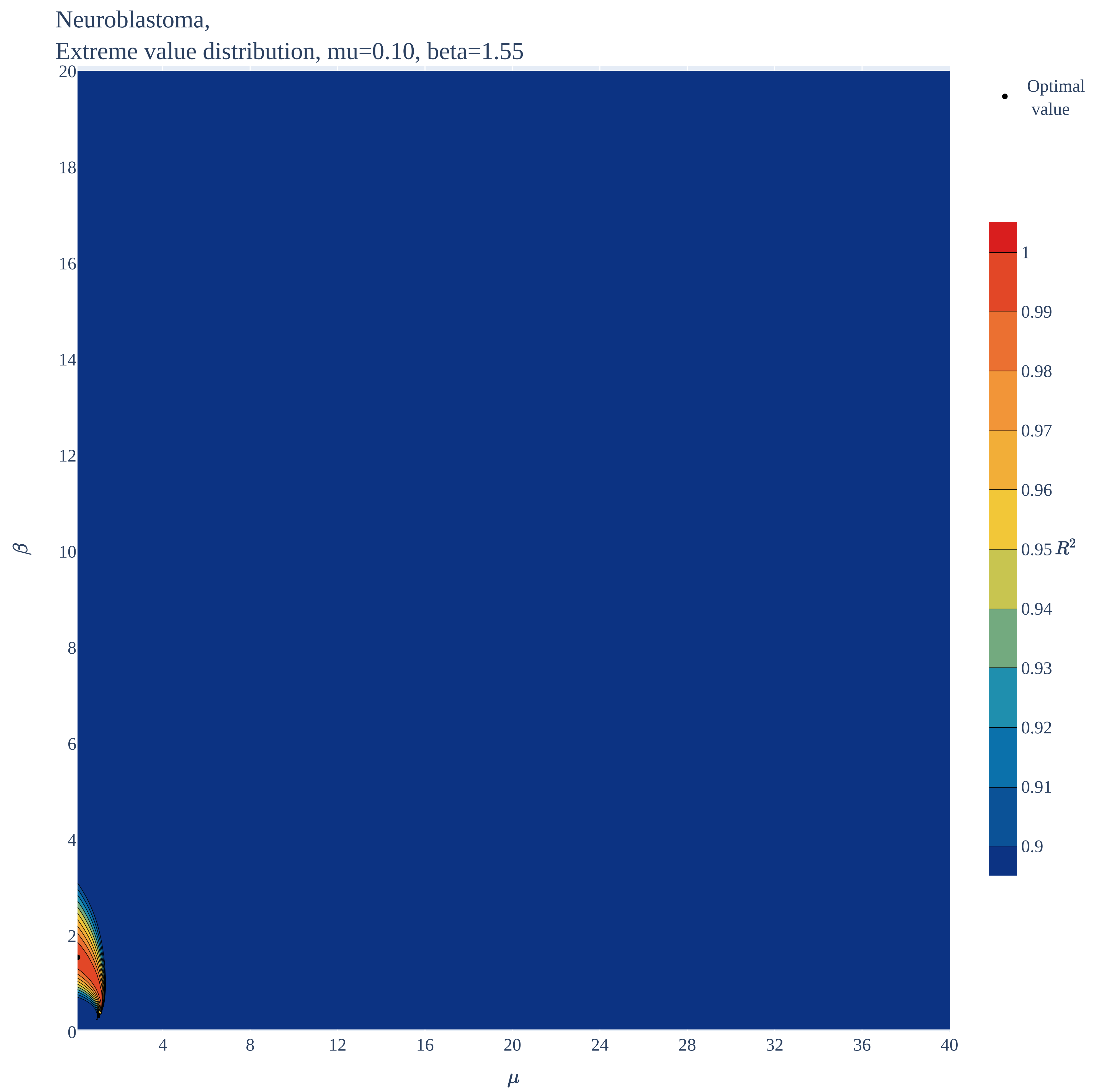

### Neuroblastoma.pdf

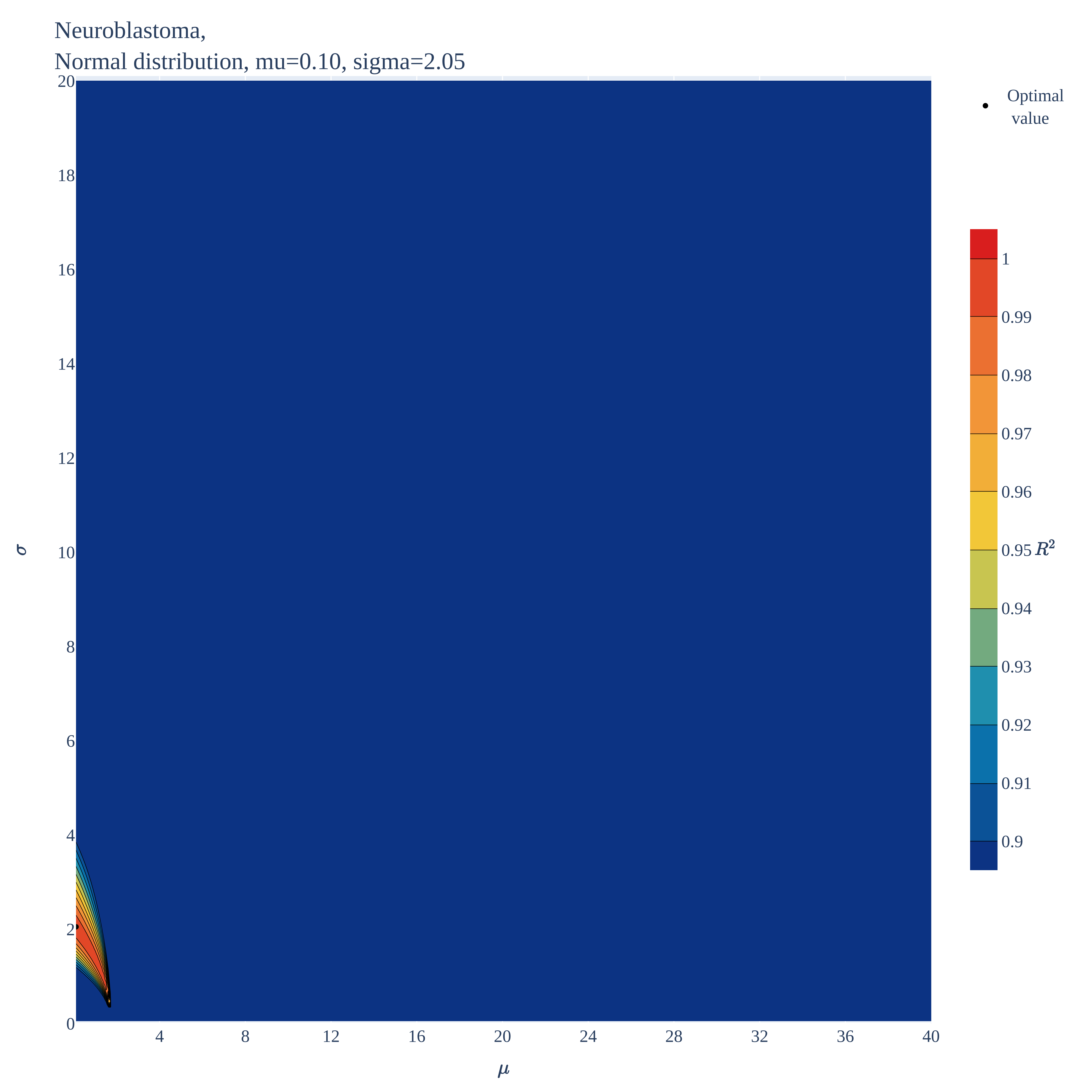

### Neuroblastoma.pdf

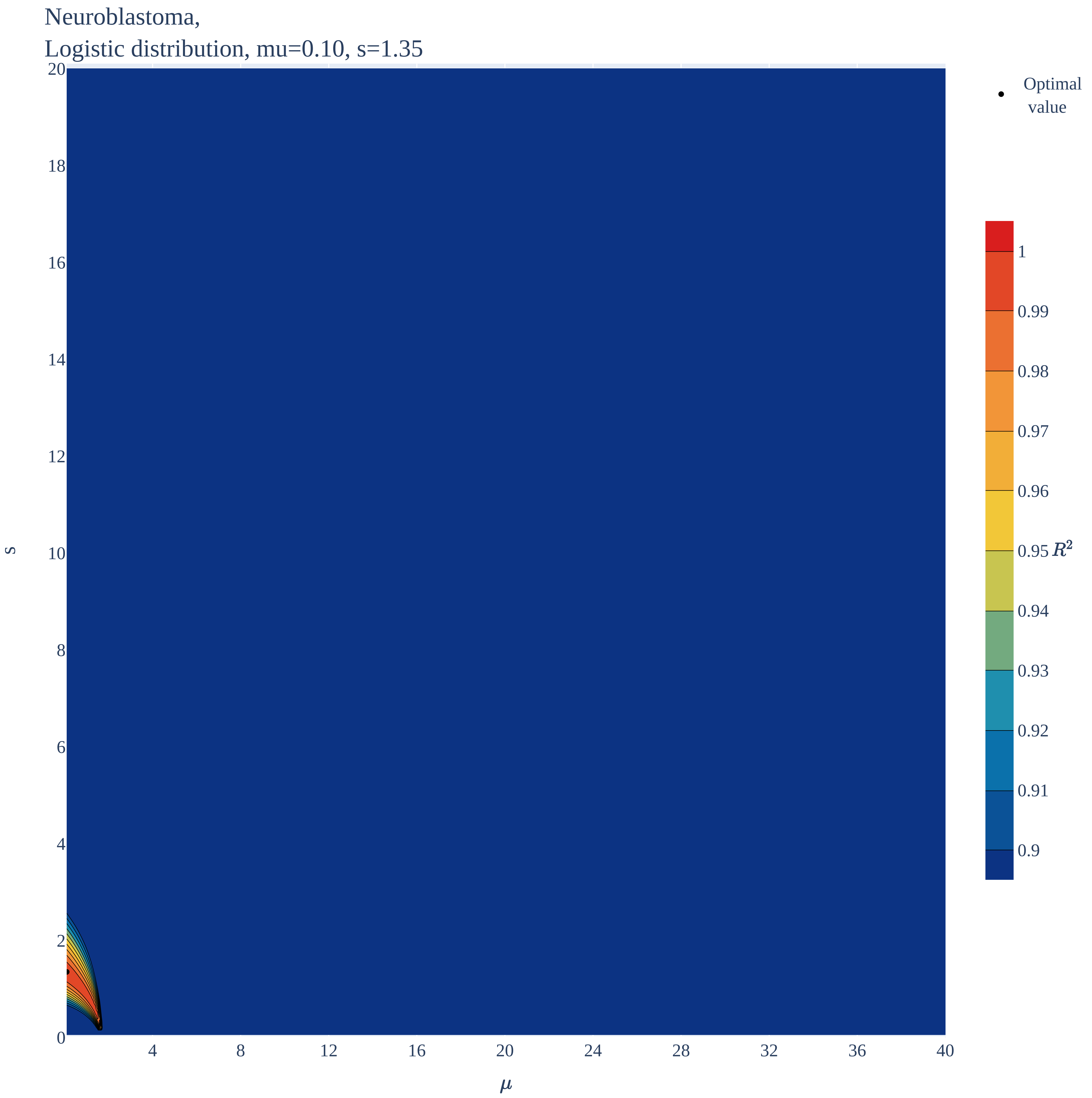

### Retinoblastoma.pdf

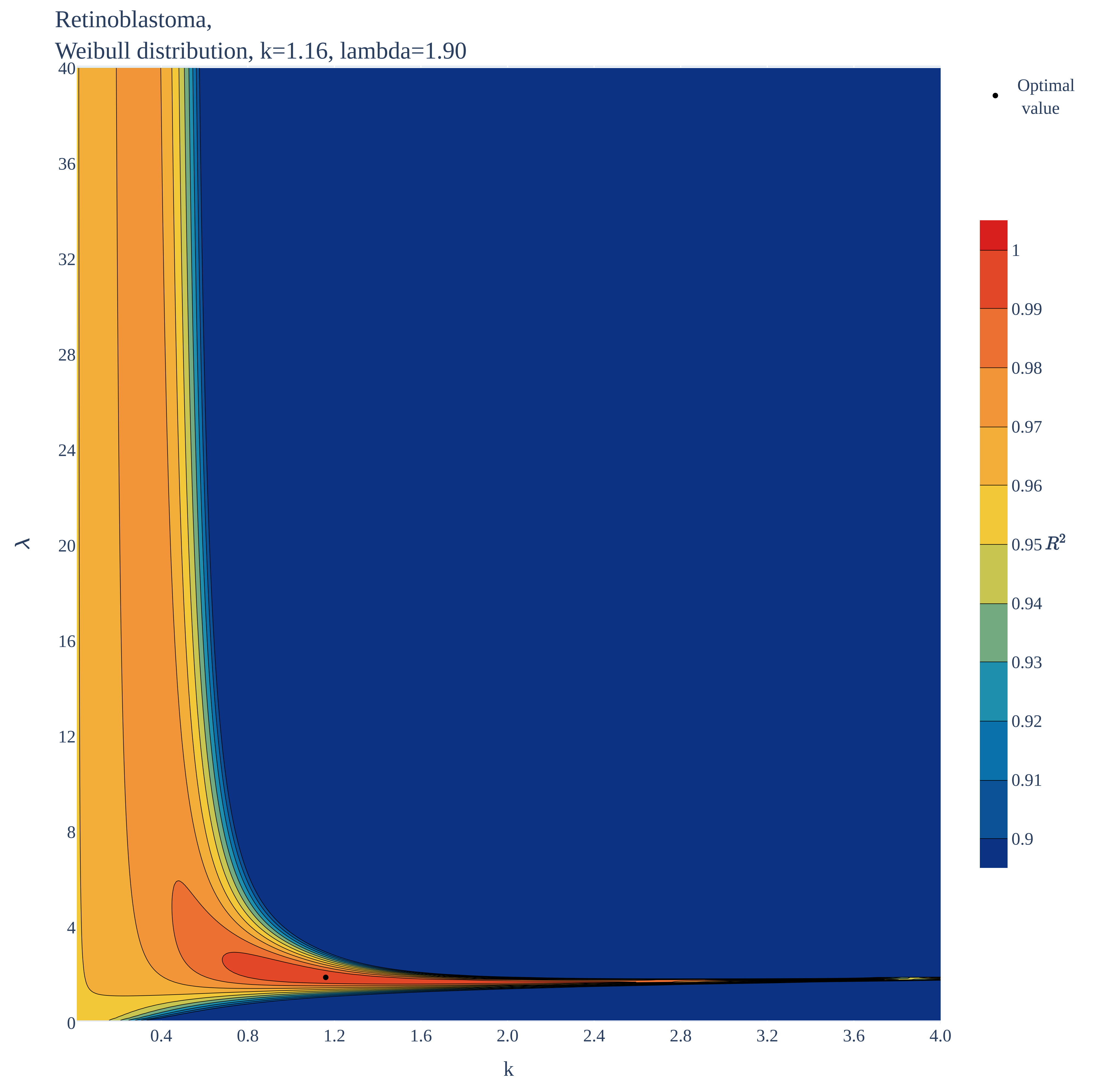

### Retinoblastoma.pdf

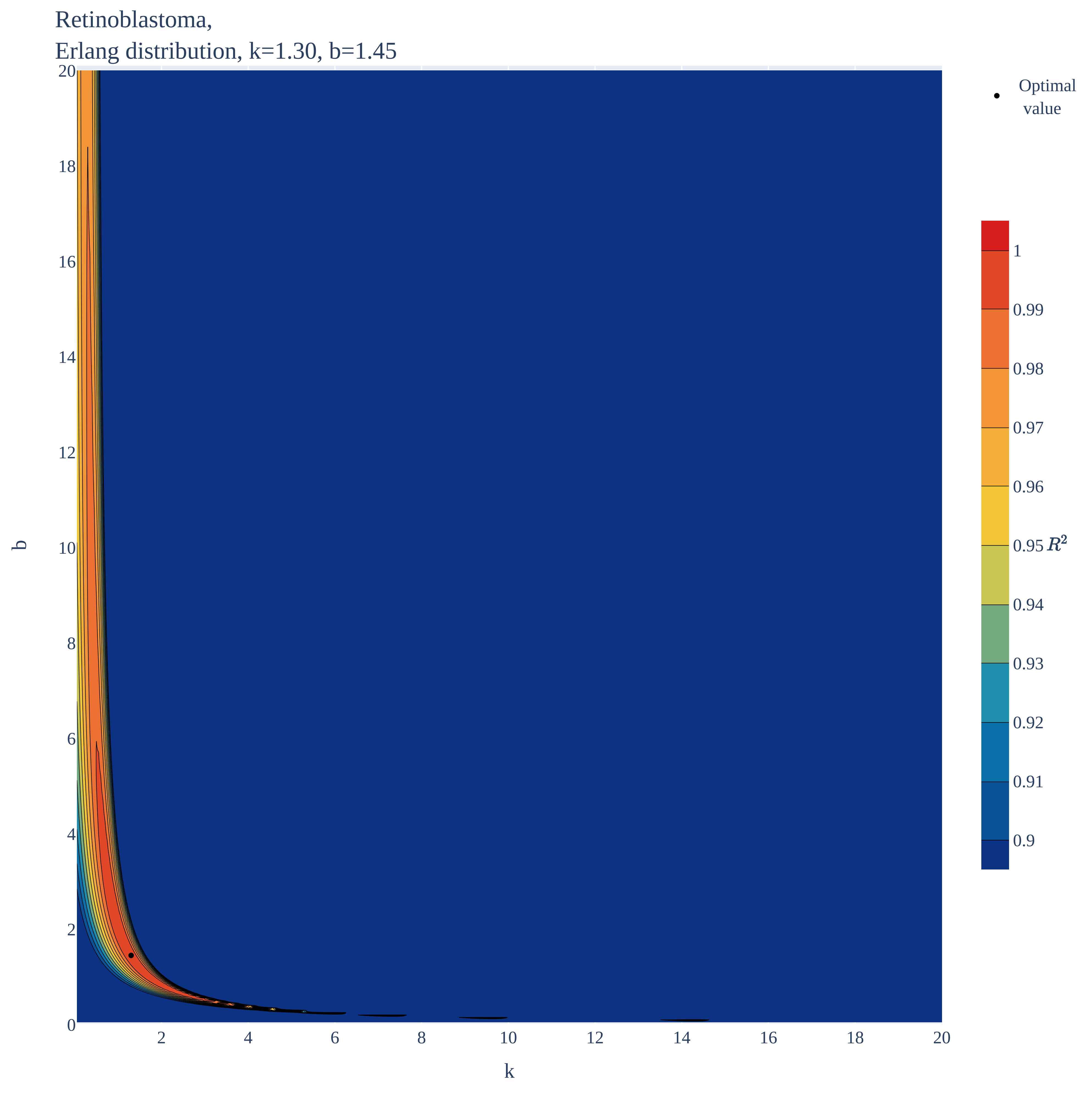

### Retinoblastoma.pdf

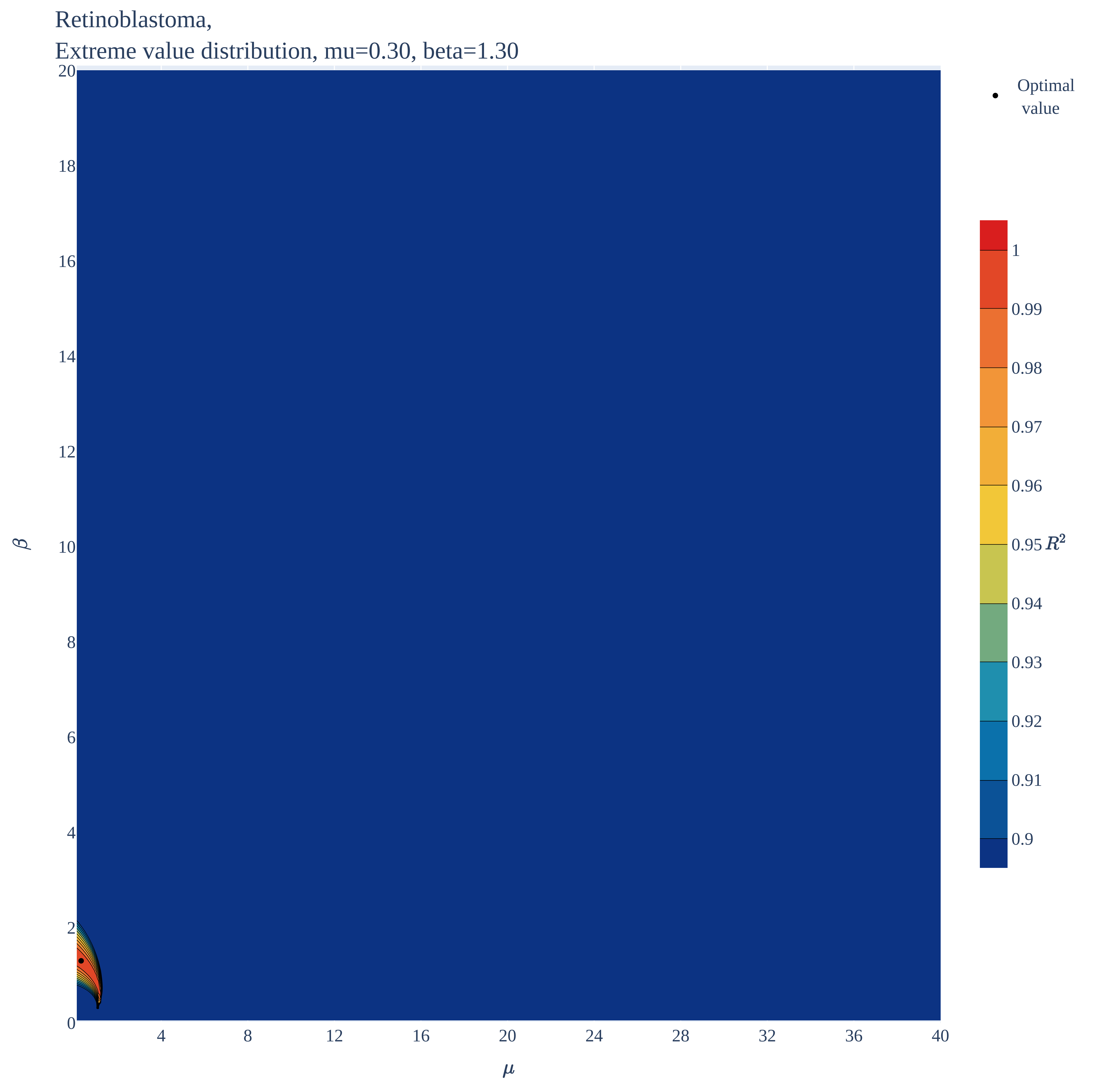

### Retinoblastoma.pdf

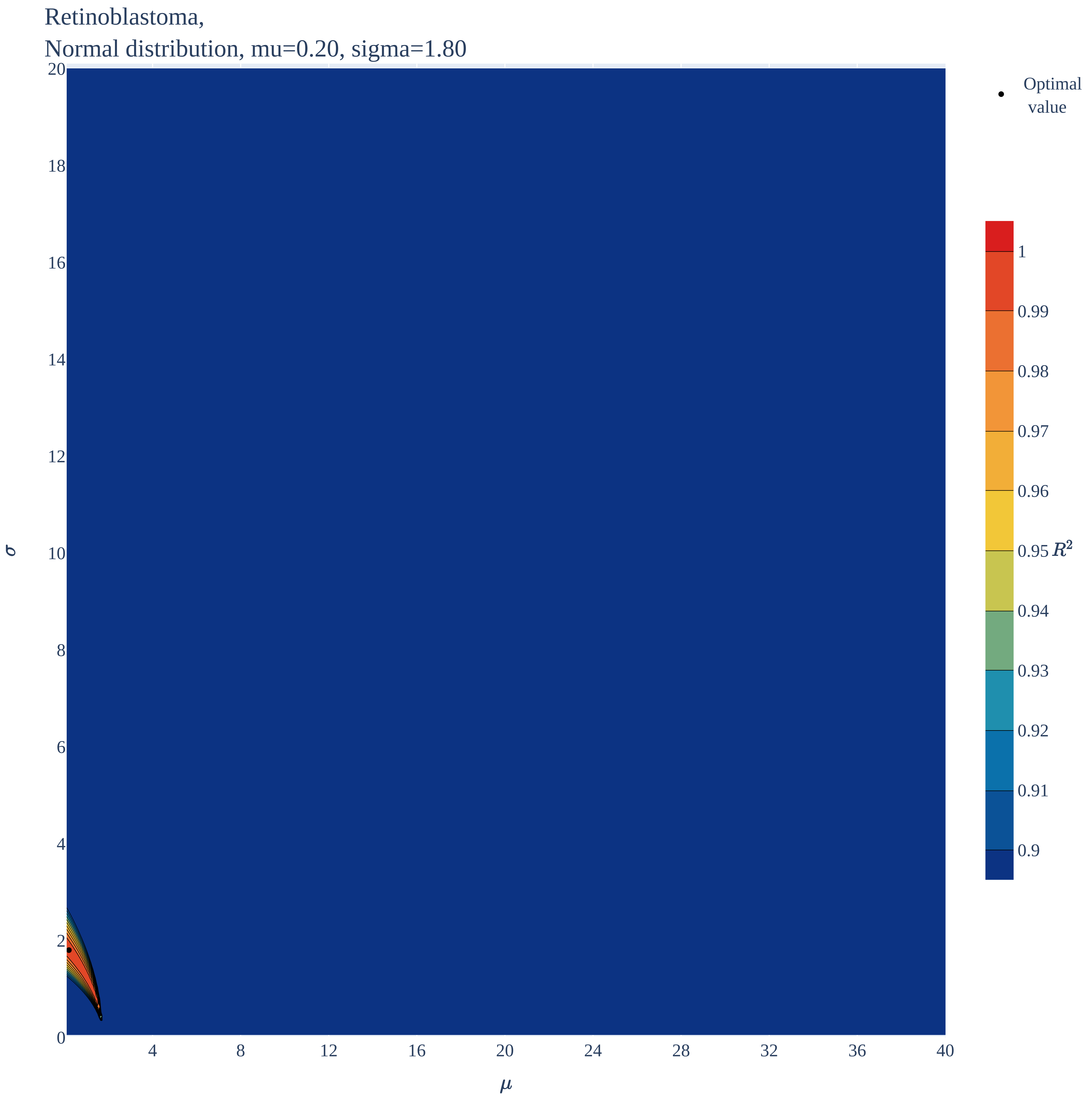

### Retinoblastoma.pdf

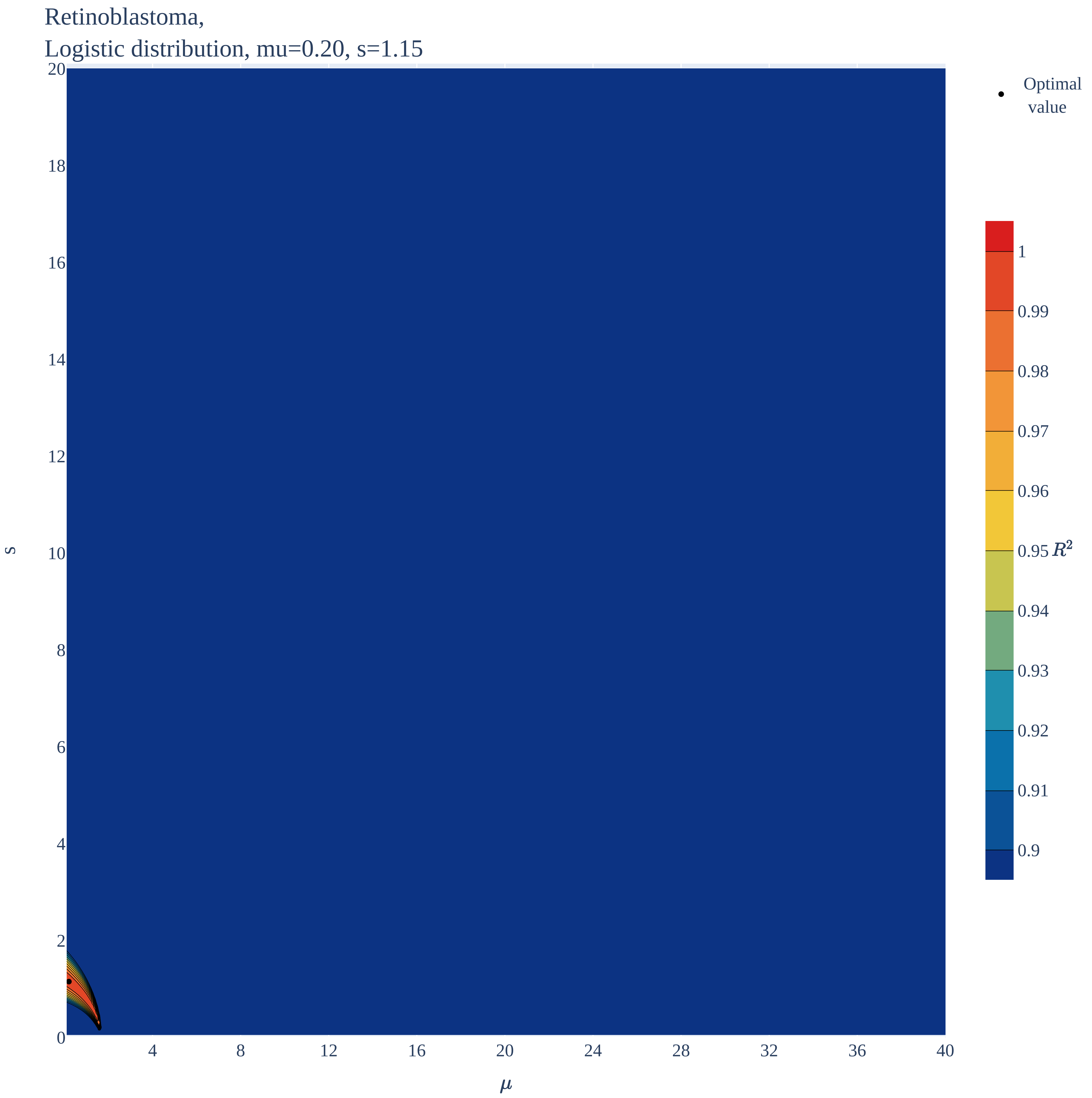
